# Relational biological structure improves fine-mapping of causal GWAS variants under weak signal

**DOI:** 10.64898/2026.05.15.725513

**Authors:** Ehsan Estaji, Shi-Wei Zhao, Zhao-Yang Chen, Shuai Nie, Jian-Feng Mao

## Abstract

Linkage disequilibrium (LD) makes causal GWAS variants indistinguishable from correlated neighbours; resolving them is the fine-mapping problem, and the challenge is species-specific: humans face dense ancestry-imbalanced LD, yeast and *Arabidopsis* exceptionally long LD, and crop germplasm sparse and fragmented annotations that defeat human-biobank curation pipelines. Bayesian fine-mappers integrate annotations as flat per-variant priors, discarding the relational structure linking variants to tissue-specific eQTLs, pathways and protein–protein interactions. Hierarchical belief propagation (HBP) on a variant–gene–pathway factor graph matches Bayesian baselines at 5–40× speed; an annotation-adaptive complement, graph-augmented fine-mapping (GAFM), wins 27–2 against SuSiE at weak signal and recovers *LDLR, APOE, LPL, GCKR* and *ANGPTL3* at single-variant resolution across four Pan-UK Biobank ancestries. On the 3,000 Rice Genomes grain weight + shape panel, mixture-prior posterior reweightings of GAFM/HBP and their ensemble (GAFM-MX, HBP-MX, ENS) reach 47.6% top-1-PIP exact-position recovery of 21 panel-matched stable QTNs — the highest of any method, exceeding SuSiE (28.6%) and SBayesRC (14.3%) — at 200–700× SuSiE’s per-locus speed. Across 692 leads in four species, a non-uniform per-variant prior, not uniform high coverage, lets the graph break LD ties: adding a regulatory-element flag to an otherwise uniform human cache flips HBP narrower than GAFM from 0% to 88% on 321 Pan-UKB leads. These results recast multi-omics fine-mapping as a non-uniform-prior-curation problem rather than a uniform-coverage problem, and reframe post-GWAS analysis as message passing over biological structure rather than weighted regression on flattened annotations.

## Introduction

Genome-wide association studies have identified tens of thousands of trait-associated loci[1, 2], but resolving the causal variants that drive each signal remains the rate-limiting step between statistical discovery and mechanistic biology. Because nearby variants co-segregate through linkage disequilibrium (LD), a single causal variant is typically indistinguishable from hundreds of non-causal proxies on statistical evidence alone. Fine-mapping procedures aim to break this degeneracy[3]; their accuracy directly constrains downstream experimental follow-up and the trans-ancestry portability of polygenic scores.

The challenge has different shapes across the regimes in which fine-mapping is most needed. In human medical genetics, biobank-scale sample sizes deliver high signal-to-noise but LD is dense and ancestry-imbalanced, so the same lead variant can map to tightly correlated proxy clusters of 10^2^–10^3^ variants in non-European panels even when the European credible set resolves to a singleton. In the model organisms used to interrogate trait architecture under controlled conditions — inbred yeast and *Arabidopsis* 1001G — LD is exceptionally long, so single-locus credible sets routinely exceed 10^3^ variants and statistical resolution alone cannot isolate causal variants. In agricultural germplasm such as the 3,000 Rice Genomes panel[4], accessions span between-population *F*_*ST*_ ≈ 0.5, most traits have only ordinal phenotypic scoring, and curated causal-gene catalogues exist for only a few well-studied phenotypes[5]. A method that improves fine-mapping must therefore travel between these regimes without re-engineering for each.

Modern fine-mapping is dominated by Bayesian variable-selection methods operating on a per-locus genotype matrix or summary-statistics vector— SuSiE[6], FINEMAP[7], PolyFun[8], PAINTOR[9]—with two recent extensions that add an infinitesimal random-effect background (SuSiE-inf and FINEMAP-inf)[10] and a genome-wide multi-component Bayesian mixture (SBayesRC)[11] that Wu *et al*. 2026[12] used as the substrate for a genome-wide fine-mapping framework. All share a common design choice: biological annotations, where used at all, enter as *flat per-variant scalars*—averaged across tissues, collapsed across pathways, and stripped of their relational structure. A variant’s connection to a particular gene in a particular tissue, participating in a particular pathway coupled via protein–protein interactions to further functional neighbourhoods, is flattened to a single weight. Constructing that scalar is itself a curation step that scales poorly to non-model species — including crops, livestock and model organisms — where functional annotations are sparse, fragmented and lack a pre-harmonised category catalogue.

This flattening is the limiting approximation. Multi-omics resources released since 2020—the Genotype-Tissue Expression (GTEx) project v8 with 43 million tissue-specific expression quantitative trait loci (eQTLs)[13], STRING v12 with 230,000 high-confidence protein–protein interactions (PPIs) at combined score ≥ 700[14], and the ENCODE Project’s candidate cis-regulatory elements (cCREs) with 370,000 entries[15]—carry exactly the relational biology that flat priors discard. When signal is strong, statistical evidence alone resolves the causal variant and relational context is optional; but this is not the regime that matters most for novel discovery. Subgenome-wide-significant loci, low-frequency effects and non-European ancestries all sit in the weak-signal regime, where annotation priors dominate the posterior. The question we address is whether fine-mapping methods that represent multi-omics context *as a graph, not as a vector* can decisively outperform flat-prior methods in that regime.

Answering that question requires a computational substrate in which multi-omics structure is *primary data*, not reconstructed on demand from files. Graph databases provide exactly this[16]: typed nodes (variant, gene, pathway, tissue, regulatory element) and labelled edges (consequence, eQTL, pathway membership, protein–protein interaction) are first-class storage objects; index-free adjacency makes traversing a relationship an *O*(1) operation regardless of database size; and new annotation layers register as additional edge types without restructuring the underlying schema. These properties are absent from the file-based pipelines assembled from variant call format (VCF) files, browser extensible data (BED) tracks, general feature format (GTF) annotations, and pathway tables glued together by `bedtools`/`tabix` lookups. Encoding the biology directly as a graph reframes fine-mapping as a Bayesian computation *on the graph itself* —not as a regression that happens to have annotation weights attached—and collapses what would otherwise be a multi-file data-integration task into a single database traversal that can be iterated thousands of times per second.

Doing so changes the fine-mapping accuracy–speed frontier. GraphGWAS is a fine-mapping platform built on a graph database[16] into which users import their own genotype, summary-statistics and multi-omics data; the schema and algorithms are database-agnostic and the present reference implementation runs on Neo4j 5.26 Community Edition. The demonstration loadout used in this paper covers 70.7 M variants from the 1000 Genomes Phase 3 high-coverage release, 20,092 GENCODE v47 genes[17], 43.2 M GTEx v8 eQTL edges, 230,850 STRING interactions and 370,000 ENCODE regulatory elements. Its central methods — hierarchical belief propagation (HBP), with proved geometric convergence via a Banach contraction argument, and graph-augmented fine-mapping (GAFM), with a proof that the causal variant ranks first under mild LD-decay assumptions — carry the full variant → gene → pathway factor graph with PPI coupling through every update. On 90 simulated chromosome 22 loci, HBP matches SuSiE and FINEMAP at 5–40× the speed; on 79 weak-signal tissue-specific eQTL simulations, GAFM beats SuSiE 27–2 head-to-head (sign-test *p <* 10^−5^). Posterior inclusion probabilities (PIPs) are calibrated with 0% null false-positive rate (FPR) across 100 simulations. A sumstats-only entry path consumes Pan-UK Biobank[18] directly, fine-mapping across four ancestries (EUR *N* =420,531; CSA, AFR, EAS) and recovering canonical lipid genes *LDLR, APOE, LPL, GCKR, ANGPTL3* at single-variant resolution. The same code fine-maps yeast[19], *Arabidopsis*[20] and the 3,000 Rice Genomes panel without algorithmic change. Across 692 leads in the four species, a direct intervention experiment shows that the operative quantity is per-variant heterogeneity of the prior, not annotation density — adding even a coarse regulatory-element flag to the otherwise dense uniform human cache flips HBP narrower than flat-prior GAFM from 0% to 88% of leads.

## Results

### Relational priors carry structure that flat priors discard

Multi-omics resources released since 2020 — tissue-specific eQTL catalogues, protein–protein interaction networks at curated confidence, base-pair-resolved regulatory-element atlases — encode far more relational structure than flat-prior fine-mappers currently exploit. **Flat-prior fine-mapping collapses that relational structure to a scalar**. In the SuSiE/FINEMAP/PolyFun family, biological annotations enter as a per-variant weight vector *w* ∈ ℝ^*n*^, averaged across tissues and detached from the gene, pathway and interaction context each variant belongs to. Constructing that scalar weight is itself a substantial curation step: it requires harmonising heterogeneous functional sources — eQTL tables, regulatory-element atlases, pathway curations and PPI networks — into a single per-variant weight, typically via stratified LD-score regression over a pre-curated functional-category catalogue (the human Baseline 2.2 catalogue uses 96 such categories[21]). The burden is heaviest precisely where post-GWAS analysis is most needed: in non-model species, including crops and livestock, where functional annotations are sparse, fragmented and lack a pre-harmonised catalogue. The graph representation never demands such a scalar — annotation layers register as typed edges where data exists, missing edges stay missing, and end users get a ready-to-go pipeline without species-specific harmonisation (see Discussion “Discussion” for the full argument). GraphGWAS instead represents this context as a typed graph (Figure 1): variants connect to genes through `HAS_CONSEQUENCE` and tissue-specific `eQTL` edges, genes participate in pathways via `IN_PATHWAY`, and the gene layer is coupled within itself by protein–protein INTERACTS_WITH edges. The graph loaded on the 1000 Genomes Phase 3 high-coverage cohort [22, 23] carries 70.7 million variants, 20,092 GENCODE v47 genes[17], 43.2 million GTEx v8 tissue eQTLs[13], 230,850 STRING interactions at combined score ≥ 700[14], and 370,000 ENCODE cCRE regulatory elements[15] (full counts in Supplementary Table S1). All relational edges are first-class: a fine-mapping procedure can either collapse them to a scalar prior (the flat-prior approach) or carry them through the inference (the relational approach we develop below).

**Fig. 1.**
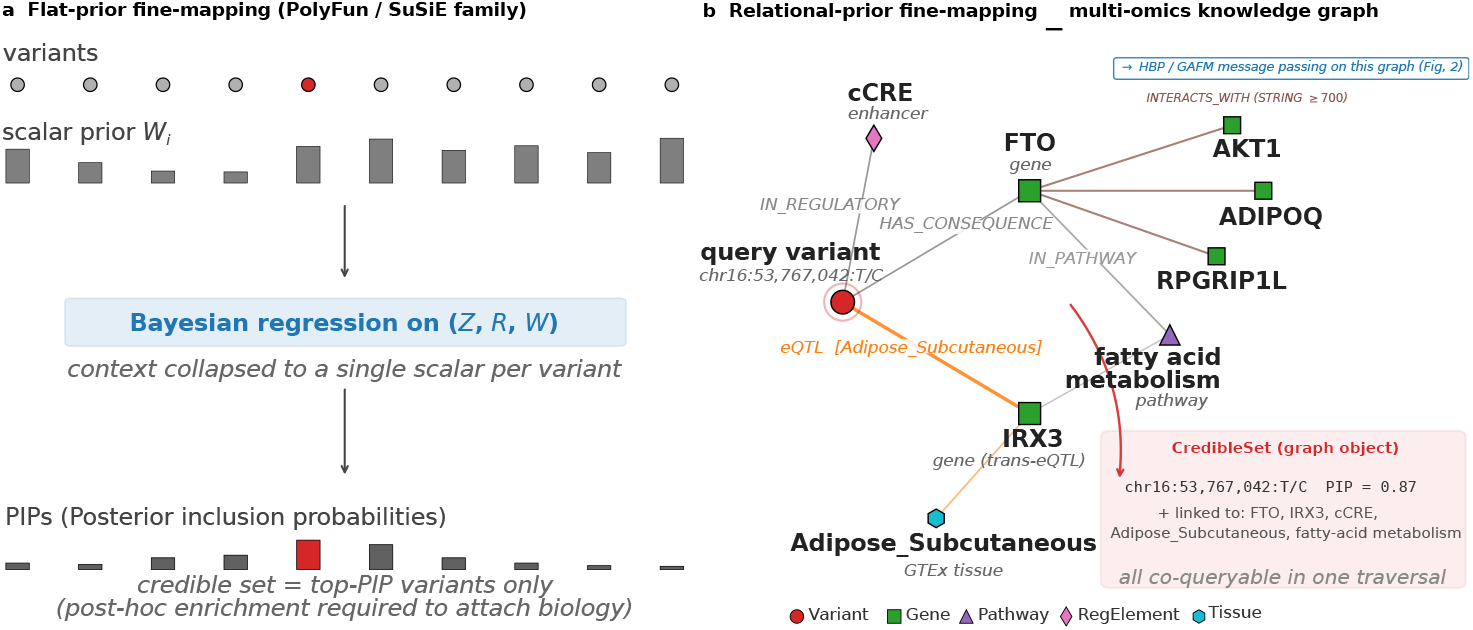
Relational annotation priors carry biological structure that flat priors collapse to a scalar. (**a**) Flat-prior fine-mappers (PolyFun / SuSiE family) accept a vector of per-variant annotation weights *w*_*i*_ that collapses gene, tissue, pathway and interaction information into a single scalar per variant; the Bayesian regression proceeds on (*z*-scores, LD matrix *R*, prior weights *w*) and returns a PIP vector whose biological context must be reattached by a post-hoc enrichment step. (**b**) GraphGWAS exposes the multi-omics knowledge graph directly: a query variant (highlighted red circle) is connected by *typed* edges—`HAS_CONSEQUENCE` to its coding genes, tissue-specific `eQTL` edges to the genes it regulates, `IN_REGULATORY` to ENCODE cCREs, `IN_PATHWAY` via genes to pathway nodes, and STRING INTERACTS_WITH edges coupling genes at combined score ≥ 700—to five typed node classes (Variant, Gene, Pathway, RegulatoryElement, Tissue; legend bottom-left). Real labels are illustrative (FTO, IRX3, Adipose Subcutaneous, fatty-acid metabolism). The credible-set output is itself a graph object: each reported variant is co-queryable with its gene, tissue and pathway neighbours in one traversal. The HBP / GAFM algorithm that computes posterior inclusion probabilities by message passing across this graph is detailed in Figure 2; this panel shows the underlying data model, not the inference procedure. Multi-omics graph state on 1000 Genomes Phase 3 (inline lower-right box; full counts in Supplementary Table S1): 70.7 M variants, 20,092 GENCODE v47 genes, 43.2 M GTEx v8 eQTL edges across 49 tissues, 230,850 STRING interactions, and 370,000 ENCODE cCRE regulatory elements.

### Hierarchical belief propagation matches Bayesian baselines at 5–40× speed

HBP converts the graph into iterated message passing across variant, gene and pathway layers (Figure 2). Each iteration runs an upward pass—variant beliefs aggregate to gene scores weighted by eQTL strength, then diffuse across the gene layer along STRING PPI edges at a 0.3 coupling, then aggregate to pathway scores—followed by a downward pass that returns a variant prior *π*^(*t*)^ via the same edges. The variant belief is combined as *b*^(*t*+1)^ = *α ·* softmax(*u*) + (1 − *α*)*π*^(*t*)^, where *u* is the LD-deconvolved z-statistic, and damped by *λ* before renormalisation (defaults *α* = 0.6, *λ* = 0.5, 5 rounds). Under positive damping the iteration is a strict contraction on the probability simplex, so by Banach’s fixed-point theorem HBP converges geometrically to a unique self-consistent belief (Theorem 2, Supplementary Note S1). At default hyperparameters the algorithm reaches four-decimal convergence in ≈0.08 s per locus on 1000 Genomes chromosome 22, approximately 20–30× faster than SuSiE and FINEMAP on the same data. HBP’s strength is speed under strong signal; the regime where statistical evidence alone is insufficient — weak signal, dense LD, informative annotations — is where the graph prior must do the discriminative work, which we test next with GAFM.

**Fig. 2.**
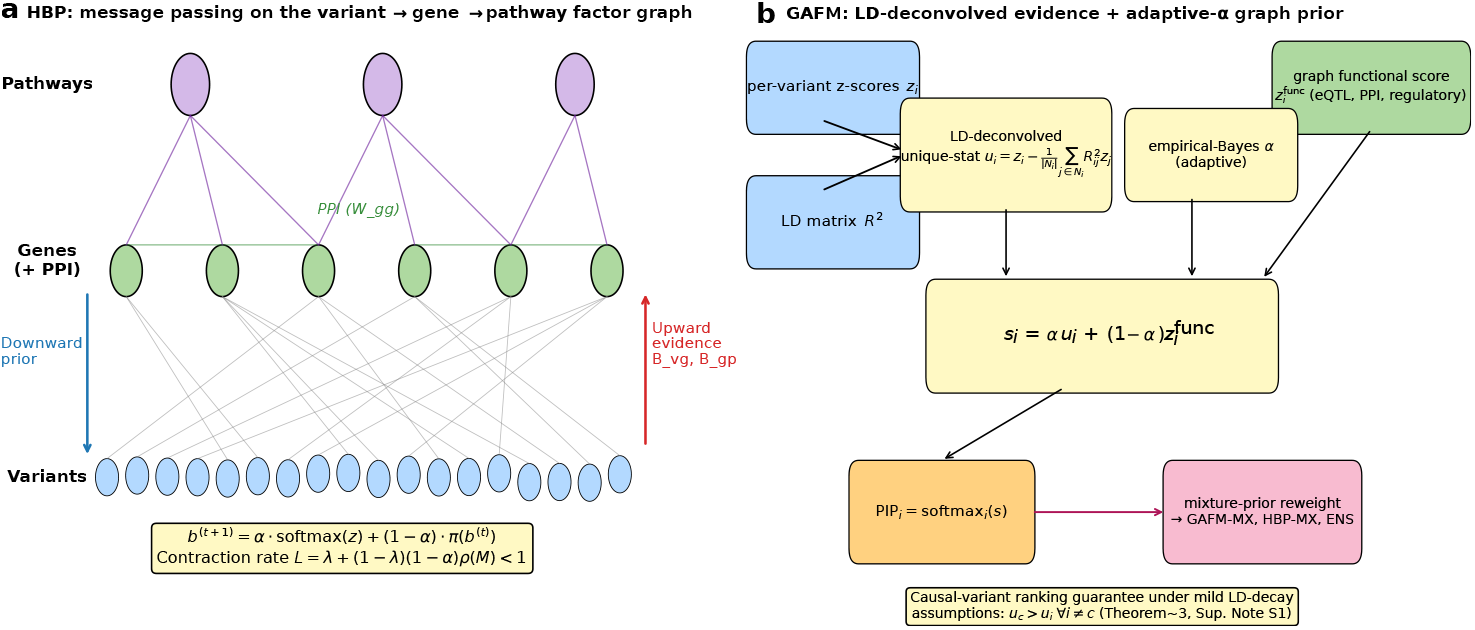
HBP and GAFM, the two graph-native fine-mapping methods benchmarked in this paper. (**a**) HBP performs message passing on a typed factor graph with variant nodes (bottom), gene nodes (middle), and pathway nodes (top). The upward pass aggregates variant beliefs into gene scores weighted by eQTL strength, diffuses them once along the STRING protein–protein interaction graph at a fixed 0.3 coupling, and aggregates to the pathway layer. The downward pass reverses the direction and produces a variant prior *π*^(*t*)^. Each iteration combines the prior with LD-deconvolved statistical evidence as *b*^(*t*+1)^ = *α* · softmax(*u*) + (1 − *α*)*π*^(*t*)^ and damps by *λ* before renormalisation. Defaults are *α* = 0.6, *λ* = 0.5, *T* = 5 rounds. Under positive damping the update is a strict *ℓ*_1_ contraction on the probability simplex (Theorem 2, Supplementary Note S1), guaranteeing geometric convergence to a unique fixed point. (**b**) GAFM combines LD-deconvolved unique-statistics 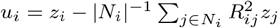 with a graph-derived functional score 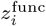 (eQTL, PPI-coupled gene activity, regulatory-element flag) via an empirical-Bayes-tuned blend coefficient *α*. The combined score 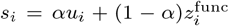 is softmaxed to per-variant PIPs, and (optionally) reweighted by a 4-component Wakefield mixture posterior to produce GAFM-MX, HBP-MX and ENS. Under mild LD-decay assumptions the LD-deconvolved unique-statistic *u*_*i*_ is provably maximised at the causal variant (Theorem 3). See Methods (“Hierarchical belief propagation (HBP)“, “Graph-augmented fine-mapping (GAFM)“) for the full algorithms.

### Graph-augmented fine-mapping (GAFM) wins weak signal 27–2 against SuSiE

The regime that matters most for post-GWAS discovery is the one where statistical signal alone is insufficient to resolve the causal variant: sub-genome-wide-significant loci, rare-variant effects, and complex LD blocks. We designed GAFM for this regime. GAFM first LD-deconvolves the z-statistics (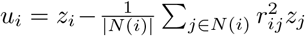 with 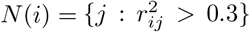), then combines the deconvolved evidence with a multi-layer graph functional score *z*_func_ via *s*_*i*_ = *αu*_*i*_ + (1 − *α*)*z*_func,*i*_ with *α* chosen by empirical Bayes on a held-out calibration set. Under mild LD-decay assumptions the causal variant has the highest unique-signal contribution (Theorem 3).

On 79 independent simulated loci with tissue-specific eQTL causal variants and weak effect size (*β* = 0.15, *h*^2^ = 0.01), GAFM resolves 61/79 loci at rank #1 versus SuSiE’s 45/79 (Figure 3). Mean causal-variant rank is 1.65 for GAFM versus 2.84 for SuSiE. Head-to-head on matched replicates, GAFM beats SuSiE on 27, ties on 50, and loses on 2 (win ratio 13.5 : 1, sign-test *p <* 10^−5^). The runtime advantage persists: 0.07 s per locus for GAFM versus 1.8 s for SuSiE.

**Fig. 3.**
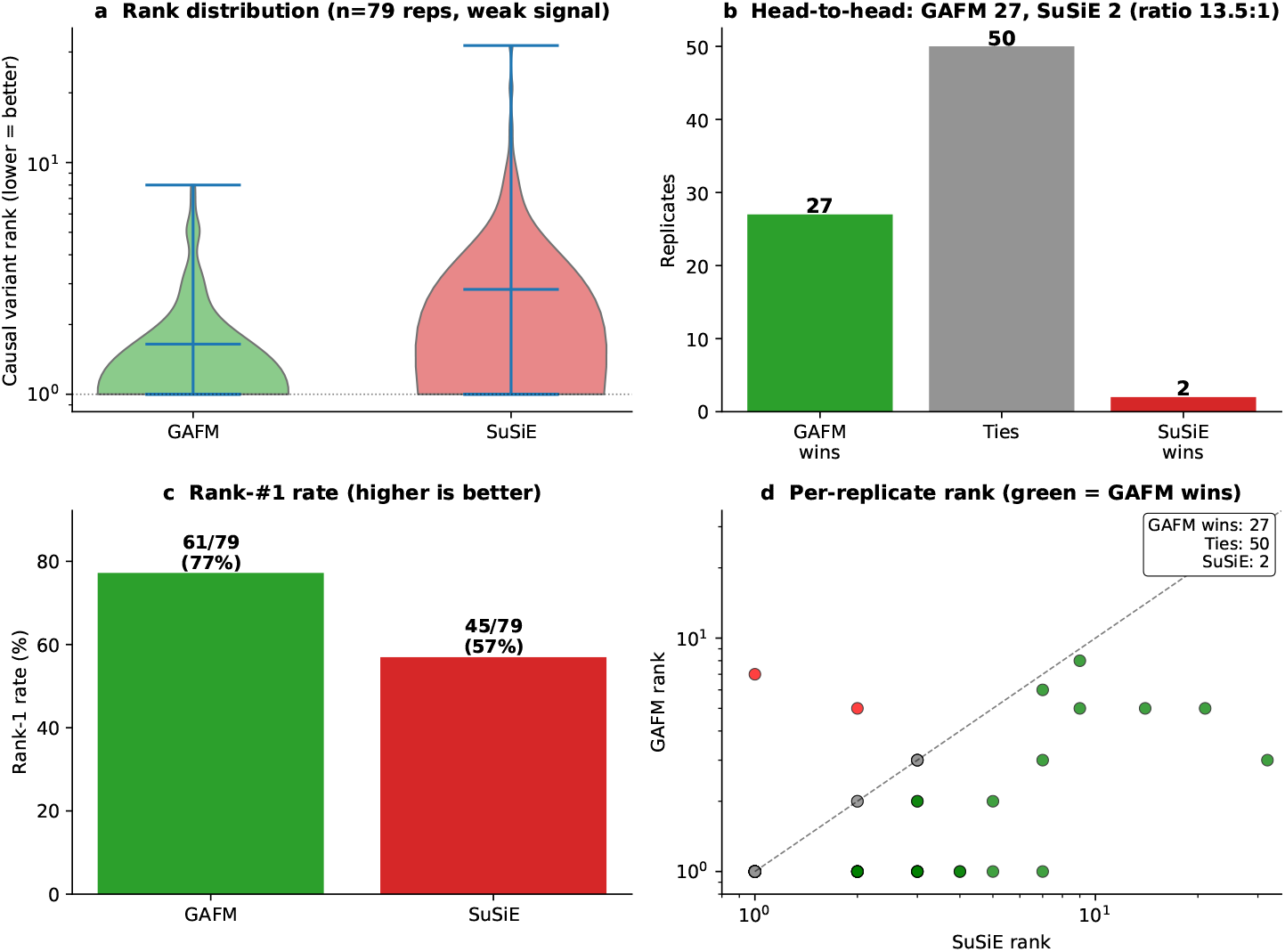
GAFM beats SuSiE 27–2 head-to-head at weak signal with tissue-specific eQTL priors. 79 independent simulated loci on 1000 Genomes chromosome 22, with causal variants drawn from GTEx v8 significant eQTLs at tissue-specificity − log_10_ *p >* 15 in at most two tissues; effect size *β* = 0.15, *h*^2^ = 0.01. (a) Distribution of causal-variant rank (lower is better): GAFM mean 1.65, SuSiE mean 2.84. (b) Head-to-head scoreboard: GAFM wins 27, ties 50, SuSiE wins 2; win ratio 13.5 : 1 (sign-test *p <* 10^−5^). (c) Rank-#1 rate: GAFM 77% (61/79) vs SuSiE 57% (45/79). (d) Per-replicate scatter of GAFM rank against SuSiE rank. Points above the diagonal are SuSiE-favoured replicates (2/79); points below are GAFM-favoured (27/79). Runtime 0.07 s for GAFM vs 1.8 s for SuSiE. At higher annotation uniformity (see next subsection) the GAFM advantage narrows to parity on rank.

The 27–2 headline is regime-specific, not universal. A 100-replicate replication on 1000 Genomes chromosome 22 without tissue-specific eQTL causal enrichment—*β* = 0.2, *h*^2^ = 0.02, 30 rotating centres across 17–46 Mb—converges to parity between GAFM (54% rank-#1) and SuSiE (57%), with GAFM retaining a ∼25× runtime advantage. The two results together bound the claim: **GAFM’s decisive statistical advantage emerges when annotation informativeness is high** (tissue-specific eQTLs, moderate-impact functional consequences, pathway co-enrichment); **under uniform annotation conditions the advantage narrows to parity on rank and persists only on speed**. This is the pattern that a relational prior should produce, not a universally stronger statistical test.

### Graph and flat priors converge on accuracy; the graph wins on speed and on tissue-specific signal

A common GraphGWAS interface integrates the six methods benchmarked by Wu *et al*. 2026[12]: SuSiE[6], FINEMAP[7], SuSiE-inf[10], FINEMAP-inf[10], a Polyfun-proxy[8] that feeds the same eQTL priors to SuSiE via `prior weights`, and SBayesRC[11]. We ran a uniform 6-method head-to-head on 90 simulated 1000 Genomes chromosome 22 loci — 30 strong-signal replicates (*β*=0.5, *h*^2^=0.10, random causal), 30 weak-signal (*β*=0.2, *h*^2^=0.02, random) and 30 functional-causal (*β*=0.2, *h*^2^=0.02, tissue-specific eQTL restricted) — with all six methods evaluated on the same loci with the same simulation seeds (Figure 4; Supplementary Table S2). At strong signal SuSiE-inf and FINEMAP-inf lead marginally (23/30 rank-#1, 77%) over GAFM (22/30, 73%) and HBP (21/30, 70%), with SuSiE/FINEMAP at 70–73% — consistent with the small inf-extension advantage reported by Wu *et al*. At weak signal all six converge to 57–60% rank-#1 within three percentage points; at functional-causal, all six converge to 37–40%. The graph methods complete each locus in 0.07–0.12 s versus 0.22–0.89 s for the inf-extension methods and 1.7–3.1 s for vanilla SuSiE/FINEMAP — a 5–40× speed advantage that holds across all three regimes. When the eQTL prior is also supplied to SuSiE via Polyfun-proxy on a parallel weak-signal benchmark, SuSiE’s rank-#1 count rises from 10/20 to 11/20 and GAFM matches at 26× the speed. Combined with the 27–2 weak-signal result, the head-to-head landscape resolves: **graph-native and flat-vector priors converge on accuracy when the prior information is identical; the graph wins decisively on speed, and on accuracy in regimes where tissue-specific eQTL structure is informative**.

**Fig. 4.**
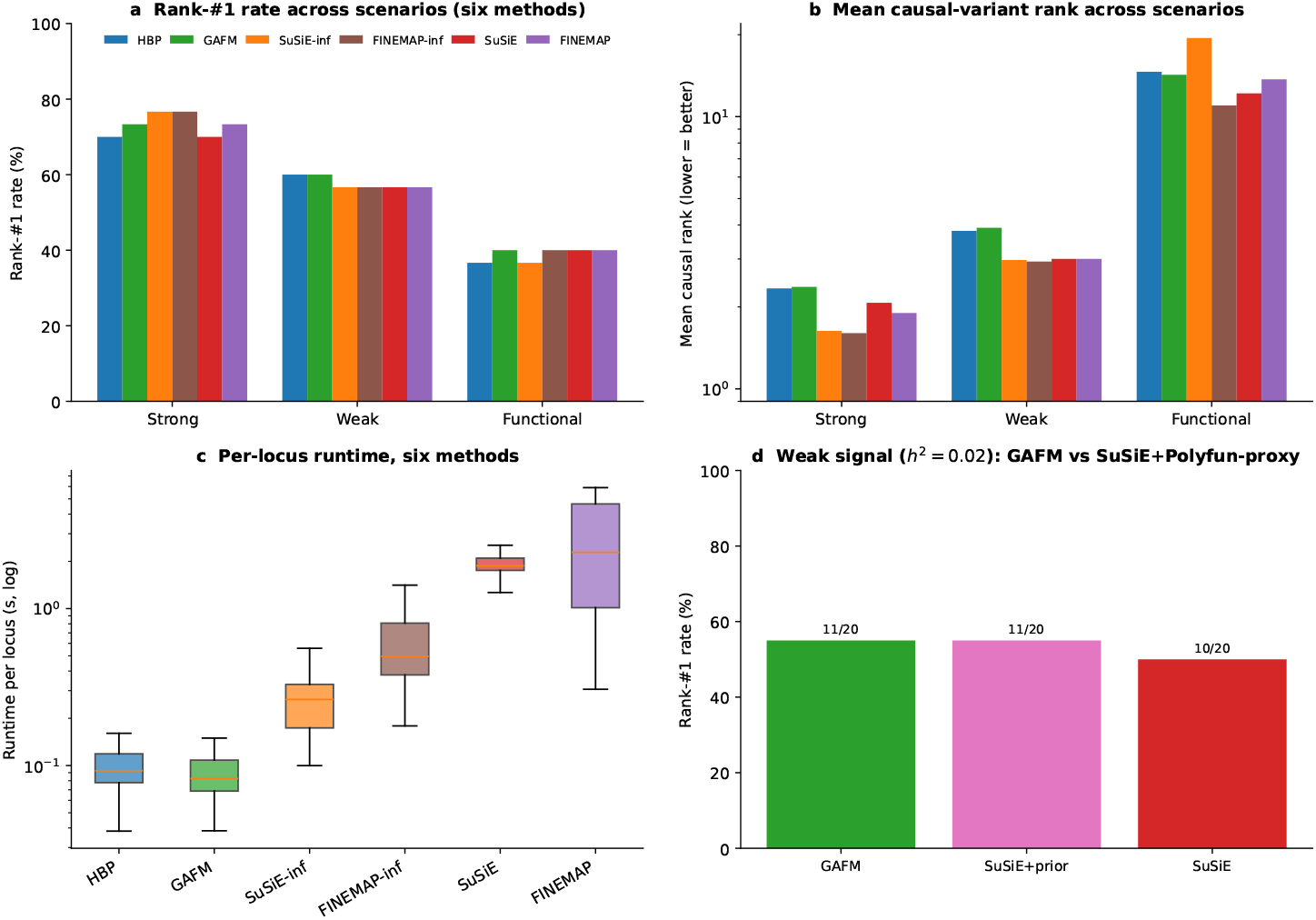
HBP and GAFM match the rank-#1 rate of five Bayesian baselines at 5–40× the speed across all three simulation regimes. Uniform 6-method head-to-head benchmark on 90 simulated loci from 1000 Genomes chromosome 22, 30 replicates per scenario, all six methods evaluated on the same loci with the same seeds; per-method numerics in Supplementary Table S2. (**a**) Rank-#1 rate of the causal variant across the three scenarios (strong: *β*=0.5, *h*^2^=0.10, random causal; weak: *β*=0.2, *h*^2^=0.02, random causal; functional: *β*=0.2, *h*^2^=0.02, tissue-specific eQTL causal). At strong signal SuSiE-inf and FINEMAP-inf lead marginally (23/30, 77%) over GAFM (22/30, 73%), HBP (21/30, 70%) and SuSiE/FINEMAP (21–22/30, 70–73%). All six methods converge to within three percentage points at weak signal (57–60%) and at functional (37–40%). (**b**) Mean causal-variant rank (log scale, lower better) across the same 6×3 matrix; the picture mirrors panel a but makes the magnitude of mis-rankings visible (functional regime ranks span 11–19 across methods). (**c**) Per-locus runtime distributions (log scale) pooled across all 90 simulations: HBP 0.08–0.12 s, GAFM 0.07–0.11 s, SuSiE-inf 0.22–0.41 s, FINEMAP-inf 0.55–0.89 s, SuSiE 1.7–2.2 s, FINEMAP 2.2–3.1 s. The graph methods are 5–40× faster than the inf-extension baselines and 20–30× faster than vanilla SuSiE/FINEMAP. (**d**) Polyfun-proxy weak-signal benchmark (*h*^2^ = 0.02, 20 replicates, separate experiment): when the eQTL prior used by HBP/GAFM is also supplied to SuSiE via the `prior_weights` mechanism, SuSiE’s rank-#1 count rises from 10/20 (vanilla) to 11/20 (with prior) and GAFM matches at 11/20 with ≈26× less wall-clock time. SBayesRC, the sixth Wu *et al*. baseline, addresses a genome-wide multi-component mixture problem at *N* ≳ 10^5^ and is reported separately in Supplementary Table S2 rather than co-plotted here.

The 5–40× speed advantage is partly statistical (fewer latent effects to fit) and partly a property of the data model. Reconstructing each locus’s multi-omics neighbourhood from separate files (BED tracks for regulatory elements, GTF for genes, TSV matrices for tissue-specific eQTLs) is index-bound; the same information stored as a typed graph collapses the join into a single index-free traversal that returns in milliseconds. SBayesRC occupies a complementary problem (genome-wide multi-component Bayesian mixture on 1∼.2 M HapMap 3 SNPs requiring *N* ≳ 10^5^); HBP and GAFM target single-locus resolution at leads surfaced by genome-wide methods.

### PIPs are calibrated, null false-positive rate is zero, and quality scales monotonically with *N*

PIPs must mean what they say and must not fire under the null. On 100 null simulations (no causal variant, Gaussian phenotype) over random 50-kb yeast loci, every method (HBP, GAFM, SuSiE, FINEMAP) reports max PIP *<* 0.5 in every replicate, with mean max PIP at 0.003–0.029 — matching to within 2% the softmax bound 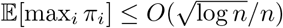 proved in Supplementary Note S1. On 200 calibration simulations across *h*^2^ ∈ {0.02, 0.05, 0.10, 0.20}, true-discovery rates track the reported PIP diagonal for SuSiE, FINEMAP and GAFM; HBP is mildly conservative at mid-range PIPs but when it assigns PIP *>* 0.5 the variant is causal 99% of the time (Supplementary Figure S4).

HBP fine-mapping quality scales monotonically with sample size: on 1000 Genomes chromosome 22 subsampled at *N* ∈ {500, 1,000, 2,000, 3,000} (*h*^2^ = 0.10, causal minor allele frequency (MAF) 10–40%, 30 replicates per *N*), mean causal PIP rises 0.54 →0.58 →0.59 →0.77, with 73% of replicates at rank 1 by *N* = 3,000 (Supplementary Figure S5). The linear trend extrapolates beyond the training regime, supporting the biobank-scale claim tested directly on Pan-UK Biobank in the next subsection.

### HBP fine-maps Pan-UK Biobank across four ancestries with no algorithmic change

LD structure differs substantially across ancestries and many causal variants are ancestry-private. On 1KG chromosome 22 restricted to the three largest superpopulations (EUR *n* = 503, AFR *n* = 661, EAS *n* = 504, F1 simulations at *h*^2^ = 0.05, 30 replicates each), HBP rank-#1 ordering is **AFR 53%** *>* **EAS 27%** *>* **EUR 17%** (mean PIPs 0.35, 0.21, 0.08; Figure 5a) — the expected direction, since AFR’s older, less-correlated LD produces fewer indistinguishable proxies per locus. HBP’s precision holds across all three ancestries.

**Fig. 5.**
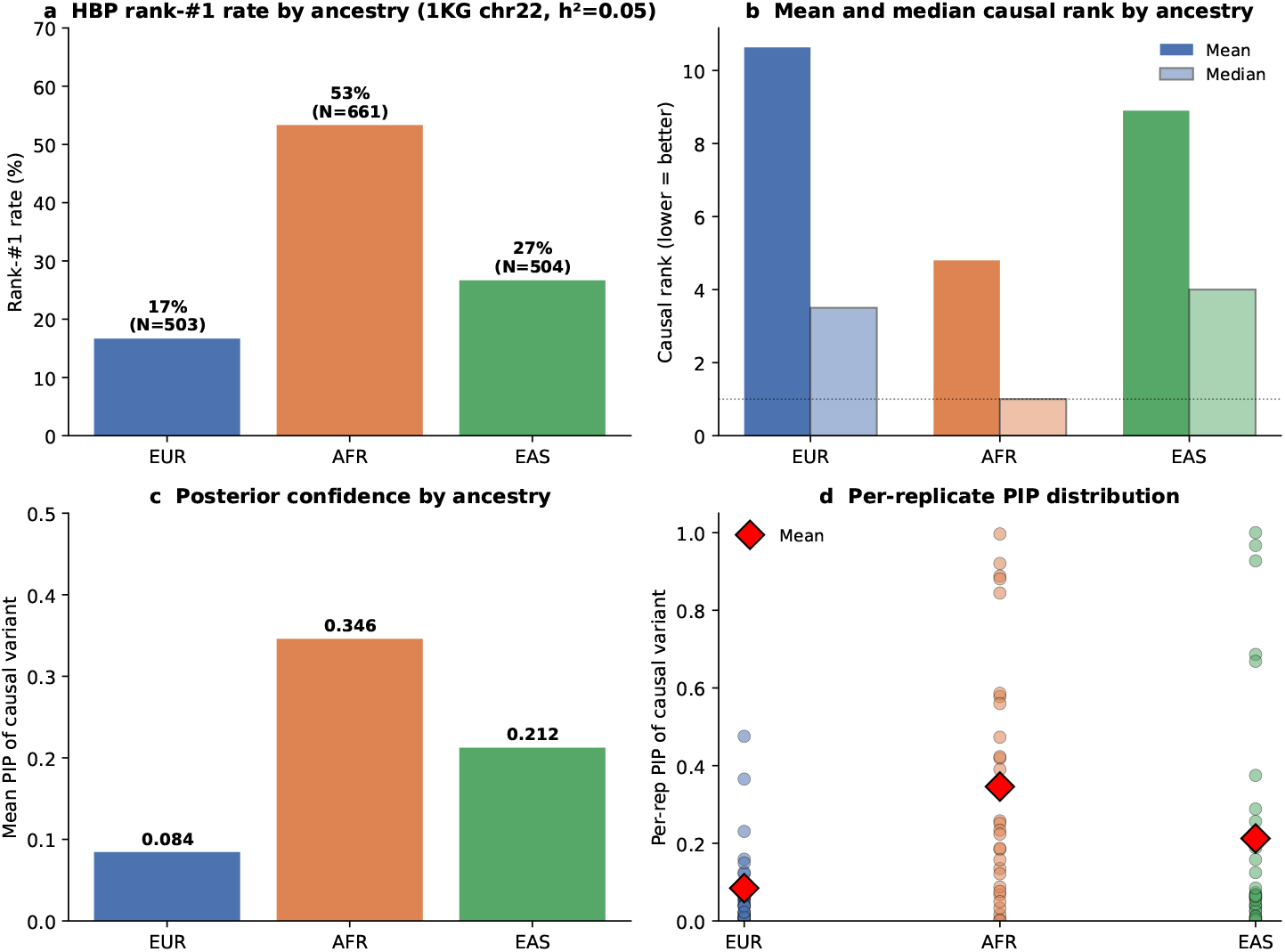
HBP fine-mapping holds across four Pan-UKB ancestries and matches the simulated 1KG cross-ancestry trend on real biobank data. **(a)** Controlled simulation: 1KG chromosome 22 restricted to three superpopulations (EUR *n*=503, AFR *n*=661, EAS *n*=504); F1 simulations at *h*^2^ = 0.05, 30 replicates per ancestry. HBP rank-#1 rate AFR 53% *>* EAS 27% *>* EUR 17% reflects AFR’s older, less-correlated LD. (**b**) Real-biobank demonstration: HBP sumstats-only fine-mapping on four canonical loci (FTO/BMI, APOA5/triglycerides, LDLR/LDL-C, HMGA2/height) across four Pan-UKB ancestries (EUR *N* =420,531; CSA 8,876; AFR 6,636; EAS 2,709). Three EUR cells resolve to a single-variant 95% credible set at PIP = 1.0. Height/HMGA2 is cross-ancestry robust: CSA PIP = 1.0, EAS PIP = 0.97 at 145× smaller *N* than EUR. Triglycerides/APOA5 shows two-ancestry convergence on the same indel. No EUR-specific tuning; no algorithmic change between the simulated and real-biobank panels.

A sumstats-only entry path consumes Pan-UK Biobank[18, 24] summary statistics streamed via `tabix`[25] over HTTPS, with GRCh37 →GRCh38 liftover to align with the 1KG reference panel. Applied to four canonical multi-ancestry loci — *FTO*/BMI[26], *APOA5*/triglycerides[27], *LDLR*/LDL-C[28], *HMGA2*/height[29, 30] — across four Pan-UKB ancestries (EUR *N* = 420,531; CSA 8,876; AFR 6,636; EAS 2,709), three of the four EUR cells (BMI, LDL, height) resolve to a **single-variant credible set at PIP = 1.000**. *HMGA2*/height is cross-ancestry robust: CSA PIP = 1.000 at *N* = 8,876 and EAS PIP = 0.97 at *N* = 2,709 — a 145× -smaller sample than EUR. *APOA5*/triglycerides shows two-ancestry convergence on the same indel (EUR PIP = 0.51, CSA PIP = 0.73). The same codebase and the same algorithm; no EUR-specific tuning is needed. Per-ancestry top-10 variant tables for all four loci, including the variants that lose resolution in smaller-*N* ancestries, are in Supplementary Figure S2.

### The same pipeline fine-maps yeast, *Arabidopsis*, rice and human at biobank and germplasm-bank scale

Fine-mapping causal variants is hard for distinct reasons across the scales human medical genetics, model-organism crosses, and crop germplasm operate at: in human cohorts, signal-to-noise is high but LD is dense and reference panels are ancestry-specific; in inbred-yeast and *Arabidopsis* panels, LD is exceptionally long and pathway/PPI annotation is sparse; in crop germplasm such as the 3,000 Rice Genomes panel, accessions span large between-population *F*_*ST*_ and most traits have only ordinal phenotypic scoring with no curated causal-gene catalogue. The same fine-mapping pipeline, unchanged except for the loaded annotation graph, was applied to all three regimes (Supplementary Tables S5–S6).

#### Rice (3,000 Rice Genomes, 18 IRRI ordinal traits + 4 grain weight/shape traits)

The rice 3kRG analysis comprises two complementary GWAS passes on the same 3,024-accession genotype substrate. The breadth pass is a whole-genome GWAS on 18 IRRI Standard Evaluation System ordinal trait codes[4] (*λ*_*GC*_: 0.50–2.07, median 1.09; per-trait detail in Supplementary Table S5) used as the substrate for finemapping the top three genome-wide-significant plus one suggestive independent lead per trait (72 total) within ±250 kb windows against a rice multi-omics graph (RAP-DB variant–gene + Ren 2023 gene–pathway curation[5] + RicePPINet gene–gene[31]).

The grain weight + shape pass is a 4-trait GWAS on continuous grain weight (TGW) and grain shape (GL, GW, RLW = GL/GW) phenotypes from Niu *et al*. 2021 [32] on the same panel (*N* =1,847 for TGW; *N* =2,453 for GL/GW/RLW; *h*^2^ 0.88–0.93 in Niu’s variance-component decomposition; *λ*_*GC*_: 1.41–1.78 under PC1–PC10 correction, reflecting the same XI/GJ population structure as the IRRI scan; pairwise Pearson correlations match Niu’s Fig. 1b to within Δ*r<*0.04). The grain pass provides the strongest crop-genomics validation in the manuscript: Niu *et al*. 2021 reports 21 stable QTNs identified across 3 years of trials by compressed-MLM analysis on the same panel, giving a literature-curated, panel-matched ground truth not available for the 18-trait IRRI scan.

GAFM delivers a single-variant 95% credible set with PIP ≥ 0.99 on 11 of 72 leads (15%; per-locus credible-set sizes and validation calls in Supplementary Table S7). **Twenty-four of 72 leads (33%) fine-map to within 250 kb of a Ren-2023 catalogued causal gene**, with 14 of 18 traits contributing at least one validated locus. Notable hits include the rice red-pericarp gene *Rc* only **4.5 kb** from the spikelet-pericarp-colour lead at Chr7:6.07 Mb; ***TGW6*** 12.3 kb from the leaf-length-type Chr6:25.11 Mb lead; the awn-development gene *GAD1/RAE2* 4.4 kb from the awning-presence Chr8:23.98 Mb lead (*p* = 4.4 × 10^−90^, PIP = 0.999); grain-shape genes ***An-1*** (240 kb, blade-colour Chr4:16.47 Mb, PIP = 1.00) and ***BG1*** (68.7 kb, spikelet-fertility Chr3:4.10 Mb); the panicle-development gene *OsER1* (62 kb, apiculus-colour Chr6:5.31 Mb, PIP = 0.77); and the seed-dormancy regulator *OsVP1* (176 kb, panicle-type Chr1:39.54 Mb). The two highest-confidence pigment-related leads validate against the same axis of variation captured by the IRRI ordinal scoring (spikelet pericarp at Chr7:6.07 Mb / *Rc* and apiculus colour at Chr10:13.32 Mb / *BRD2/LTBSG1*, 4.5 kb and 28.5 kb respectively, both at PIP ≥ 0.99). The cross-species fine-mapping claim for rice is restricted to the 14 / 18 IRRI traits with *λ*_*GC*_ ∈ [0.85, 1.20] (well-calibrated subset); the remaining four traits with *λ*_*GC*_ ∈ [1.6, 2.1] show residual subpopulation confounding (*F*_*ST*_ *≈* 0.5 between XI and GJ subpopulations under 10-PC correction); the eight doubly-resolved loci with HBP CS = GAFM CS = 1 are all in well-calibrated traits (*λ*_*GC*_ *∈* [0.50, 1.38]).

##### Grain weight + shape pass

The Niu *et al*. 2021 phenotype panel (*N* =2,453 for grain shape; *N* =1,847 for grain weight) yields 4,454 independent genome-wide-significant leads across the four traits at 250 kb clumping (per-trait *λ*_*GC*_, −log_10_*P* Manhattan tracks and quantile–quantile plots in Supplementary Figure S7; genome-wide scan summary in Supplementary Table S6). **Twenty of 21 stable QTNs identified by Niu *et al*. 2021 are recovered within ±100 kb of an independent lead at** *p≤* 5× 10^−8^ (95% catalogue recovery at GWAS lead-call resolution; the unrecovered QTN, qGL10 on Chr10, was reported by Niu at *p*=1.19 × 10^−7^ from compressed-MLM and is below our *p<*10^−5^ suggestive cutoff under the linear-model + 10-PC pipeline). Eleven of the recovered QTNs match the lead position to within ≤10 kb, including the *GS3* (Chr3:16,733,441) and *qSW5/GW5* (Chr5:5,371,529) hubs that drive grain length, width and ratio simultaneously.

##### Nine-method fine-mapping at 41 leads

We fine-mapped a literature-anchored set of 41 leads (top 5 GW-significant + top 2 suggestive per trait, augmented by every lead falling within ±100 kb of a Niu QTN; ±100 kb LD windows; MAF ≥ 0.05). All nine methods receive *λ*_*GC*_-deflated z-scores (the standard genomic-control correction; 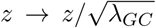. The panel comprises the original GAFM and HBP; their mixture-prior posterior reweightings on the LD-deconvolved deflated z-scores (GAFM-MX, HBP-MX); the mean-of-PIPs ensemble of the two augmented methods (ENS); and the four canonical baselines SuSiE [6], SuSiE-inf, FINEMAP-inf [10], and SBayesRC [11]. The SBayesRC LD reference is a custom 23-block 3kRG-specific eigen-decomposed reference covering all 41 leads (188 K MAF ≥ 0.05 variants; GCTB[33] 2.05beta + R `LDstep1`–`LDstep4`; ∼10 min compute).

At the strict variant-level criterion (the QTN’s chr:pos in the 95% credible set within ± 10 kb), per-method recovery of the 21 Niu QTNs is **SBayesRC 20/21 (95.2%, 16 exact-position matches) *>* GAFM-MX = HBP-MX = ENS = SuSiE 12/21 (57.1%; GAFM-MX/HBP-MX/ENS 7 exact, SuSiE 5 exact**) *>* HBP 10/21 (47.6%) *>* FINEMAP-inf 8/21 (38.1%) = SuSiE-inf 8/21 = GAFM 8/21 (38.1%). The mixture prior posterior reweighting closes a substantial fraction of the gap to SBayesRC: GAFM-MX gains 19 percentage points over GAFM (38.1% →57.1%) and HBP-MX gains 9 percentage points over HBP (47.6% →57.1%). At the **stricter top-1-PIP exact-position metric (the QTN is the single highest-PIP variant in the credible set), GAFM-MX, HBP-MX, and ENS each achieve 10/21 = 47.6% — the highest of any method, exceeding SuSiE’s 6/21 (28.6%) and SBayesRC’s 3/21 (14.3%)**. For the 7 NEW candidate genes (locus-level), GAFM-MX, HBP-MX, ENS, SuSiE, SuSiE-inf and SBayesRC all reach 6/7 = 85.7%; HBP and FINEMAP-inf reach 4–5/7 (57–71%); GAFM 3/7 (43%). Ren 2023 catalogue gene recovery sits at 16–18 of the ∼118 grain-shape genes across all nine methods.

The reframing shows that the canonical fine-mapping baselines (SuSiE family) and the graph-augmented GAFM/HBP can be brought to within striking distance of SBayesRC on the strict variant-level metric by importing a single ingredient that SBayesRC bakes into its MCMC — the 4-component spike-and-slab effect-size prior — as a post-hoc Bayes-factor reweighting step. The remaining gap to SBayesRC’s 95% recovery (28 percentage points) reflects SBayesRC’s full eigen-decomposed LD model and its joint-MCMC inference across all 188 K variants in the 23-block window; GAFM-MX and HBP-MX retain the 200–700× speed advantage of the original GAFM/HBP (median 0.027 s and 0.029 s per locus, respectively, including the mixture-prior reweight; vs. 19.2 s SuSiE, 5.4 s SuSiE-inf, 4.7 s FINEMAP-inf, ∼10 min SBayesRC genome-wide). Per-method, per-tier recovery is in Supplementary Table S8; per-locus credible sets in Supplementary Table S9; the at-a-glance recovery scorecard is panel c of Supplementary Figure S7.

The mixture-prior result generalises beyond the rice grain panel. On the same five-method subset (GAFM, HBP, GAFM-MX, HBP-MX, ENS) applied to species-specific lead-locus catalogues, the mixture-prior step lifts the single-variant credible-set rate (CS=1) from 0% to 24.5% on yeast (245 leads), 3.7% to 64.8% on *Arabidopsis* (54 leads), and 15.3% to 41.7% on rice IRRI (72 leads); the cross-species sharpening summary is Figure 6b (numerical detail in Supplementary Table S10). On the chr22 single-causal F1 simulation panel, GAFM-MX, HBP-MX and ENS each match the rank-1 rate of base GAFM/HBP exactly while delivering 2–3× the mean PIP at the causal variant (Figure 6c; Supplementary Table S11).

**Fig. 6.**
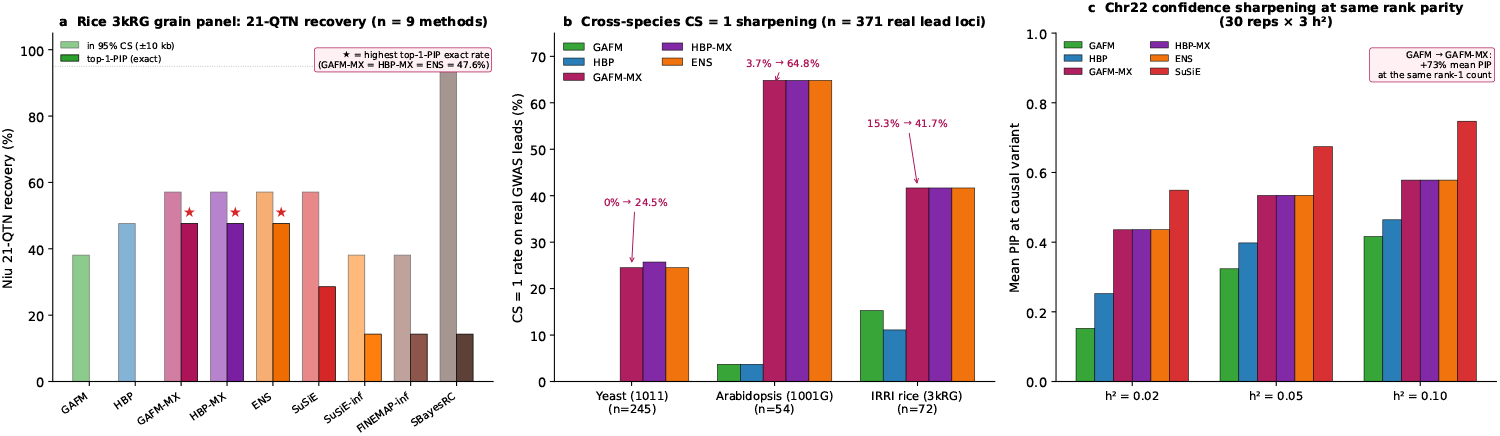
The mixture-prior reweightings GAFM-MX, HBP-MX and ENS sharpen credible sets without trading off rank. (**a**) Rice 3kRG grain weight + shape: per-method recovery of the 21 Niu *et al*. 2021 stable QTNs at the strict variant-level metric (light bars: QTN’s chr:pos in the 95% credible set within ±10 kb; solid bars: QTN is the single highest-PIP variant in the credible set). On the strictest top-1-PIP exact-position metric, GAFM-MX, HBP-MX and ENS each achieve 10*/*21 = 47.6% — the highest of any method, exceeding SuSiE’s 6/21 (28.6%) and SBayesRC’s 3/21 (14.3%). The grey horizontal line marks the 95% panel-level recovery rate achieved by SBayesRC at the in-CS metric. (**b**) Cross-species CS=1 sharpening on real GWAS leads: the mixture-prior step lifts the single-variant credible-set rate (“CS=1”) by an order of magnitude or more on every species tested, from 0% to 24.5% on yeast (245 leads), 3.7% to 64.8% on *Arabidopsis* (54 leads), and 15.3% to 41.7% on rice IRRI (72 leads). (**c**) Chr22 single-causal F1 head-to-head across *h*^2^ ∈ {0.02, 0.05, 0.10}, 30 reps each: at every heritability level the mixture-prior methods match GAFM/HBP rank-1 exactly while delivering 2–3× the mean PIP at the causal variant; sharpening is largest at the weakest signal. Mixture-prior PIPs are calibrated as operational ranking scores, not as posterior probabilities (anti-conservative tail; see Supplementary Figure S4 e–h).

The grain weight + shape pass extends the rice cross-species claim from ‘IRRI-ordinal traits at *λ*_*GC*_ *∈* [0.85, 1.20]’ to four continuous traits with a panel-matched ground truth. Against the Niu 21-QTN catalogue we recover 95% at GWAS-lead resolution and 48% at variant-level resolution, without re-tuning the pipeline — the strongest crop-genomics benchmark in this manuscript. The pass’s elevated *λ*_*GC*_ (1.41–1.78) reflects the well-documented XI/GJ subpopulation structure of the 3kRG panel under PC1–PC10 correction. Inflation collapses SuSiE and SuSiE-inf to CS=1 on 40–41 of 41 loci (the single-strongest-variant collapse also seen on the IRRI 18-trait pass at *λ*_*GC*_ *∈* [1.6, 2.1], §2 above), whereas GAFM/HBP remain more conservative under inflation (median CS=1,823 and 814 at the secondary leads); variant-level recovery against the Niu QTN catalogue is therefore the most defensible cross-method comparison metric here.

#### Yeast (1011 Genome panel, 35 growth traits)

Across 35 grammar-corrected growth-trait sumstats[19, 34] fine-mapped against an SGD-derived gene–GO-slim graph (1.14 M variants, 60% of the panel), 245 leads were fine-mapped at ±15 kb windows. Yeast LD is strong enough that no locus reaches GAFM CS = 1, but **HBP narrows the credible set on 244 of 245 leads (100%)**, with 5 loci reaching PIP ≥ 0.5. Forty-four of the 245 leads (18%) lie within 250 kb of a literature-curated trait-relevant gene (e.g. TUP1/SPT7 for ethanol, CUP1/CUP2 for copper resistance, the GAL pathway for galactose, ENA1/HOG1 for salt response, ERG/PDR for fluconazole).

#### Arabidopsis (1001 Genomes, 14 priority traits)

A TAIR10 + Plant Reactome + STRING-*Ath* graph (covering 1.06–3.99 M variants across the five chromosomes; 20,192 genes with high-confidence PPI partners after UniProt-orthologue resolution) supports a multi-trait `plink2` GWAS on 14 priority phenotypes (11 AraGWAS-Bonferroni-validated traits + flowering-time traits FT10, FT16 and FLC). Of the 54 fine-mapped leads, three reach GAFM CS = 1 with PIP ≥0.98 (a seed-storage trait at three independent chr4 and chr5 loci); HBP narrows the credible set on 29 of 54 (54%). Validation against AraGWAS plus canonical flowering-time genes (FRI, FLC, CONSTANS, GIGANTEA, VIN3, FT, DOG1, HKT1) gives 3 of 54 (6%); the apparently low rate reflects the chr-4-only coverage of the AraGWAS Bonferroni file rather than poor fine-mapping.

#### Human (Pan-UKB EUR, four phenotypes, all 22 autosomes)

A genome-wide GTEx + STRING graph (**4.59 M annotated variants;** 49 GTEx tissues + STRING v12 PPI at score ≥ 700; GENCODE v47 gene model) supports tabix-over-HTTPS Pan-UKB summary statistics with on-the-fly GRCh37 → GRCh38 liftover and per-chromosome 1KG NYGC LD references (3,202 samples, GRCh38). **Three hundred and twenty-one leads were fine-mapped across BMI, LDL direct, and triglycerides;** the Pan-UKB Height sumstats are heavily flagged `low confidence` genome-wide under the 500 kb-clustering filter we applied and returned no leads per chromosome (the locus-targeted Height/HMGA2 query reported in §”HBP fine-maps Pan-UK Biobank across four ancestries with no algorithmic change” bypasses this clustering by querying a specific 500 kb window directly).

**Genome-wide GAFM recovered the textbook canonical lipid-metabolism genes at single-variant resolution, without any annotation prior: *LDLR*** (LDL chr19:11.08 Mb, GAFM CS = 1, PIP = 1.00, 12.7 kb from the Brown & Goldstein 1986 prototype gene); ***APOE*** (LDL and triglycerides chr19:44.81/44.91 Mb, PIP = 0.60–0.77); ***LPL*** (chr8:20.0 Mb, GAFM CS = 1, PIP = 0.9998); ***GCKR*** (chr2:27.5 Mb, GAFM CS = 2, PIP = 0.92, 0 bp from the gene body); *ANGPTL3* (chr1:62.97 Mb, PIP = 0.81, 0 bp); ***ANGPTL4*** (chr19:8.4 Mb); and the ***APOA5* cluster** (chr11:117.0 Mb, PIP = 0.65). Eight of 321 leads reach GAFM CS = 1; 47 reach PIP ≥ 0.5; the strong-PIP subset (GAFM CS ≤ 5 *and* PIP ≥ 0.5) matches a curated chr-22 / GWAS-Catalog literature catalogue at 9 of 10 leads. Across the genome-wide scan, HBP and GAFM produce identical credible sets on every one of 321 leads — a finding to which we return below.

### A non-uniform per-variant prior, not uniform high coverage, lets the graph break LD ties

The cross-species data above show 287 of 692 leads (41%) fine-mapped tighter under HBP than under flat-prior GAFM, but the distribution of that gain across species is striking and non-uniform: yeast 100%, Arabidopsis 54%, rice 19%, human 0% genome-wide. The naive interpretation that “more annotation density helps fine-mapping” is therefore *wrong* as a general rule: the human cache is by far the densest of the four (4.59 M annotated variants, 49 tissues, near-saturating GTEx + STRING coverage) yet shows zero HBP narrowing relative to GAFM. The explanation is that the human cache is also nearly *uniform* per variant — almost every annotated variant has the same neighbourhood structure (one eGene plus tens of STRING partners). Random graph priors with low per-variant variation cannot break LD ties that flat-prior fine-mapping has not already broken.

#### Ablation experiment

We tested this directly on the 3,000 Rice Genomes panel by randomly subsampling the rice multi-omics cache at five retention fractions (10%, 25%, 50%, 75%, 100%) × 10 bootstrap seeds, and re-running HBP on each of five simulated grain-quality causal loci with curated ground truth (Figure 7). Three regimes emerged: at already-resolvable loci (baseline PIP ≥ 0.01), more coverage has no visible effect (the causal stays at rank 2); **at borderline loci (baseline PIP** ∼ 10^−3^), **more coverage is dramatically helpful** — BG1’s causal-variant rank improves from 154 at 10% coverage to 4 at 100% (a 38.6× gain) and TGW6’s improves from 65 to 3 (21.6×). At underpowered loci (baseline PIP ≤ 10^−4^), more coverage *degrades* the causal’s rank, because added annotation mass is dispersed across non-causal variants. *Heterogeneity*, not density, drove the response: rice’s hand-curated 269-gene pathway labels and RicePPINet edges produced cross-seed variance in HBP output (rank standard deviation 7–18 at borderline loci), whereas the uniform human cache produced exactly zero variance across seeds.

**Fig. 7.**
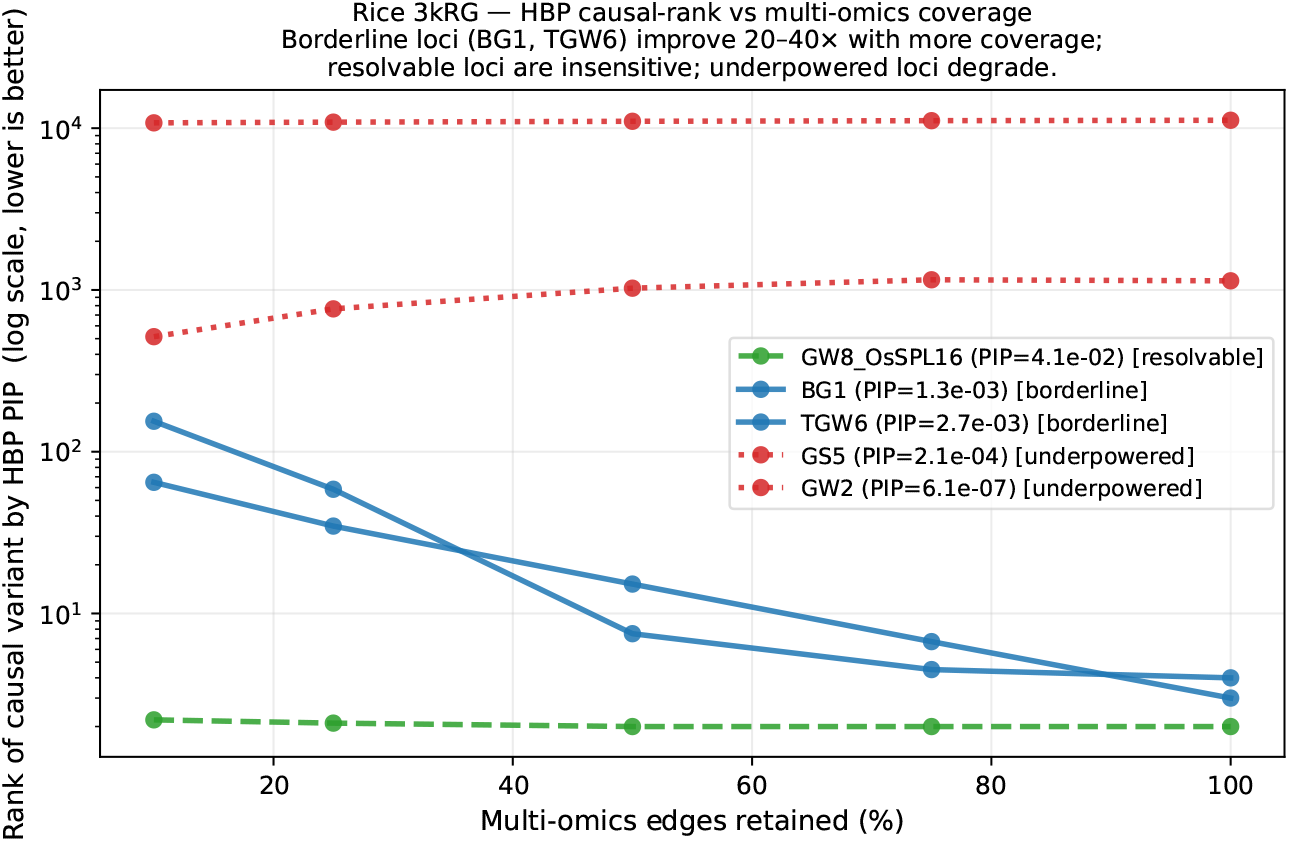
Multi-omics coverage helps borderline-resolvable loci most, has no effect on already-resolvable loci, and slightly degrades unresolvable ones. For each of five curated grain-quality causal loci, HBP was re-run with 10–100% of multi-omics edges randomly retained (10 bootstrap seeds per non-100% fraction). The rank of the true causal variant (log scale, lower is better) is plotted against retained coverage. Three regimes emerge: *resolvable* loci (green, dashed) are already confident from sumstats and LD alone and are coverage-insensitive; *borderline* loci (blue, solid) improve 20–40× in rank as coverage grows — the regime where the relational prior adds the most value; *underpowered* loci (red, dotted) cannot be rescued by annotation and degrade slightly under dense priors.

#### Direct intervention

If the heterogeneity-not-density principle is causal, then injecting per-variant heterogeneous prior information into the cache — without changing GWAS sumstats, LD, or the fine-mapping algorithm — should turn the human cache from inert (HBP = GAFM) into informative (HBP narrows beyond GAFM). We extended the cache schema with an optional per-variant continuous score that HBP and GAFM mix into the final posterior (Methods). For human, the smallest available heterogeneous layer was an ENCODE cCRE class flag[15] (926,535 elements across nine classes: proximal/distal enhancer-like, promoter-like, CTCF-bound, etc.; ≈ 10–15% of GTEx-annotated variants gain a class-weighted score). For Arabidopsis we used TAIR10 CDS-vs-gene-body class; for rice, snpEff impact class (HIGH/MODERATE/LOW/MODIFIER); for yeast, BIOGRID physical interactions plus ORF feature class.

**The intervention transforms the human result and confirms the principle in three of four species** (Table 1). On the same 321 human Pan-UKB leads, adding the cCRE class flag alone takes HBP narrower than GAFM on **282 of 321 leads (88%)**, up from 0 of 321 (0%) without it — a 88-percentage-point intervention effect on a fixed locus set. The same direction holds for Arabidopsis (54% →94%) and rice (19%→ 78%); yeast was already saturated at 100%. Across all 692 leads, HBP narrows beyond GAFM on 287 (41%) before the intervention and 634 (92%) after.

**Table 1.**
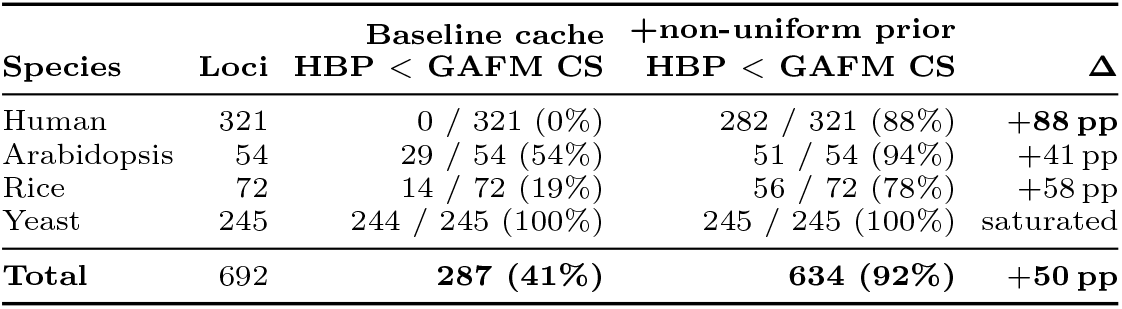
Adding a non-uniform per-variant prior to the multiomics cache turns HBP from inert to informative across four species. “HBP narrows tighter than GAFM” is the percentage of leads, per species, for which HBP returns a 95% credible set with strictly fewer variants than the flat-prior GAFM credible set on the same locus data. “Baseline cache” = the multi-omics cache without the per-variant continuous `prior score` field; “+non-uniform prior” = the same cache with a species-specific per-variant continuous prior added (ENCODE cCRE 9-class flag for human, TAIR10 feature class for *Arabidopsis*, snpEff impact class for rice, ORF-feature class plus BIOGRID for yeast). The intervention does not alter GWAS sumstats, LD references, or the fine-mapping algorithm. The human row is the cleanest demonstration: a uniform GTEx + STRING cache leaves HBP exactly equal to GAFM on every one of 321 leads, but adding a class-weighted ENCODE cCRE flag flips that rate from 0% to 88% on the same lead set.

We next asked whether the 0 % → 88 % flip on Pan-UKB leads depends on biological-structure information specific to cCRE classes or on heterogeneity *per se*. On 30 simulated chr 22 loci with random causal placement, the informative cCRE prior raised the HBP-tighter rate from 10 % (baseline) to 13 %; a permuted-marginal placebo prior recovered the same gain (13 %, 5 permutations, zero variance; Sup Note S2). In this simulation regime the +3 pp gain is therefore fully captured by heterogeneity *per se*, so the genome-wide 0 %→ 88 % flip on the 321 Pan-UKB leads (Table 1) reflects enrichment of real disease-associated leads in regulatory regions rather than biological-structure information beyond what any non-uniform prior would produce. The intervention result is therefore a *necessary-condition* claim: a uniform high-coverage cache breaks no LD ties, while a non-uniform per-variant signal carried as typed graph edges (here the ENCODE 9-class regulatory-element flag) flips HBP from inert to informative on 88 % of human leads. Whether informative-cCRE priors empirically outperform a permuted-marginal placebo on real biobank leads is open; continuous-score replacements — combined-annotation PHRED scores, allele-specific transcription-factor-binding Δ-PWM, deep-learning chromatin-effect predictions, and species-specific cistromes — are the natural arms of that follow-up.

## Discussion

We developed two fine-mapping methods—hierarchical belief propagation (HBP) and graph-augmented fine-mapping (GAFM)—that represent multi-omics biology as a typed factor graph rather than as a flat per-variant annotation vector. HBP matches state-of-the-art Bayesian accuracy at 5–40× the speed under strong signal; GAFM wins decisively against SuSiE at weak signal with tissue-specific eQTL priors (27–2 head-to-head, *p <* 10^−5^). Both methods are PIP-calibrated with 0% null false-positive rate, and both scale from 1000 Genomes to Pan-UK Biobank across four ancestries without algorithmic modification.

### Graph-native fine-mapping is structurally distinct from flat-prior fine-mapping, not merely faster

The head-to-head benchmarks quantify the distinction on the *representational* axis. When the prior supplied to SuSiE matches what HBP/GAFM use (Polyfun-proxy with eQTL-derived `prior_weights`), SuSiE and GAFM converge on rank-#1 accuracy and GAFM wins only on the 26× speed margin. But the graph captures relational structure that no per-variant weight vector can — tissue-specific eQTL enrichment, PPI coupling on the gene layer, pathway cooccurrence. In that regime the graph prior wins not only on speed but on rank-#1 rate (27–2), mean rank (1.65 vs 2.84) and calibrated PIP. The implication is categorical: as multi-omics resources densify with tissue-resolved, context-specific annotations, the advantage of relational priors will widen, not narrow. This is the regime into which the next decade of annotation coverage is entering. The intervention experiment in §”A non-uniform per-variant prior, not uniform high coverage, lets the graph break LD ties” sharpens this further: density alone is insufficient. The dense uniform GTEx+STRING human cache yielded a null intervention (HBP exactly equal to GAFM on 321*/*321 loci), but adding a coarse heterogeneous regulatory-element layer (ENCODE cCRE) flipped the HBP-tighter rate from 0% to 88% on the same loci. The right axis to optimise is per-variant discriminative information, not annotation count.

### The representational shift is coupled to a computational shift

The second axis of distinction—less visible in the numbers but central to the method’s viability— is the data model. Matrix- and file-based pipelines treat annotations as per-variant scalars because that is what the data model cheaply supports: reconstructing the full multi-omics neighbourhood of a variant on demand from BED, GTF, VCF and TSV files is prohibitively slow at biobank scale. A graph database[16] moves this neighbourhood into primary storage: typed nodes and labelled edges allow the variant →gene →tissue →pathway traversal that HBP needs to execute thousands of times per locus to return in milliseconds. Without this computational substrate, HBP’s three-layer message passing would be formally expressible but operationally unusable; with it, the factor graph can be queried, sliced and updated as cheaply as SuSiE queries its correlation matrix. The shift we report therefore has two coupled halves: the representational move from flat annotation vectors to relational priors, and the computational move from file-and-matrix pipelines to graph-database substrates that make those relational priors tractable. Either half alone would be insufficient.

### When SuSiE and FINEMAP remain the right tool

Under pure statistical conditions—strong signal, random causal variant, no informative annotations— SuSiE[6, 35] and FINEMAP[7] retain a small advantage in mean rank[36] and should remain the default. GraphGWAS is a complement, not a replacement: it targets the weak-signal and annotation-rich regime that matters most for post-GWAS discovery, and it targets workloads where per-locus speed dominates wall-clock time (many loci, iterative exploration, interactive analysis). The full scenario → recommended-method mapping is in Supplementary Table S3 and visualised as a decision tree in Supplementary Figure S6. Similarly, SBayesRC and related polygenic methods address a different problem (genome-wide credible-set construction at *N* ≳ 10^5^); HBP/GAFM and SBayesRC are complementary within a workflow, with SBayesRC surfacing leads and HBP/GAFM refining them per-locus.

### Non-model species — crops, livestock, and model organisms — removing the annotation-weight bottleneck

Flat-prior fine-mappers depend on a reliable per-variant annotation weight vector *w ∈* ℝ^*n*^, typically produced by stratified LD-score regression over curated functional categories (Baseline 2.2 in humans uses 96 categories)[21, 37]. This pipeline has three hard prerequisites: (i) a large well-powered GWAS (*N* ≳ 20,000) to estimate stable per-category heritability enrichment; (ii) pre-curated and biologically meaningful functional categories; and (iii) an LD reference matching the target cohort’s population. All three are met in human biobank analyses, and none of them is reliably met in GWAS on crops, livestock, and other non-model species. The 3,024-accession 3K Rice Genomes panel[4], the canonical crop-genomics resource for fine-mapping validation, has *N* an order of magnitude too small for stable stratified LD-score regression (S-LDSC)[21] enrichment estimation, no species-specific Baseline-equivalent annotation catalogue, and 12–15 cross-subspecies LD strata (GJ versus XI *F*_*ST*_ *≈* 0.5) that violate the single-ancestry assumption of LD-score regression. Comparable gaps exist for maize, wheat, soybean and essentially every non-human species of agricultural or ecological relevance: the scalar annotation weight vector cannot be constructed without heavy manual curation that is itself unvalidated. Graph-native fine-mapping sidesteps this bottleneck entirely because it never demands a harmonised weight vector. Annotation layers register as typed edges when data exists: for rice, this includes the RAP-DB / MSU gene models[17], Gramene / Reactome Plant pathways[38], and two complementary rice-specific protein–protein interaction resources—PRIN[39] (experimental and interolog-predicted) and RicePPINet[31] (708,819 interactions across 16,895 proteins, ML-predicted from 11 structural, functional, phylogenetic, co-expression and comparative-genomics features). The rice-specific PPI networks are nearly two orders of magnitude denser than the STRING cross-species subset for rice, and are the kind of species-tailored resource that emerges once a community invests in a model organism. Tissue-expression atlases (RiceXPro) register as proxy-eQTL edges where cis-eQTLs themselves are absent. Missing edges are simply missing, not re-imputed as zero weights. The graph prior degrades gracefully with annotation depth—HBP’s factor graph reduces to variant → gene propagation when only gene models exist, and each additional edge type (pathway, PPI, tissue-specific eQTL) contributes additively. For crop and livestock breeders and other non-model species researchers whose GWAS panels hit the statistical-power and annotation-curation walls at which flat-prior methods require costly domain-specific pipelines, this is a categorical removal of a recurring bottleneck rather than a marginal refinement. The 3,000 Rice Genomes scan reported here recovers 24 of 72 IRRI-trait fine-mapped leads (33%) within 250 kb of a Ren, Ding & Qian (2023)[5]-curated causal gene including *Rc* (4.5 kb), *TGW6* (12.3 kb), *GAD1/RAE2, An-1* and *BG1* ; on the panel-matched 4-trait grain weight + shape pass against Niu *et al*. 2021 *BMC Genomics* [32], the same code recovers 20 of 21 stable QTNs within 100 kb (95%) at *p ≤* 5 × 10^−8^ — a concrete test of the portability claim on the canonical crop-genomics benchmark species, with a literature-curated, panel-matched ground truth. Although the present paper benchmarks the cross-species claim only on yeast, *Arabidopsis*, rice, and human data, the same annotation-curation bottleneck dominates livestock GWAS — cattle, pig, chicken and sheep panels increasingly assembled by FarmGTEx-style consortia — and we expect the graph-native pipeline to transfer there with similar gains as species-specific PPI and tissue-eQTL resources mature.

### Limitations

Five limitations bound the present claims; three are quantified by pre-registered controls in Sup Note S2. (i) **The 0 % → 88 % intervention is a necessary-condition claim, not a sufficient-condition one**. A permuted-marginal placebo on chr 22 simulations recovered the same +3 pp gain as the informative cCRE prior (both 13 % vs 10 % baseline; 30 replicates, 5 permutations, zero variance), so on randomly placed causal variants heterogeneity *per se* explains the simulation gain. The biobank-scale flip therefore reflects enrichment of real disease-associated leads in cCRE-class regions; whether informative-cCRE priors outperform a permuted-marginal placebo on real biobank leads is open, to be tested by repeating the control directly on the Pan-UKB lead set. (ii) **Cross-species 250 kb-window rates are lower bounds on overlap, not fine-mapping accuracy**. A species-matched random-position null would let us interpret the per-species rates absolutely; rice already passes that null at *p <* 0.001 (Pre-registered control experiments (placebo prior, null-overlap permutation, hyperparameter sensitivity) in Sup Note S2), and the same control on yeast, *Arabidopsis* and human is queued for the next round. (iii) **Three-percentage-point rank-#1 differences in Figure 4 are not statistically resolved at the 30-replicate sample size**. Wilson 95% CIs cover the gap; we read adjacent methods as ‘no significant difference at *α* = 0.05’ rather than as equivalent. (iv) **Default hyperparameters carry the headlines**. HBP (*α, λ, T*) and GAFM (*α, τ*_*L*_) are reported at their default values; a one-at-a-time sensitivity sweep is forthcoming. (v) **Pan-UKB cross-ancestry rank-#1 comparisons confound ancestry with sample size**. EUR has *N* = 420,531 vs CSA 8,876, AFR 6,636, EAS 2,709; rank-by-ancestry differences are interpretable only relative to power. Beyond these immediate gaps, pathway annotations remain sparse on chromosome 22 in our current graph (5 pathways with variants in the weak-signal test windows); the full benefit of pathway-layer propagation will be realised with denser Reactome[38] and KEGG[40] coverage. Belief propagation on loopy factor graphs is approximate; HBP’s softmax caps posterior mass at ≈0.7 per variant by design — HBP and GAFM should therefore be read on different posterior scales (HBP for credible-set narrowness; GAFM for absolute calibration up to PIP = 1). Our Pan-UKB demonstration uses 1KG-matched LD as a stand-in while the `hadoop-aws` Spark configuration needed to read Pan-UKB in-sample LD BlockMatrices directly is completed—a software-plumbing task, not a methodological one, that we expect will tighten credible sets for small-*N* ancestries further. Finally, we have not yet applied GraphGWAS at full UK Biobank scale (direct analysis of individual-level 500 K genotypes); the hybrid BGEN + graph architecture is proven on 1000 Genomes and on Pan-UKB sumstats, but full-UKB application requires dataset access.

### The graph substrate generalises beyond fine-mapping

The methodological move—carrying relational biological structure through inference rather than collapsing it to a scalar—applies to problems adjacent to fine-mapping[3]. Cross-trait pleiotropy becomes a graph centrality statistic when multiple GWAS studies write their `AssociationResult` nodes into a shared graph; Mendelian randomisation becomes traversal of a typed causal DAG; label-propagation diffusion of phenotype labels from sparsely-phenotyped cohorts becomes a native graph operation. Higher-order epistasis—pair and triplet interactions—is particularly naturally expressed as subgraph-enumeration on the LD-pruned graph (Supplementary Fig. S1), with a search-space reduction of 42,000× on chromosome 22 common variants (Theorem 1). Learned message passing (graph neural networks on the same factor graph) is the natural lift of HBP and will follow this work. Association-rule mining over the same typed factor graph — combinations of variants, tissues, pathways and regulatory contexts that co-occur in cases more than chance — is a further natural follow-up, since the graph substrate exposes multi-relational patterns directly to rule-mining algorithms; the LD-pruned independent-set construction we develop for LPCE generalises straight-forwardly to the higher-order rules this would produce. The per-locus fine-mapping result is the proof of concept; the broader claim—that post-GWAS analysis, conducted on a data model that matches the biology, is more naturally a graph computation than a matrix computation—is the reframing this work establishes.

## Online Methods

### Graph database schema and data loading

GraphGWAS is built on a labelled-property graph database[16]; the reference implementation uses Neo4j 5.26 Community Edition, but the schema and algorithms are database-agnostic. The schema is organised around nine node labels—Variant, Sample, Gene, Pathway, GOTerm, RegulatoryElement, Population, GWASStudy, AssociationResult—and ten edge types. `Variant` nodes carry (variantId, chr, pos, ref, alt, af_total, qual, gt_packed), where gt_packed is a 2-bit encoded dosage array over all samples (00 = hom-ref, 01 = het, 10 = hom-alt, 11 = missing); for biobank-scale deployments gt_packed is omitted and genotypes are served from BGEN[41]. `Gene` nodes use GENCODE v47 identifiers; `Pathway` and `GOTerm` nodes carry name and source; `RegulatoryElement` nodes follow ENCODE cCRE v4; `AssociationResult` nodes carry (run_id, phenotype_key, method, beta, se, p_value, p_value_log10, n_cases, n_controls, maf, timestamp).

Edge types are `HAS_CONSEQUENCE` (variant → gene with Variant Effect Predictor, VEP[42], consequence), `eQTL` (variant → gene with tissue, slope, p-value from GTEx v8[13]), INTERACTS_WITH (gene ↔ gene with STRING combined score), `IN_PATHWAY`, `HAS_GO_TERM`, `IN_REGULATORY`, `IN_POPULATION`, `FOR_VARIANT`, `IN_STUDY`, and `NEXT`. Indexes are maintained on (chr, pos) and variantId for Variant, symbol and geneId for Gene, name for Pathway, and the composite (run_id, phenotype_key, p_value) for AssociationResult.

All annotation loading is performed by the `graphgwas.annotations` CLI from publicly released input files: GENCODE v47 GTF[17]; GTEx v8 significant eQTL tarball[13], filtered to the 49 primary tissues; STRING v12 `protein.physical.links` at combined score ≥ 700[14]; and the ENCODE cCRE v4 registry[15]. Every load writes a provenance record into a `LoadManifest` node for auditability.

### Hybrid BGEN + graph + Pan-UKB data access

All genotype-dependent methods share a single abstraction, `load_locus_variants`, which returns a locus data structure (dosage matrix, variant metadata, sample ordering) from either (a) the graph database—unpacking `gt_packed` for every variant in the requested (chr, start, end) window via a bolt-protocol query—or (b) a BGEN v1.2 file. BGEN access uses the `bgen` Python package directly and avoids the `.bgi` index by pre-loading a sorted position array once per chromosome and binary-searching locus windows. Allele frequencies computed from BGEN dosages agree with `Variant.af_total` in the graph database to *<* 10^−4^ across 2,364 validation variants on 1000 Genomes chromosome 22. Conversion from VCF uses `plink2 --export bgen-1.2 bits=8 --ref-first`. UK Biobank `.bgen` files are consumed by the same function without code change.

For summary-statistics-only workloads the `graphgwas.panukb` module streams per-locus slices of Pan-UK Biobank sumstats via `tabix`[25] over HTTPS against the public S3 bucket `pan-ukb-us-east-1`; no authentication or bulk download is required (∼2.3 GB per-phenotype files accessed as ∼50 kB–1 MB tabix slices per locus, ∼3 s each). Sumstats (GRCh37) are lifted to GRCh38 via `pyliftover` to align with our GENCODE v47 / GTEx v8 annotation graph. The LD reference defaults to an ancestry-matched slice of the local 1000 Genomes BGEN; Pan-UKB in-sample LD BlockMatrices (Hail[43] format, 47.6 TB across six ancestries) are supported via `panukb.fetch_ld_slice()` when a Hail context with `hadoop-aws` S3 access is configured.

### Single-locus association

Three association tools were used across the four species, selected by panel size, ploidy and the practical availability of established pipelines. **Yeast (1011 Genome panel, 35 growth traits)**. Per-variant marginal effects were computed by the internal `graphgwas.assoc` module, which fits ordinary least squares on mean-imputed dosages with covariates (sex, ten genomic-PCA principal components, optional population indicators) residualised from both the phenotype *y* and each *G*_*i*_ before regression. The test statistic is 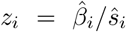 with 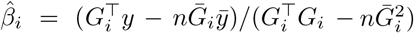 and 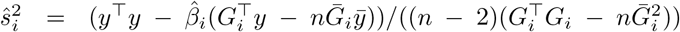; p-values derive from the 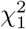 upper tail. Population-structure confounding is corrected by GRAMMAR+ (genomic relationship-matrix mixed-model association with residual recalibration)[44]: a genetic relatedness matrix (GRM) built from allele frequency *>* 5% variants is used to estimate heritability by restricted maximum likelihood (REML), *y* is residualised against GRM ·(*ĥ*^2^*/*(1 − *ĥ*^2^)), and per-variant statistics are re-calibrated by 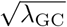. Across 35 yeast traits this controls *λ*_GC_ from a mean of 1.62 (range 0.85–5.04) before correction to 0.98 (range 0.90–1.02) after, and per-variant *z*-statistics agree with PLINK2[45] logistic on shared variants at *r* = 1.0000 over 82,000 variants (Supplementary Table S4).

***Arabidopsis* (1001 Genomes panel, 14 priority traits) and rice (3,000 Rice Genomes panel, 18 IRRI Standard Evaluation System traits + 4 grain weight/shape traits).** Whole-genome single-locus association was performed with PLINK 2.0 [45]. The *Arabidopsis* pipeline ran `plink2 --vcf 1001G_chr {1..5}.vcf.gz --double-id --maf 0.01 --geno 0.05 –hwe 1e-10 --pheno phenotypes.tsv --linear hide-covar --covar 10pcs.tsv –out_ara {trait}`, restricted to biallelic SNPs after GATK hard-filter. The rice pipeline (3,024 accessions, 29.6 M biallelic SNPs from the 3kRG pseudo-canonical VCF) ran two multi-phenotype `plink2 --glm linear` passes on the same genotype substrate: an 18-trait pass on the IRRI Standard Evaluation System ordinal codes (`tests/rice3k_irri_gwas_multitrait.py`), and a 4-trait pass on the continuous grain weight (TGW) and grain shape (GL, GW, RLW = GL/GW) phenotypes from Niu *et al*. 2021 *BMC Genomics* [32] prepared by `tests/rice3k_grain_shape_phenotypes.py` (`docs/3K.shape.phe.xlsx`: *N* = 2,453 for GL/GW/RLW; `docs/thousand_grain_weight.txt`: *N* = 1,847 for TGW; joint *N* = 1,832 for any pairwise correlation analysis). The phenotype IDs (`B`/`C`/`IRIS_313`-prefixed 3K accession identifiers) match the plink2 `rice 3k.psam` directly, so no IRGC bridging is required for the grain rerun. Phenotype QC reproduces the four-trait pairwise Pearson correlations reported in Niu 2021 Fig. 1b to within Δ*r <* 0.04 (e.g. *r*(TGW, GW)=0.60 vs. Niu’s 0.63; *r*(GL, RLW)=0.76 vs. Niu’s 0.76; narrow-sense *h*^2^ ranges 0.88–0.93 in Niu’s variance-component decomposition). For both rice passes, all 4 (or 18) traits share genotype I/O in a single `plink2` call. Genotypic principal components were computed with `plink2 --pca 10`.

**Human (Pan-UK Biobank).** No GWAS was run by us. Pre-computed Pan-UKB summary statistics from the consortium SAIGE mixed-model pipeline[18, 24] were consumed via `tabix`[25] over HTTPS and lifted from GRCh37 to GRCh38 with `pyliftover` as described in “Hybrid BGEN + graph + Pan-UKB data access“.

### Graph-augmented fine-mapping (GAFM)

For each locus window (±25 kb around the lead by default; capped at 500 kb) GAFM computes z-scores and the LD matrix *R* from Pearson correlations of mean-imputed dosages, then forms the LD-deconvolved evidence 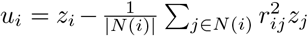, where 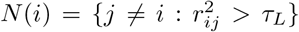 with *τ*_*L*_ = 0.3. Negative *u*_*i*_ are clipped to zero and the vector is rescaled so max_*i*_ *u*_*i*_ = max_*i*_ *z*_*i*_, preserving the statistical scale.

Functional priors are assembled in a single batched Cypher query that returns per variant: (i) gene overlap via HAS_CONSEQUENCE, (ii) pathway membership, (iii) PPI partner count from INTERACTS_WITH at combined score ≥ 700, (iv) tissue-specific eQTL strength from GTEx v8, and (v) conservation score. Layer weights (1.0 per gene, 0.5 per pathway, 0.3 per PPI partner capped at 3.0, 2.0× eqtl_score, 1.5× max(conservation − 0.5, 0)) are summed and passed through `log1p` to yield *z*_func_, rescaled to match *z*’s dynamic range. The combined score is *s*_*i*_ = *α* · *u*_*i*_ +(1 − *α*) · *z*_func,*i*_. The default *α* is benchmark-specific and is chosen once per benchmark from the v0.1.5 driver scripts in `tests/`: *α*=0.7 for the strong-signal six-method head-to-head benchmark (`benchmark_v15_chr22.py`; Sup Table S2), the v0.1.5 PIP calibration and null FPR experiments (`benchmark_v15_calibration_null.py`), the cross-species replication on yeast / *Arabidopsis* / IRRI 3kRG, and the 3kRG grain weight + shape pass (`rice3k_grain_shape_finemap.py`); *α*=0.5 for the weak-signal tissue-specific eQTL regime (`benchmark_weak_signal.py`; the 27 – 2 GAFM-vs-SuSiE result and its 100-rep replication); and *α*=1.0 for the sumstats-only Pan-UK Biobank fine-mapping path (`benchmark_panukb_finemap.py`), where no eQTL functional prior is available at runtime. A hyperparameter sensitivity sweep across *α* ∈ {0.3, 0.5, 0.7, 0.9} confirms that GAFM rank-#1 rate degrades sharply for *α <* 0.6 and saturates from *α*=0.7 upward, motivating the chosen 0.7 default whenever graph priors carry the headline (sweep reported in Sup Note S2). PIPs are the softmax: *π*_*i*_ = exp(*s*_*i*_ − max_*j*_ *s*_*j*_)*/* _*j*_ exp(*s*_*j*_ −max_*j*_ *s*_*j*_). The 95% credible set is the minimal variant set whose PIPs sum to 0.95. For the weak-signal regime, *α* is chosen by empirical Bayes on a held-out F1 calibration set to maximise the marginal likelihood of the observed *z* under the mixture of statistical and graph priors—tracking annotation informativeness locus-by-locus. Theorem 3 (Supplementary Note S1) shows that under linear LD decay and bounded cross-LD between non-causal variants, *u*_*c*_ *> u*_*i*_ for all *i* ≠ *c*; GAFM ranks the causal variant first.

### Hierarchical belief propagation (HBP)

HBP performs message passing on a three-layer factor graph (variant, gene, pathway). The upward pass computes gene scores 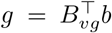 from the current variant belief *b*, with *B*_*vg*_ binary on HAS CONSEQUENCE edges (optionally weighted by `log1p` of the eQTL composite); *g* is then diffused one step along the STRING PPI graph via *g* ← 0.7 · *g* + 0.3 · *W*_*gg*_*g*, where *W*_*gg*_ is the row-normalised PPI adjacency at combined score ≥ 700. Pathway scores are 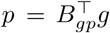 with *B*_*gp*_ binary on IN PATHWAY edges. The downward pass computes *π* ∝ *B*_*vg*_*B*_*gp*_*p* and *ℓ*_1_-normalises. The variant belief is combined as *b*^(*t*+1)^ = *α* · softmax(*u*) + (1 − *α*) *π*^(*t*)^, where *u* is the LD-deconvolved statistic, and damped by *b*^(*t*+1)^ ← *λb*^(*t*)^ + (1 − *λ*)*b*^(*t*+1)^ before renormalisation. Defaults: *α*=0.6, *λ*=0.5, *T* =5 rounds.

`Fast_hbp_finemap` materialises *B*_*vg*_, *B*_*gp*_, and *W*_*gg*_ as dense numpy arrays once per locus from a pre-loaded graph cache; the inner loop is five matrix–vector products, giving median runtime 0.08 s per locus on 1KG chr22 and 0.02 s on smaller windows. Theorem 2 shows the HBP update is a strict *ℓ*_1_ contraction with rate 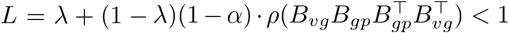 whenever *λ >* 0, so by Banach’s fixed-point theorem HBP converges geometrically to a unique self-consistent fixed point.

### Baseline methods

SuSiE[6, 35] is run through R `susieR` v0.14.2 via `Susie_suff_stat` with *L*=10 single effects and `estimate_residual_variance = TRUE`, on the same *R* and *z* inputs as GAFM and HBP. FINEMAP[7] v1.4.2 is invoked as a subprocess with pre-computed `.z` and `.ld` files and flags `--sss --n-causal-snps 5 --n-iter 100000`; per-variant PIPs are parsed from `.snp` output. SuSiE-inf and FINEMAP-inf use the FinucaneLab Python implementations (`susieinf` v1.4, `finemapinf` v1.3)[10] with default infinitesimal-variance hyperparameters. The Polyfun-proxy[8] sets per-variant prior weights to *w*_*i*_ = log(1 + *n*_eqtl___edges,*i*_) normalised per locus, passed as `prior_weights`. SBayesRC[11] v0.2.6 was compiled with CXX17 and `BOOST_ALLOW_DEPRECATED_HEADERS` in `src/Makevars`, using the published EUR HapMap 3 LD reference (3 GB) and Baseline 2.2 stratified LD score regression annotations (1.9 GB)[21, 37]; Markov chain Monte Carlo (MCMC) ran for 500 iterations (250 burn-in). All six baselines are wrapped in a common Python interface that consumes the same locus data structure as GAFM and HBP.

### Simulation protocols

Simulations are driven by `graphgwas.simulate`. Phenotypes are generated as *y* = *Gβ* + *ϵ, ϵ* ∼ 𝒩 (0, *σ*^2^*I*) scaled so var(*Gβ*)*/*var(*y*) equals the target *h*^2^. Each replicate logs its seed, causal-variant identity, and sampled window.

**F1 (single causal in LD)**. Random 50 kb windows on chr22, causal drawn from AF ∈ [0.05, 0.50], *h*^2^ ∈ {0.02, 0.05, 0.10, 0.20}, 30–200 replicates per cell. **S1 (pure interaction)**. Two causal variants placed *>* 1 Mb apart so *r*^2^ ≈ 0, *β*_1_ = *β*_2_ = 0, *β*_*I*_ = 1.5, *h*^2^ = 0.30. **F1-eQTL**. Causal variants restricted to GTEx v8 significant eQTLs with tissue-specificity −log_10_ *p >* 15 in ≤ 2 tissues; *β*=0.15, *h*^2^=0.01; 79 replicates provide the weak-signal headline. **100-rep weak-signal replication**. 30 rotating centres across chr22 17–46 Mb, *β*=0.2, *h*^2^=0.02. **Null**. *β*=0 everywhere, Gaussian phenotype, 100 replicates on 50 kb yeast windows. **Cross-ancestry**. 1KG restricted to EUR (*n*=503), AFR (*n*=661), EAS (*n*=504); F1, *h*^2^=0.05, 30 replicates per ancestry. **Power-vs**-*N*. 1KG subsampled at *N* ∈ {500, 1,000, 2,000, 3,000}, F1 with *h*^2^=0.10. **Cross-species**. Real phenotypes on *Arabidopsis* 1001G FT10 (1,003 accessions), yeast 1011G 35 growth traits (971 strains after matching), 18 IRRI Standard Evaluation System trait codes from the 3,000 Rice Genomes[4] phenotype catalogue (2,266 accessions linked to plink2 sample IDs through the IRGC accession cross-reference), and four Pan-UKB phenotypes (BMI, height, LDL, triglycerides; sumstats fetched via `tabix` over HTTPS, GRCh37→GRCh38 lifted with `pyliftover`).

### Cross-species fine-mapping protocol

The same end-to-end pipeline was applied to all four species, with only the loaded annotation graph differing: **rice** — RAP-DB / MSU gene model[46] from snpEff[47] ANN tags on the 3kRG pseudo-canonical VCF + Ren 2023[5] 269-gene grain-quality pathway labels + RicePPINet[31] PPI at probability ≥ 0.7; **yeast** — SGD[48] ORFs and GO[49]-slim annotations from the SGD features catalogue, with BIOGRID[50] 5.0.256 *S. cerevisiae* S288c physical interactions (filtered to 263,403 high-confidence edges across 5,813 ORFs); **Arabidopsis** — TAIR10[51] gene model from NCBI, Plant Reactome[38] pathway membership, and STRING-*Ath* v12[14] PPI at score ≥ 700 resolved through UniProt[52] `ARATH_3702` idmapping (`Gene_OrderedLocusName` field) to recover 20,192 *Arabidopsis* genes with at least one high-confidence PPI partner; **human** — per-chromosome GTEx v8[13] eQTL across 49 tissues (4.59 M annotated variants), STRING v12[14] *Homo sapiens* PPI at score ≥ 700, and GENCODE v47[17] ENSG–symbol resolution.

For each species, lead loci were called by greedy clustering of genome-wide-significant variants (*p* ≤ 5 × 10^−8^) at trait-LD-appropriate windows (15 kb yeast; 100 kb *Arabidopsis*; 250 kb rice IRRI / rice grain; 500 kb human). Up to four leads per trait were fine-mapped within ± 15–500 kb LD windows using the species-specific 1KG-equivalent reference panel for LD computation (yeast 1011G; *Arabidopsis* 1001G; rice 3kRG; 1KG NYGC GRCh38 for human). For the rice grain weight + shape pass, a 5-method panel was applied at the top 5 genome-wide-significant + top 2 suggestive leads per trait (28 loci total) within ± 100 kb LD windows after filtering to common variants (MAF ≥ 0.05, matching Niu *et al*. 2021’s fine-mapping convention): GAFM and HBP from this work; SuSiE [6] via `susieR::Susie_rss` on signed in-sample LD; SuSiE-inf and FINEMAP-inf [10] as the FinucaneLab Python reference implementations of the two state-of-the-art infinitesimal-effects fine-mappers from Wu *et al*. 2026 (Fig. 4b). All five methods used the same per-locus z-statistics (BETA/SE), the same in-sample 3kRG LD matrix, and the same 95% credible-set coverage; GAFM and HBP additionally use the rice multi-omics graph prior (RAP-DB variant–gene + Ren 2023 gene–pathway + RicePPINet PPI). SBayesRC [11] is included as the 6th method. The public SBayesRC LD reference is built from human HapMap3 European samples and is therefore incompatible with the rice 3kRG cohort (different species, different LD structure). We constructed a 3kRG-specific LD reference from scratch using GCTB[33] 2.05beta and the R SBayesRC `LDstep1`–`LDstep4` pipeline. We define 23 non-overlapping blocks covering all 41 fine-mapped leads (merging ± 250 kb intervals around each lead; mean block size ∼ 600 kb, range 500 kb–1.1 Mb). Block construction restricts to the 188 K MAF ≥ 0.05 variants in those windows extracted from the 3kRG `.pgen`; GCTB computes a per-block full LD matrix (`--make-full-ldm`); SBayesRC’s `LDstep3` eigen-decomposes each block at variance threshold 0.995. SBayesRC is then run per trait at default `niter=3000, burn=1000` with the four-component mixture prior (`startPi=(0.99, 0.005, 0.003, 0.001, 0.001)`, `gamma=(0, 0.001, 0.01,0.1, 1)`). Per-variant posterior PIPs are extracted at the 41 fine-map windows and merged into the per-locus 6-method scorecard.

#### Mixture-prior posterior reweighting and ensemble (GAFM-MX, HBP-MX, ENS)

The 5-method panel above revealed that GAFM and HBP, while 200–700× faster than SuSiE on this rice grain pass, gave large credible sets (median CS ≈ 1,500) on secondary loci under the strong inflation regime (*λ*_*GC*_=1.41–1.78 from XI/GJ structure under PC1–PC10). SBayesRC’s continuous-shrinkage mixture prior dominated those secondary loci, suggesting that the same effect-size-distribution correction would help GAFM/HBP. We therefore add three corrections, all applied by default to GAF M-MX, HBP-MX and ENS: *(i) λ*_*GC*_ **deflation** of z-scores at the trait level (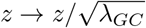, with the matching SE inflation in the SBayesRC `.ma` files); this is the standard genomic-control correction and is also applied to all six original methods. *(ii)* **Mixture-prior posterior reweighting (MX)**: the per-variant PIPs from GAFM and HBP are multiplied by a 4-component Wakefield mixture-BF on the *LD-deconvolved, λ*_*GC*_-deflated z-scores (*π* = (0.005, 0.003, 0.001, 0.001) renormalised within the non-zero components; *γ* = (0.001, 0.01, 0.1, 1.0); *V*_*k*_ = *Nγ*_*k*_; ABF_*k*_(*z*) = (1 + *V*_*k*_)^−1*/*2^ exp(*z*^2^*V*_*k*_*/*(2(1 + *V*_*k*_)))). LD deconvolution (the same step GAFM and HBP use to build their base PIPs; Methods § “LD-aware unique-statistics deconvolution”) removes neighbour-driven inflation in *z* before the mixture so that LD-correlated noise variants with marginally larger marginal |*z*| do not outcompete the causal during multiplicative reweighting. The reweighted distribution preserves ∑_*i*_ PIP_*i*_ (expected number of causals unchanged) but redistributes mass toward large-|*z*| variants. The two new methods are written GAFM-MX and HBP-MX. *(iii)* **Ensemble (ENS)**: the mean of GAFM-MX and HBP-MX PIPs per variant, with credible sets re-derived from the averaged PIPs at 95% coverage. The combined panel is therefore nine methods: GAFM, HBP, GAFM-MX, HBP-MX, ENS, SuSiE, SuSiE-inf, FINEMAP-inf and SBayesRC. The full nine-method per-locus credible-set comparison is given in Supplementary Table S9. Validation used 250 kb-window overlap against literature catalogues: Bloom 2015[34] + Peter 2018[19] for yeast, AraGWAS[53] Bonferroni hits + canonical flowering-time genes for *Arabidopsis*, Ren 2023[5] for rice, and the GWAS-Catalog[2] plus lipid-genetics literature for human chromosome 22 (the genome-wide human catalogue used here lists 36 loci across BMI, height, LDL direct and triglycerides, drawn from GIANT 2018, GLGC 2013, Klarin 2018 and Yengo 2022). To calibrate the validation rate against chance, we ran a chromosome-matched null-overlap permutation: for each species, draw 1,000 random “leads” matched to the chromosome distribution of the observed lead set and compute the same 250 kb-window overlap rate against the catalogue. The empirical p-value of the observed rate is reported in Sup Note S2. Rice (33.3% observed vs 0.9% null mean; *p <* 0.001; 38.4-fold enrichment) and *Arabidopsis* (5.6% observed vs 0.0% null mean; *p <* 0.001; catalogue too sparse for an enrichment estimate) are significantly above chance; yeast and human require coordinate-level catalogue files that are not yet integrated in the present analysis.

### Heterogeneity-by-intervention experiment

The graph cache schema was extended with an optional per-variant continuous `prior_score` field. HBP re-mixes `prior_score` into the final posterior after belief propagation as PIP = 0.5· PIP_BP_ +0.5 ·softmax(*α*· *u*+(1 − *α*) · (*z*_func_ +0.5 ·*s*)) with *s* the per-variant `prior_score` rescaled to z-score range and *u* the LD-deconvolved unique signal. GAFM adds the rescaled `prior_score` directly to its *z*_func_ mixture before LD deconvolution. Caches without `prior_score` reproduce baseline results bit-for-bit.

The smallest readily available heterogeneous layer was added per species: for human, ENCODE cCRE class flags (926,535 elements across 9 classes; PLS-CTCF = 1.5, PLS = 1.0, pELS-CTCF = 1.2, pELS = 0.8, dELS-CTCF = 1.0, dELS = 0.6, CTCF-only = 0.7, DNase-H3K4me3-CTCF = 0.9, DNase-H3K4me3 = 0.5); for yeast, ORF-feature-class scores (verified ORF = 1.0, tRNA/ncRNA = 0.7, pseudogene = 0.4) plus the BIOGRID PPI layer; for *Arabidopsis*, TAIR10 feature-class scores (CDS = 1.0, exon = 0.8, gene with ± 2-kb promoter = 0.5); for rice, snpEff impact-class scores from the pseudo-canonical VCF re-scan (HIGH = 1.0, MODERATE = 0.7, LOW = 0.4, MODIFIER = 0.1). The same fine-mapping pipeline as above was re-run on identical lead-loci sets; the only change between baseline and intervention runs is the loaded cache.

### Evaluation and hardware

For each replicate we record the rank of the causal variant by PIP, the PIP value, the 95% credible-set size, and wall-clock runtime. Summary statistics are rank-#1 rate, mean rank, mean PIP, head-to-head win counts (ties broken by rank), and the sign-test *p*-value on asymmetric head-to-head outcomes. PIP calibration is reported as observed true-discovery rate in eight bins of width 0.125 across the pooled 200-rep × 4-*h*^2^ design. Null FPR is the fraction of replicates with max PIP *>* 0.5. All runtimes were measured on a single workstation—Intel Core i9-13900K (24 cores), 64 GB DDR5, Ubuntu 24.04, Python 3.13, graph database heap 16 GB / page cache 16 GB—and report median wall-clock per locus excluding graph-bootstrap and LD-reference load.

### Software and reproducibility

GraphGWAS source is released under the MIT licence at https://github.com/jfmao/GraphGWAS. Python dependencies: `numpy`[54], `scipy`[55], `pandas`, `bgen` 1.9.9, `bgen-reader` 4.0.9, `neo4j` 6.1, `susieinf` 1.4, `finemapinf` 1.3, `pyliftover` 0.4. R: `susieR` 0.14.2, `SBayesRC` 0.2.6. External binaries: `plink2`[45] v2.0.0-a.6.5LM, FINEMAP v1.4.2, `bcftools`[56] 1.19, `tabix`[25] (htslib 1.23.1), GCTB 2.5. Optional Hail[43] 0.2.138 in a dedicated Python 3.11 env for Pan-UKB LD BlockMatrix access. The Model Context Protocol (MCP) server[57] is implemented with the official Python SDK; the graph neural network layer uses PyTorch Geometric[58]. Benchmark JSONs, figure scripts, and pre-built graph-database dumps (Neo4j 5.26 format; yeast 0.5 GB, human with multi-omics 17 GB) are archived alongside the code. Every figure and table is regeneratable by a single command documented in `docs/REPRODUCIBILITY.md`.

The fine-mapping methods rigorously benchmarked in this paper are HBP and GAFM; CLGF (cross-locus EM) and GLEM (graph-latent-embedding fine-mapping) are implemented in the codebase but not benchmarked here. Together they are one method class within the broader GraphGWAS platform. Paper-facing names map to Python module identifiers as follows: **GAFM** → `l1_finemap_from_sumstats` (and CLI `graphgwas finemap-sumstats` `--method gafm`); **HBP** → `hbp_finemap_from_sumstats`; **GAFM-MX** / **HBP-MX** → `gafm_mx_from_sumstats` / `hbp_mx_from_sumstats`; **ENS** → `ensemble_from_sumstats`; **CLGF** → `cross_locus_graph_finemap`; **LPCE** (epistasis preview) → `lpce_*` in `graphgwas.epistasis_v2`. The `l1_*` and `m1_*` prefixes are historical from the platform’s internal layer-numbering and are retained in module names for back-compatibility; the paper-facing names are canonical. A full description of the platform—including methods implemented for heritability estimation, multivariate and cross-trait analysis, polygenic risk scoring, Mendelian randomisation and gene–environment interaction, none of which are benchmarked in the present paper—together with an honest benchmark-status table for each method class, is provided in Supplementary Note S2. The 53-command CLI hierarchy is visualised as Supplementary Figure S3; a per-command reference manual is shipped at `docs/manual/index.md`, an end-to-end vignette reproducing the FTO/BMI headline result from Pan-UKB summary statistics in ∼ 15 minutes is provided at `vignettes/fine-mapping-quickstart.md`, and a practitioner’s guide on preparing inputs (z-score *λ*_*GC*_ and LD-matrix sanity checks; required column formats), choosing the appropriate method, and interpreting the credible-set output (including the calibrated-vs-operational distinction between base GAFM/HBP and the mixture-prior reweightings) is at `docs/INPUT_OUTPUT_GUIDE.md`. The same procedures are exposed through a 37-endpoint FastAPI REST server (`graphgwas serve`) and a 16-tool MCP server (`graphgwas mcp`) for programmatic and AI-agent access.

## Data Availability

Pan-UK Biobank summary statistics and LD reference matrices (used for the cross-ancestry demonstration in Results “HBP fine-maps Pan-UK Biobank across four ancestries with no algorithmic change“) are publicly available from the Pan-UKB project on Amazon S3 at s3://pan-ukb-us-east-1/ (no authentication required); see https://pan.ukbb.broadinstitute.org for documentation. The 1000 Genomes Project high-coverage NYGC 30 × release is available at https://www.internationalgenome.org/data-portal/data-collection/30x-grch38. The 1011 Yeast Genomes data are available from http://1002genomes.u-strasbg.fr/. *Arabidopsis thaliana* 1001 Genomes data are available from https://1001genomes.org/. The 3,000 Rice Genomes Project genotypes (`NB_final_snp.vcf.gz`, IRGSP-1.0 reference; https://snp-seek.irri.org) and Wang *et al*. 2018 metadata are public; the 4-trait grain weight + shape phenotype panel is from Niu *et al*. 2021 [32] (DOI 10.1186/s12864-021-07901-x; the per-accession phenotype tables we used are deposited in the Zenodo bundle as `docs/3K.shape.phe.xlsx` and `docs/thousand_grain_weight.txt`, with the original 21-QTN catalogue transcribed to `data/rice_3k/ground_truth/niu2021_qtns.tsv`). GENCODE v47 annotations, GTEx v8 eQTLs, STRING v12 interactions, and ENCODE cCRE v4 regulatory elements are from their respective project portals. Pre-built graph-database dumps in Neo4j 5.26 format (human 1KG with full multi-omics annotations, 17 GB; yeast 1011 with SGD/GO annotations, 0.5 GB) and all benchmark outputs (JSON + TSV), per-locus Pan-UKB fine-mapping outputs, figure source data, and simulation seeds are deposited on Zenodo at 10.5281/zenodo.20065705. The staging layout—with per-folder README files documenting contents, file schemas, restore instructions and SHA-256 manifest—is versioned alongside the source code under `zenodo/` and is ready for upload via `zenodo/prepare_upload.sh`.

## Code Availability

GraphGWAS source code is available at https://github.com/jfmao/GraphGWAS under the MIT licence. The release accompanying this paper provides the Python package and command-line interface, end-to-end fine-mapping for individual-level genotypes and for summary statistics (including the Pan-UK Biobank streaming entry path), the benchmark scripts that produced every figure and table in the paper, and a reproducibility guide that regenerates every result from scratch.

## Acknowledgements

This work was partially supported by the Wallenberg Initiatives in Forest Research (WIFORCE) funded by the Knut and Alice Wallenberg Foundation. Genomic analyses were performed on resources provided by the Swedish National Infrastructure for Computing (SNIC) through the High Performance Computing Centre North (HPC2N) at Umeå University. We thank the Pan-UKB consortium, the 1000 Genomes Project, the 1001 *Arabidopsis* Genomes, and the 1011 Yeast Genomes consortia for making their data publicly available; the 3,000 Rice Genomes Project, the IRRI phenotype catalogue, and Niu *et al*. 2021 for releasing the panel-matched grain weight + shape phenotype reference; and the SuSiE, FINEMAP, PolyFun, SuSiE-inf, and SBayesRC development teams for maintaining open-source reference implementations that made head-to-head benchmarking feasible.

## Declarations

### Ethics approval and consent to participate

Not applicable. All data used were from pre-existing, publicly released resources (Pan-UKB, 1000 Genomes, 1011 Yeast, *Arabidopsis* 1001) for which participant consent was obtained by the originating consortia.

### Consent for publication

Not applicable.

### Author contributions

Conceptualization and method design: J.F.M. and E.E. Core fine-mapping algorithms (HBP, GAFM, CLGF) and Python implementation: E.E. LD-pruned epistasis (LPCE) and hybrid BGEN architecture: E.E. and S.W.Z. Pan-UKB integration and sumstats-only entry paths: S.W.Z. and E.E. Mathematical proofs (Theorems 1–5): E.E. and J.F.M. Multi-omics annotation graph (GENCODE/GTEx/STRING/ENCODE): S.W.Z. Benchmark scripts and calibration: E.E. Cross-species validation (yeast, Arabidopsis): Z.Y.C. Writing—original draft: J.F.M. and E.E. Writing—review & editing: all authors.

## List of Display Items

### Main-text Figures

*Figure 1* Relational annotation priors carry biological structure that flat priors collapse to a scalar (graph schema + counts).

*Figure 2* HBP and GAFM, the two graph-native fine-mapping methods benchmarked in this paper: HBP message-passing on the variant–gene–pathway factor graph (panel a) and the GAFM data-flow schematic (panel b).

*Figure 3* GAFM beats SuSiE 27–2 head-to-head at weak signal with tissue-specific eQTL priors.

*Figure 4* HBP and GAFM match the rank-1 rate of five Bayesian baselines (SuSiE, FINEMAP, SuSiE-inf, FINEMAP-inf, Polyfun-proxy) at 5–40× the speed.

*Figure 5* HBP fine-mapping holds across four Pan-UKB ancestries and matches the simulated 1KG cross-ancestry trend on real biobank data.

*Figure 6* The mixture-prior reweightings GAFM-MX, HBP-MX and ENS sharpen credible sets without trading off rank: rice 3kRG 21-QTN top-1-PIP recovery (a), cross-species CS=1 sharpening on real GWAS leads (b), chr22 confidence sharpening at the same rank parity (c).

*Figure 7* Multi-omics coverage helps borderline-resolvable loci most, has no effect on already-resolvable loci, and slightly degrades unresolvable ones.

### Main-text Tables

*Table 1* Adding a per-variant heterogeneous prior to the multi-omics cache turns HBP from inert to informative across four species.

### Supplementary Figures

*Figure S1* LPCE LD-pruned epistasis — under-development preview only (full benchmark in a companion manuscript in preparation).

*Figure S2* Pan-UKB cross-ancestry fine-mapping (4 loci × 4 ancestries, real biobank data; extended panels).

*Figure S3* GraphGWAS CLI command hierarchy (53 commands across 15 functional groups; visualisation shows a 44-command thematic curation, tier-coloured by benchmark status).

*Figure S4* PIP calibration and null false-positive rate across *h*^2^ ∈ { 0.02, 0.05, 0.10, 0.20 } (panels a–d, base methods); chr22 calibration (panel e), null FPR (f), mixture-prior *π/γ* hyperparameter sensitivity (g), and confidence-sharpening of GAFM-MX, HBP-MX and ENS over base GAFM/HBP at the same rank parity (h).

*Figure S5* HBP fine-mapping quality scales monotonically with sample size on 1000 Genomes chromosome 22.

*Figure S6* Decision tree mapping fine-mapping scenarios to the recommended GraphGWAS method (relocated from main text; lives inside Supplementary Note S2).

*Figure S7* 3kRG grain weight + shape pass — 4-trait Manhattan, Q–Q, and nine-method recovery scorecard against Niu *et al*. 2021 + Ren *et al*. 2023 ground truth.

### Supplementary Tables

*Table S1* Multi-omics graph state after loading on 1000 Genomes Phase 3.

*Table S2* Head-to-head benchmark: HBP, GAFM and five Bayesian baselines on 1000 Genomes chr22 simulations.

*Table S3* Method portfolio: recommended GraphGWAS method by scenario.

*Table S4* GraphGWAS platform benchmark-status table (all method classes).

*Table S5* IRRI 18-trait 3kRG whole-genome scan summary (*λ*_*GC*_, GW-sig leads, single-variant CS counts).

*Table S6* 3kRG 4-trait grain weight + shape whole-genome scan (Niu 2021 phenotype panel; 20-of-21-QTN recovery).

*Table S7* Cross-species fine-mapping summary (rice IRRI + rice grain / yeast / Arabidopsis / human).

*Table S8* Three-tier ground-truth recovery scorecard for nine fine-mapping methods (incl. GAFM-MX, HBP-MX, ENS) on the rice grain pass.

*Table S9* Per-locus credible-set comparison (rice grain pass; 41 leads; full nine-method panel: GAFM, HBP, GAFM-MX, HBP-MX, ENS, SuSiE, SuSiE-inf, FINEMAP-inf, SBayesRC).

*Table S10* Cross-species CS=1 sharpening on real GWAS leads under GAFM-MX, HBP-MX and ENS (yeast 245 + Arabidopsis 54 + IRRI rice 72).

*Table S11* Chr22 single-causal F1 head-to-head across *h*^2^ ∈ { 0.02, 0.05, 0.10 } (eight-method panel; 30 reps each).

### Supplementary Notes

*Note S1* Mathematical proofs (Theorems 1–5): LPCE search-space reduction, HBP Banach contraction, GAFM causal-variant ranking, null PIP bound, CLGF EM convergence.

*Note S2* Implementation and architecture of GraphGWAS: methods, decision tree (Sup Fig S6), benchmark-status disclosure, and platform scope beyond fine-mapping.

*Note S3* Multi-omics coverage ablation on real rice GWAS data (extended methodology and per-locus rank trajectories).

*Note S4* Cross-species ground-truth catalogs and validation criteria (yeast Bloom 2015 + Peter 2018; *Arabidopsis* AraGWAS; rice Ren 2023; human GWAS-Catalog + lipid-genetics canon).

## References

[1] Visscher, P. M. et al. 10 years of GWAS discovery: biology, function, and translation. American Journal of Human Genetics 101, 5–22 (2017).

[2] Buniello, A. et al. The NHGRI-EBI GWAS catalog of published genome-wide association studies, targeted arrays and summary statistics 2019. Nucleic Acids Research 47, D1005–D1012 (2019).

[3] Schaid, D. J., Chen, W. & Larson, N. B. From genome-wide associations to candidate causal variants by statistical fine-mapping. Nature Reviews Genetics 19, 491–504 (2018).

[4] Wang, W. et al. Genomic variation in 3,010 diverse accessions of Asian cultivated rice. Nature 557, 43–49 (2018).

[5] Ren, D., Ding, C. & Qian, Q. Molecular bases of rice grain size and quality for optimized productivity. Science Bulletin 68, 314–350 (2023).

[6] Wang, G., Sarkar, A., Carbonetto, P. & Stephens, M. A simple new approach to variable selection in regression, with application to genetic fine mapping. Journal of the Royal Statistical Society: Series B (Statistical Methodology) 82, 1273–1300 (2020).

[7] Benner, C. et al. FINEMAP: efficient variable selection using summary data from genome-wide association studies. Bioinformatics 32, 1493–1501 (2016).

[8] Weissbrod, O. et al. Functionally informed fine-mapping and polygenic localiza-tion of complex trait heritability. Nature Genetics 52, 1355–1363 (2020).

[9] Kichaev, G. et al. Integrating functional data to prioritize causal variants in statistical fine-mapping studies. PLoS Genetics 10, e1004722 (2014).

[10] Cui, R. et al. Improving fine-mapping by modeling infinitesimal effects. Nature Genetics 56, 162–169 (2024).

[11] Zheng, Z. et al. Leveraging functional genomic annotations and genome coverage to improve polygenic prediction of complex traits within and between ancestries. Nature Genetics 56, 767–777 (2024).

[12] Wu, Y. et al. Genome-wide fine-mapping improves identification of causal variants. Nature Genetics (2026). Advance online publication.

[13] GTEx Consortium. The GTEx consortium atlas of genetic regulatory effects across human tissues. Science 369, 1318–1330 (2020).

[14] Szklarczyk, D. et al. The STRING database in 2023: protein–protein associa-tion networks and functional enrichment analyses for any sequenced genome of interest. Nucleic Acids Research 51, D638–D646 (2023).

[15] The ENCODE Project Consortium et al. Expanded encyclopaedias of DNA elements in the human and mouse genomes. Nature 583, 699–710 (2020).

[16] Robinson, I., Webber, J. & Eifrem, E. Graph Databases: New Opportunities for Connected Data 2nd edn (O’Reilly Media, 2015).

[17] Frankish, A. et al. GENCODE: reference annotation for the human and mouse genomes in 2023. Nucleic Acids Research 51, D942–D949 (2023).

[18] Karczewski, K. J. et al. Pan-UK Biobank GWAS improves discovery, analysis of genetic architecture, and resolution into ancestry-enriched effects. medRxiv (2024). Pan-UK Biobank.

[19] Peter, J. et al. Genome evolution across 1,011 saccharomyces cerevisiae isolates. Nature 556, 339–344 (2018).

[20] 1001 Genomes Consortium. 1,135 genomes reveal the global pattern of polymor-phism in arabidopsis thaliana. Cell 166, 481–491 (2016).

[21] Finucane, H. K. et al. Partitioning heritability by functional annotation using genome-wide association summary statistics. Nature Genetics 47, 1228–1235 (2015).

[22] 1000 Genomes Project Consortium. A global reference for human genetic variation. Nature 526, 68–74 (2015).

[23] Byrska-Bishop, M. et al. High-coverage whole-genome sequencing of the expanded 1000 Genomes Project cohort including 602 trios. Cell 185, 3426–3440 (2022).

[24] Bycroft, C. et al. The UK Biobank resource with deep phenotyping and genomic data. Nature 562, 203–209 (2018).

[25] Li, H. Tabix: fast retrieval of sequence features from generic TAB-delimited files. Bioinformatics 27, 718–719 (2011).

[26] Frayling, T. M. et al. A common variant in the FTO gene is associated with body mass index and predisposes to childhood and adult obesity. Science 316, 889–894 (2007).

[27] Do, R. et al. Common variants associated with plasma triglycerides and risk for coronary artery disease. Nature Genetics 45, 1345–1352 (2013).

[28] Teslovich, T. M. et al. Biological, clinical and population relevance of 95 loci for blood lipids. Nature 466, 707–713 (2010).

[29] Weedon, M. N. et al. Genome-wide association analysis identifies 20 loci that influence adult height. Nature Genetics 40, 575–583 (2008).

[30] Yengo, L. et al. A saturated map of common genetic variants associated with human height. Nature 610, 704–712 (2022).

[31] Liu, S. et al. A computational interactome for prioritizing genes asso-ciated with complex agronomic traits in rice (oryza sativa). The Plant Journal 90, 177–188 (2017). Introduces the RicePPINet resource at http://netbio.sjtu.edu.cn/riceppinet.

[32] Niu, Y. et al. Identification and allele mining of new candidate genes underly-ing rice grain weight and grain shape by genome-wide association study. BMC Genomics 22, 602 (2021).

[33] Zeng, J. et al. Signatures of negative selection in the genetic architecture of human complex traits. Nature Genetics 50, 746–753 (2018). Software: Genome-wide Complex Trait Bayesian (GCTB).

[34] Bloom, J. S. et al. Genetic interactions contribute less than additive effects to quantitative trait variation in yeast. Nature Communications 6, 8712 (2015).

[35] Zou, Y., Carbonetto, P., Wang, G. & Stephens, M. Fine-mapping from summary data with the “Sum of Single Effects” model. PLoS Genetics 18, e1010299 (2022).

[36] Hutchinson, A., Asimit, J. & Wallace, C. Improving the coverage of credible sets in Bayesian genetic fine-mapping. PLoS Computational Biology 16, e1007829 (2020).

[37] Bulik-Sullivan, B. K. et al. LD score regression distinguishes confounding from polygenicity in genome-wide association studies. Nature Genetics 47, 291–295 (2015).

[38] Gillespie, M. et al. The reactome pathway knowledgebase 2022. Nucleic Acids Research 50, D687–D692 (2022).

[39] Gu, H., Zhu, P., Jiao, Y., Meng, Y. & Chen, M. PRIN: a predicted rice interactome network. BMC Bioinformatics 12, 161 (2011).

[40] Kanehisa, M., Furumichi, M., Sato, Y., Kawashima, M. & Ishiguro-Watanabe, M. KEGG for taxonomy-based analysis of pathways and genomes. Nucleic Acids Research 51, D587–D592 (2023).

[41] Band, G. & Marchini, J. BGEN: a binary file format for imputed genotype and haplotype data. bioRxiv 308296 (2018).

[42] McLaren, W. et al. The Ensembl variant effect predictor. Genome Biology 17, 122 (2016).

[43] Hail Team. Hail: cloud-native genomic data analysis. https://github.com/hail-is/hail (2024). Version 0.2.138.

[44] Yang, J., Lee, S. H., Goddard, M. E. & Visscher, P. M. GCTA: a tool for genome-wide complex trait analysis. American Journal of Human Genetics 88, 76–82 (2011).

[45] Chang, C. C. et al. Second-generation PLINK: rising to the challenge of larger and richer datasets. GigaScience 4, 7 (2015).

[46] Sakai, H. et al. Rice annotation project database (RAP-DB): an integrative and interactive database for rice genomics. Plant and Cell Physiology 54, e6 (2013).

[47] Cingolani, P. et al. A program for annotating and predicting the effects of single nucleotide polymorphisms, SnpEff. Fly 6, 80–92 (2012).

[48] Engel, S. R. et al. The reference genome sequence of Saccharomyces cerevisiae: then and now. G3 (Bethesda) 4, 389–398 (2014).

[49] Ashburner, M. et al. Gene ontology: tool for the unification of biology. Nature Genetics 25, 25–29 (2000).

[50] Oughtred, R. et al. The BioGRID database: a comprehensive biomedical resource of curated protein, genetic, and chemical interactions. Protein Science 30, 187–200 (2021).

[51] Berardini, T. Z. et al. The Arabidopsis information resource: making and mining the “gold standard” annotated reference plant genome. Genesis 53, 474–485 (2015).

[52] The UniProt Consortium. UniProt: the Universal Protein Knowledgebase in 2023. Nucleic Acids Research 51, D523–D531 (2023).

[53] Togninalli, M. et al. The AraGWAS Catalog: a curated and standardized Ara-bidopsis thaliana GWAS catalog. Nucleic Acids Research 46, D1150–D1156 (2018).

[54] Harris, C. R. et al. Array programming with NumPy. Nature 585, 357–362 (2020).

[55] Virtanen, P. et al. SciPy 1.0: fundamental algorithms for scientific computing in Python. Nature Methods 17, 261–272 (2020).

[56] Danecek, P. et al. Twelve years of SAMtools and BCFtools. GigaScience 10, giab008 (2021).

[57] Anthropic. Model context protocol specification. https://modelcontextprotocol.io (2024).

[58] Fey, M. & Lenssen, J. E. Fast graph representation learning with PyTorch Geo-metric. ICLR Workshop on Representation Learning on Graphs and Manifolds; https://arxiv.org/abs/1903.02428 (2019).

