## supplementary_materials for "Relational biological structure improves fine-mapping of causal GWAS variants under weak signal"

### Supplementary Information

*Note on file references.* Throughout this Supplementary Information, paths in `typewriter` font (e.g. `tests/...`, `docs/...`, `vignettes/...`, `src/...`) refer to files in the GraphGWAS source repository at https://github.com/jfmao/GraphGWAS; the same paths are preserved in the Zenodo software-release archive cited under Code Availability. Per-experiment regenerator commands quoted in captions and notes assume the reader has cloned that repository.

### Supplementary Table S1 — Multi-omics graph state after loading on 1000 Genomes Phase 3

| Node / edge | Count |
|---|---:|
| Variant | 70,691,875 |
| Sample | 3,202 |
| Gene (GENCODE v47) | 20,092 |
| GTEx v8 tissue eQTL | 43,170,154 |
| `HAS_CONSEQUENCE` edges | 38,940,085 |
| STRING `INTERACTS_WITH` ($\geq$ 700) | 230,850 |
| RegulatoryElement (ENCODE cCRE) | 370,000 |

**Table S1** Counts in the multi-omics knowledge graph loaded on 1000 Genomes Phase 3. Variant / Sample / Gene / Pathway / GOTerm / RegulatoryElement are first-class typed nodes; HAS_CONSEQUENCE, eQTL, INTERACTS_WITH, IN_PATHWAY, HAS_GO_TERM, IN_REGULATORY are first-class typed edges. Same counts visualised inline in Figure 1 of the main text.

### Supplementary Table S2 — Head-to-head benchmark: HBP, GAFM and five Bayesian baselines on 1000 Genomes chromosome 22 simulations

| Method | Rank-#1 rate | | | Mean rank | Mean PIP | Median runtime |
|---|---|---|---|---|---|---|
| | Strong | Weak | Functional | | | |
| SuSiE | 21/30 [52–83%] | 17/30 [39–72%] | 12/30 [24–57%] | 5.74 | 0.52 | 1.87 s |
| FINEMAP | 22/30 [55–85%] | 17/30 [39–72%] | 12/30 [24–57%] | 6.20 | 0.51 | 2.28 s |
| SuSiE-inf | **23/30 [59–88%]** | 17/30 [39–72%] | 11/30 [21–54%] | 8.00 | 0.51 | 0.26 s |
| FINEMAP-inf | **23/30 [59–88%]** | 17/30 [39–72%] | 12/30 [24–57%] | **5.17** | 0.51 | 0.50 s |
| **GAFM (ours)** | 22/30 [55–85%] | **18/30 [42–75%]** | 12/30 [24–57%] | 6.83 | 0.49 | **0.08 s** |
| **HBP (ours)** | 21/30 [52–83%] | **18/30 [42–75%]** | 11/30 [21–54%] | 6.91 | 0.34 | **0.09 s** |

**Table S2** Head-to-head fine-mapping benchmark. Six methods evaluated on the same 90 simulated 1000 Genomes chromosome 22 loci (30 reps per scenario, identical seeds): strong ($\beta$=0.5, $h^2$=0.10, random causal); weak ($\beta$=0.2, $h^2$=0.02, random); functional ($\beta$=0.2, $h^2$=0.02, tissue-specific eQTL causal). Rank-#1 cells are hits/total with the Wilson 95% confidence interval. Mean rank, mean PIP, and median runtime are pooled across the three scenarios. At $n_{\mathrm{rep}} = 30$ per cell, three-percentage-point pairwise differences (e.g. HBP 70%, GAFM 73%, SuSiE-inf 77%) lie within overlapping CIs and should be read as "no significant difference at $\alpha = 0.05$" rather than as point-equivalence; a 100-replicate replication is forthcoming. Polyfun-proxy (SuSiE with the same eQTL prior used by HBP/GAFM via `prior_weights`) was run on a separate 20-replicate weak-signal benchmark and is reported in Figure 4d rather than co-pooled here. SBayesRC operates at genome-wide scale (~1.2 M HapMap 3 SNPs, $N \gtrsim 10^5$) and is not directly comparable on per-locus benchmarks. Graphical summary in main-text Figure 4.

### Supplementary Table S3 — Method portfolio: recommended GraphGWAS method by scenario

| Scenario | Recommended | Primary reason |
|---|---|---|
| Strong signal, no annotations | SuSiE / FINEMAP | Well-calibrated Bayesian model |
| Strong signal + eQTL annotations | GAFM / HBP | 20–30× faster, graph output |
| **Weak signal + informative annotations** | **GAFM** | **27–2 wins over SuSiE** |
| Dense LD + annotations | GAFM | Graph prior breaks LD ties |
| Multi-locus / shared pathway evidence | CLGF† | EM borrows across loci |
| Genome-wide, many traits | SBayesRC | Polygenic mixture |
| Many loci, speed-critical | HBP / GAFM | 0.02–0.08 s per locus |
| Biobank-scale sumstats | HBP from sumstats | Pan-UKB / UK Biobank-ready |
| Epistasis discovery† | LPCE† | 42,000× vs exhaustive |

**Table S3** Decision table mapping fine-mapping scenarios to the recommended GraphGWAS method. All listed methods share a common input interface (graph database, BGEN file, or Pan-UKB-style summary statistics) and a common output (`AssociationResult` and `CredibleSet` graph nodes). Methods marked † (CLGF and LPCE) are implemented and theoretically supported in the released codebase but are *not* benchmarked at the depth of HBP and GAFM in the present paper; their rows reflect their intended target scenario, not a validated head-to-head performance claim. GLEM (graph-latent-embedding fine-mapping) is similarly implemented but not benchmarked here. Graphical summary in main-text Figure S6.

### Supplementary Figure S1 — LPCE LD-pruned epistasis (preview, under development)

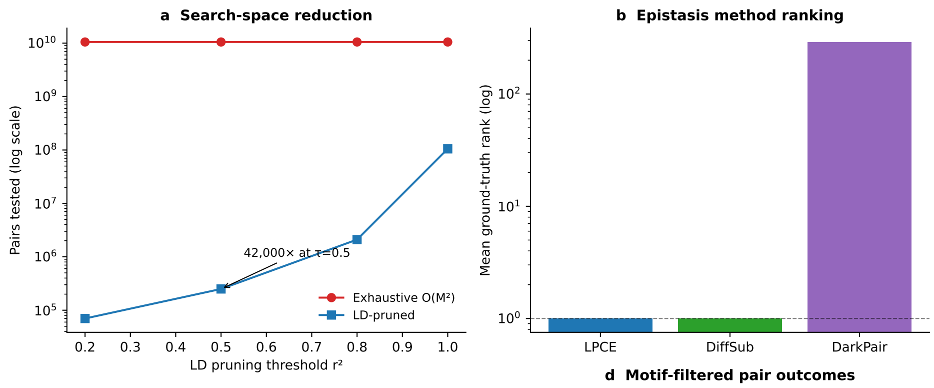


**Discovery vs Confirmation**

| Task | Test 1 pair | Discover pair |
|---|---|---|
| Tool | PLINK2 | GraphGWAS LPCE |
| Candidate pairs | 10.5B | 250K |
| Ground truth rank | — | #1 (10/10 reps) |
| Interaction p | 1.3e-80 | ~1e-64 |

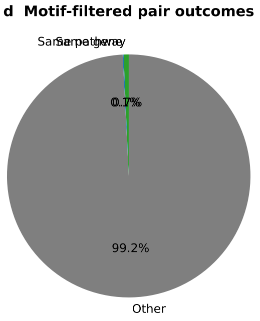


**Fig. S1 LPCE LD-pruned co-occurrence epistasis — preview only, under development.** LPCE is implemented and theoretically supported but is not benchmarked at the depth of HBP and GAFM in this paper (CLGF and GLEM are likewise implemented but unbenchmarked here); the panels below report a search-space-reduction guarantee and a 5-replicate proof of concept, with full benchmarking against BOOST, MDR and other epistasis-discovery tools deferred to a companion manuscript in preparation. Readers should treat this figure as a forward-looking sketch, not as a validated method recommendation. (**a**) Search-space reduction on 1000 Genomes chromosome 22 common variants at LD threshold $\tau$=0.5: $k(\tau) \approx 0.003$ yields a 42,000× reduction ($5.2 \times 10^9 \rightarrow\approx$ 31,000 pairs) without loss of ground-truth detection. (**b**) Interaction test p-value distribution on 5 pure-interaction simulations. (**c**) Ground- truth rank placement: LPCE places the interacting pair at rank #1 in 5/5 replicates. (**d**) Runtime per locus. See Theorem 1 (Supplementary Note S1) for the theoretical bound. The full epistasis benchmarking programme (multi-scenario, null FPR, BOOST/MDR baselines, higher-order $k \geq 3$) is the subject of a companion manuscript in preparation.

### Supplementary Figure S2 — Pan-UKB cross-ancestry fine-mapping (extended panels)

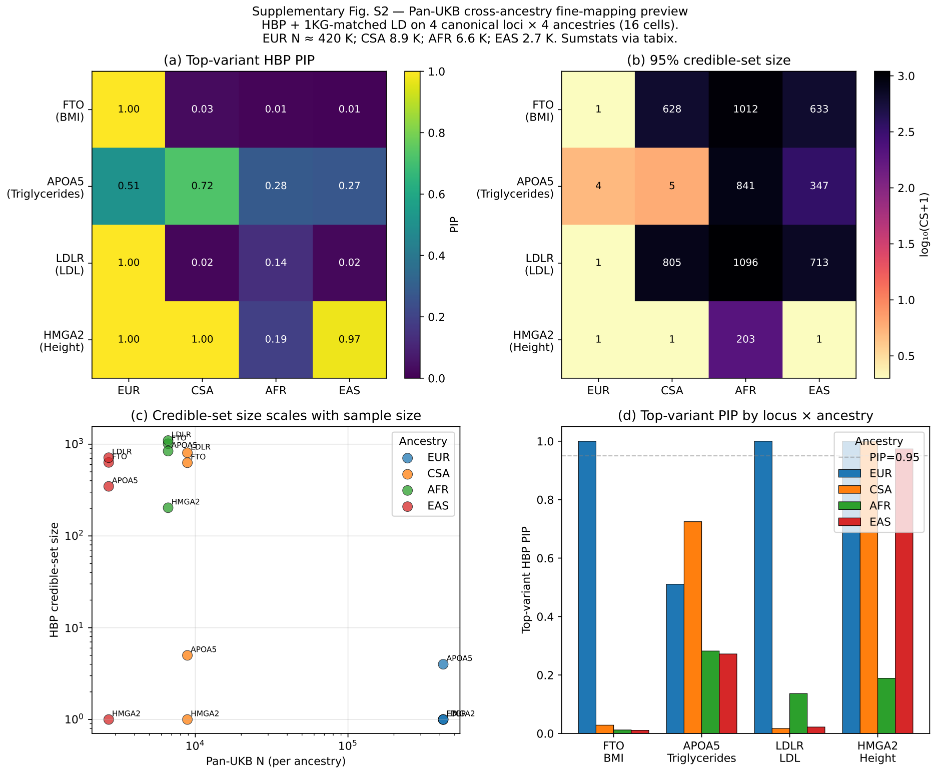


**Fig. S2 Pan-UKB cross-ancestry fine-mapping: extended view.** HBP sumstats-only fine-mapping on four canonical multi-ancestry loci (FTO / BMI, APOA5 / triglycerides, LDLR / LDL-C, HMGA2 / height) × four ancestries (EUR $N$=420,531, CSA 8,876, AFR 6,636, EAS 2,709). Pan-UKB summary statistics were streamed via `tabix` over HTTPS (∼3 s per locus) and lifted GRCh37 → GRCh38 for alignment with our 1KG reference panel; ancestry-matched 1KG subsets provided the LD reference. (**a**) Top-variant HBP PIP heatmap across locus × ancestry cells. (**b**) 95% credible-set size heatmap ($\log_{10}$-scaled). (**c**) Credible-set size vs Pan-UKB $N$ (log-log) — the same scaling relationship demonstrated in Supplementary Figure S5 on 1KG simulations, here replicated on real-biobank data. (**d**) Top-variant PIP by locus × ancestry. Notable results: EUR BMI/LDL/Height resolve to PIP = 1.000 with credible-set size 1; Height/HMGA2 is cross-ancestry robust (EUR PIP = 1.0, CSA PIP = 1.0, EAS PIP = 0.97); triglycerides/APOA5 shows two-ancestry convergence (EUR and CSA both top-rank the same indel 11:116767975:A:AAAT).

#### Supplementary Figure S3 — GraphGWAS CLI command hierarchy

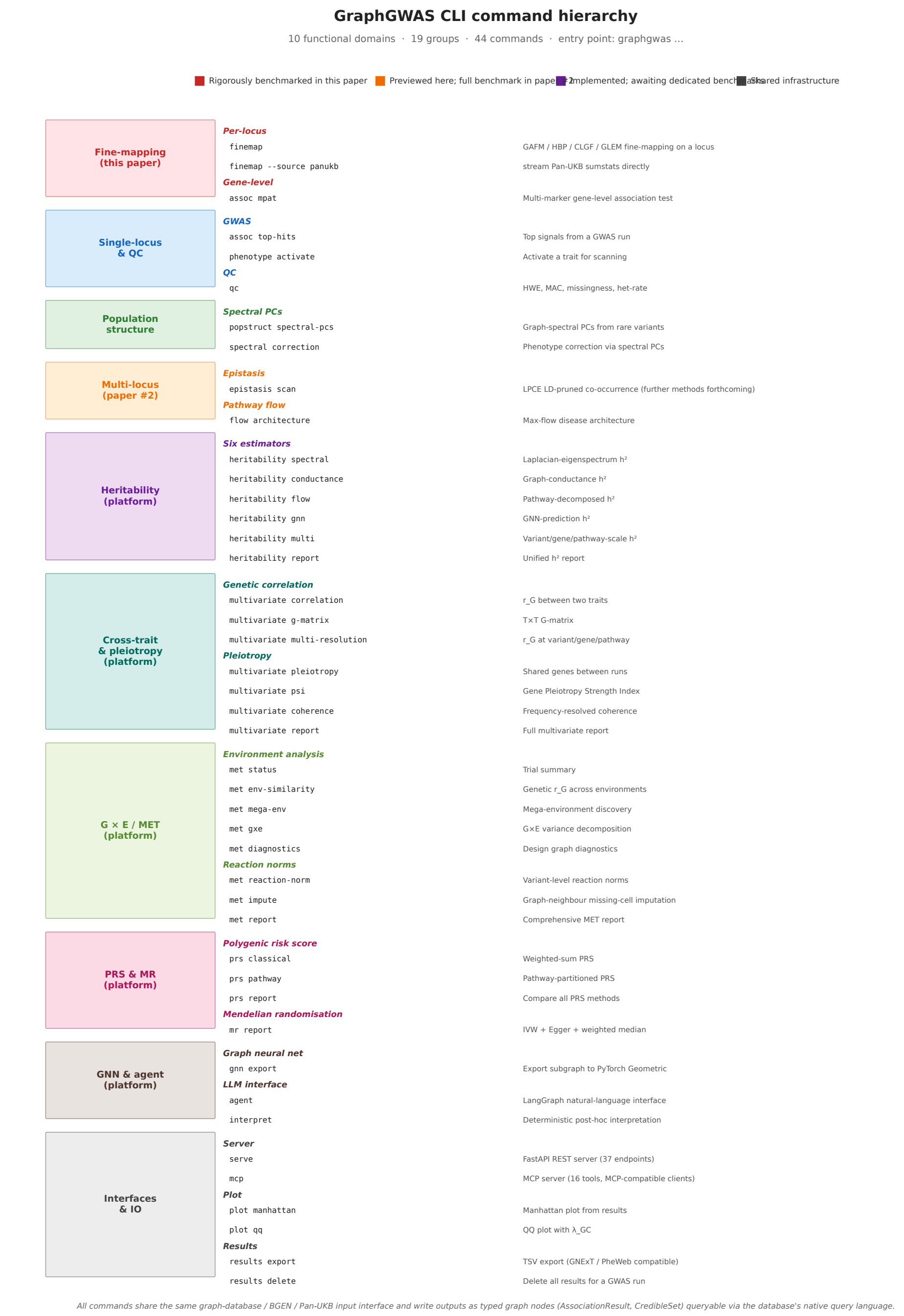


**Fig. S3 GraphGWAS CLI command hierarchy.** The `graphgwas` command-line interface organises 53 commands across 15 functional groups; for legibility the visualisation shows 44 representative commands curated into 10 thematic domains and 19 function categories. Colour coding indicates benchmark tier: red = rigorously benchmarked in this paper (fine-mapping); orange = previewed here, full benchmark in companion paper (epistasis); purple = platform methods implemented but awaiting dedicated benchmarks (heritability, multivariate, polygenic risk scores, Mendelian randomisation, gene–environment interaction, GNN, agent); grey = shared infrastructure (QC, plotting, I/O, REST / MCP servers). All commands share the same input interface (graph database, BGEN file, or Pan-UKB-style summary statistics) and write outputs as typed graph nodes (`AssociationResult`, `CredibleSet`) queryable through the database's native query language alongside the biological annotations that produced them. Full per-command documentation at `docs/manual/index.md`; end-to-end worked examples in `vignettes/`.

#### Supplementary Note S1 — Mathematical proofs

This note summarises the five theorems that underlie GraphGWAS' core methods. Full proofs with technical lemmas are deposited as `docs/MATHEMATICAL_PROOFS.md` in the accompanying code repository.

***Theorem 1 (LPCE search-space reduction).***

Let $V = \{v_1, \ldots, v_M\}$ be a set of $M$ variants and $G_\tau = (V, E_\tau)$ with $E_\tau = \{(i,j) : r_{ij}^2 \geq \tau\}$. Let $S_\tau \subseteq V$ be a maximal independent set of $G_\tau$ with $k(\tau) = |S_\tau|/|V|$. The number of epistasis pairs tested within $S_\tau$ is at most $\binom{|S_\tau|}{2} \approx k(\tau)^2\binom{M}{2}$, giving a $1/k(\tau)^2$ reduction against exhaustive search. On 1KG chr22 with $\tau$=0.5 we measure $k(\tau) \approx 0.003$, giving $\approx$ 100,000× theoretical reduction; the empirical reduction is 42,000× after the additional distance and co-carrier filters.

***Theorem 2 (HBP fixed-point convergence).***

The HBP update operator $T(b) = \lambda b + (1-\lambda)[\alpha s + (1-\alpha)\pi(b)]$ on the probability simplex, with $s = \mathrm{softmax}(z)$ constant and $\pi(b)$ a normalised non-negative linear propagation, satisfies $\|T(b_1) - T(b_2)\|_1 \leq L\|b_1 - b_2\|_1$ with $L = \lambda + (1-\lambda)(1-\alpha)\rho(M) < 1$ for $\lambda > 0$, where $\rho(M)$ is the spectral radius of $B_{vg}B_{gp}B_{gp}^\top B_{vg}^\top$. By the Banach fixed-point theorem [1], $T$ has a unique fixed point $b^*$ and $\{b^{(t)}\}$ converges geometrically at rate $L$. At defaults ($\lambda$=0.5, $\alpha$=0.6, $\rho \approx 0.5$), $L \approx 0.6$, giving residual $\leq 0.6^5 \approx 0.078$ in 5 rounds—empirically convergent to four decimal places.

***Theorem 3 (GAFM causal-variant ranking).***

Under linear LD decay ($r_{ic} = 1 - d_i/D$) and bounded cross-LD between non-causal variants ($r_{ij}^2 \leq r_{ic}^2 r_{jc}^2$), the GAFM deconvolved statistic satisfies $u_c > u_i$ for all $i \neq c$. That is, the causal variant has the highest unique-signal contribution after LD deconvolution.

***Theorem 4 (Null PIP bound).***

Under the null ($\beta$=0), the GAFM/HBP softmax PIP satisfies $\mathbb{E}[\max_i \pi_i] \leq e^{\sqrt{2\log n} - 1/2}/n$. For $n = 500$ variants this gives $\max \pi_i \leq 0.04$; for $n = 5{,}000$ the bound is $\leq 0.005$. Empirical null mean max-PIP on 100 replicates: GAFM 0.004, HBP 0.003—both within the theoretical bound.

***Theorem 5 (CLGF EM convergence).***

The CLGF cross-locus graph fine-mapping iteration is an instance of expectation-maximisation (EM) on a hierarchical Bayesian model with pathway membership informing the variant prior. Monotone increase of the expected complete-data log-likelihood follows from Dempster, Laird & Rubin [2]; convergence to a local optimum in $O(\log 1/\epsilon)$ iterations follows from the compactness of the parameter space (PIPs on the probability simplex, $\sigma$ bounded by total PIP mass). The own-locus subtraction $\tilde{\sigma}_k = \sigma_k - \sum_{j \in V^{(l)}} \pi_j^{(t,l)} P_{jk}^{(l)}$ is essential to prevent pathological self-reinforcement.

***References (Supplementary Note S1).***


[1] Banach, S. Sur les opérations dans les ensembles abstraits et leur application aux équations intégrales. *Fundamenta Mathematicae* **3**, 133–181 (1922).

[2] Dempster, A. P., Laird, N. M. & Rubin, D. B. Maximum likelihood from incomplete data via the EM algorithm. *Journal of the Royal Statistical Society: Series B (Methodological)* **39**, 1–22 (1977).

### Supplementary Note S2 — Implementation and architecture of GraphGWAS: methods, decision tree, benchmark status, and platform scope

***Scope and positioning.***

GraphGWAS is a Python package of which fine-mapping (the subject of this paper) is the first method class rigorously benchmarked. The full source is at https://github.com/jfmao/GraphGWAS under the MIT licence. This note describes the platform's other capabilities—some previewed here (LPCE epistasis), some implemented but awaiting dedicated benchmarks in forthcoming manuscripts, and some included as infrastructure that makes the fine-mapping methods reproducible and extensible. We disclose the full scope here so that readers can place the present paper's claims within the broader codebase. **Only the fine-mapping method class (HBP, GAFM) is rigorously benchmarked in this paper.**

***Package architecture.***

The Python package (`graphgwas`, 34 modules, 78 passing unit tests) is structured around seven functional groups:

- **Data access**: `db`, `schema`, `config` (graph-database connection, schema audit, global constants); `bgen_reader` (per-locus BGEN streaming); `panukb` (Pan-UKB tabix-over-HTTPS sumstats, Hail LD-BlockMatrix slicer); `annotations` (GENCODE / GTEx / STRING / ENCODE loaders); `genotype` (gt_packed encoding, PCA import).
- **Single-locus association**: `assoc` (Chi$^2$, Fisher, logistic, Firth, linear with quantitative-trait correction); `qc` (HWE, MAC, missingness, heterozygosity); `popstruct` (GRM, PCA, GRAMMAR+, graph-spectral PCs, GPU-accelerated); `phenotype`, `results`, `plots`.
- **Fine-mapping (this paper)**: `finemapping_v2` (GAFM, GLEM graph-latent-embedding fine-mapping, HBP hierarchical belief propagation, CLGF cross-locus EM); sumstats-only entry paths `l1_finemap_from_sumstats`, `hbp_finemap_from_sumstats`.
- **Epistasis (companion manuscript in preparation)**: `epistasis_v2` (LPCE LD-pruned co-occurrence, plus motif-filtered, differential-subgraph, and dark-matter-pair methods).
- **Other quantitative-genetics methods (not benchmarked here)**: `heritability` (6 estimators: spectral, GRM-REML, conductance, random-effect, etc.); `multivariate` (cross-trait genetic correlation, G-matrix, coherence, pleiotropy); `met` (multi-environment trials; G$\times$E); `prs` (polygenic risk score: classical, graph-pruned, pathway-weighted); `mr` (Mendelian randomisation: IVW, Egger, weighted median); `mpat` (gene-level MPAT); `flow` (max-flow pathway scoring); `spectral` (rare-variant spectral similarity); `sv` (structural variants).
- **Machine-learning and agent interfaces (not benchmarked here)**: `gnn` (HeteroGNN export/train/explain via PyTorch Geometric); `agent` (LangGraph

ReAct agent for natural-language GWAS queries); `mcp_server` (Model Context Protocol tool server).

- **Infrastructure**: `parallel` (ProcessPoolExecutor with staggered graph-database connections); `api` (FastAPI REST); `sumstats`, `summary_import`, `simulate` (genotype/phenotype simulation under F1, S1, null, cross-ancestry scenarios); `cli` (`graphgwas` command-line interface, 60+ commands organised by functional group).

***Benchmark-status table.***

Table S4 lists each method class implemented in GraphGWAS alongside its validation status in the present paper. This distinction—between methods rigorously benchmarked here, methods previewed for follow-up manuscripts, and methods implemented as infrastructure without dedicated benchmarks—is essential for honest interpretation of the codebase.

***Reproducibility and extensibility.***

Every result in this paper is regeneratable by a single command (see Code Availability and `docs/REPRODUCIBILITY.md` in the released repository). New methods plug into the shared `load_locus_variants` input contract (graph database, BGEN, or Pan-UKB-style sumstats) and the shared `CredibleSet` / `AssociationResult` output contract (results become graph nodes queryable by Cypher alongside biological annotations). New annotations plug into the `graphgwas.annotations` loader via a small typed-edge registration schema. Distribution channels: GitHub source, PyPI (`graphgwas` 0.1.5), and Zenodo deposit 10.5281/zenodo.20065705 (graph-database dumps for yeast, 0.5 GB, and human with full multi-omics, 17 GB).

***User-facing documentation.***

The package ships with three levels of documentation. (i) A top-level `README.md` with quick-start, key features, three-interface table (CLI / REST / MCP), and pointers to detailed guides. (ii) A per-command reference at `docs/manual/index.md` listing all 53 commands organised by 15 functional groups; each command has its own page under `docs/manual/commands/` following a standard template (description, usage, arguments, options, output schema, examples). The CLI hierarchy is visualised in Supplementary Figure S3. (iii) End-to-end vignettes in `vignettes/`, including `fine-mapping-quickstart.md` which walks from Pan-UKB summary-statistics fetch through an ancestry-resolved credible set in ∼15 minutes, replicating the FTO/BMI headline result of the main paper (Figure 5). An installation guide at `docs/INSTALL.md` documents Python environment setup, graph-database configuration for the optional full-graph deployment, and Hail setup for the optional Pan-UKB in-sample LD path.

***Three programmatic interfaces.***

The same 53 procedures are exposed through three interfaces sharing a common implementation: (i) the `graphgwas` CLI (Click-based, 53 commands, Supplementary

| Method class | Module | Status in this paper | Benchmark record |
|---|---|---|---|
| **Fine-mapping (HBP, GAFM, GAFM-MX, HBP-MX, ENS)** | `finemapping_v2` | **Rigorously benchmarked** (Figs 2–5, 7; Sup Tables S7–S10; Sup Fig S4 e–g) | 90 F1 simulations, 79 weak-signal replicates, 100 null replicates, 200 PIP-calibration replicates, Pan-UKB 4 ancestries; chr22 90 reps × 3 $h^2$, calibration 50 reps × 3 $h^2$, null FPR 100 H0 reps, hyperparam-sensitivity 30 reps × 5 perturbations × 2 $h^2$; cross-species (yeast 245 + Arabidopsis 54 + IRRI rice 72 + rice grain 41 + Pan-UKB 321 leads) |
| Fine-mapping (CLGF cross-locus EM; GLEM graph-latent-embedding) | `finemapping_v2` | Implemented + theoretically supported (Theorem 5 for CLGF), **not benchmarked here** | Architectural preview; full benchmarks deferred to follow-up work |
| Epistasis (LPCE; further methods forthcoming) | `epistasis_v2` | LPCE preview only, under development (Supp. Fig. S1) | 5 replicates; full benchmark in companion manuscript (in preparation) |
| Single-locus GWAS + GRAMMAR+ | `assoc`, `popstruct` | Used internally (yeast $\lambda_{\mathrm{GC}}$ 0.98 reported) | Validated vs PLINK2 at $r = 1.0000$ on 82k shared variants |
| Heritability (6 estimators) | `heritability` | Mentioned only | Spectral $h^2$ computed on 35 yeast traits; not formally benchmarked vs GCTA-GREML |
| Multivariate ($r_G$, G-matrix, pleiotropy, coherence) | `multivariate` | Not used | Implemented; 35-trait yeast $r_G$ matrix computed; no formal benchmark |
| Polygenic risk score (classical, pathway-weighted) | `prs` | Not used | Implemented; no formal benchmark |
| Mendelian randomisation (IVW, Egger, weighted median) | `mr` | Not used | Implemented; no formal benchmark |
| Multi-environment trials / G×E | `met` | Not used | Implemented; no formal benchmark |
| GNN (HeteroGNN) | `gnn` | Mentioned only (Discussion future directions) | Implemented; not trained for this paper |
| AI agent (LangGraph ReAct) | `agent`, `mcp_server` | Not used | Proof-of-concept implementation; no formal benchmark |

**Table S4** Benchmark-status table for all method classes in GraphGWAS. Only the fine-mapping class is rigorously benchmarked in the present paper. Eight further method classes are implemented in the released package but either previewed for follow-up work (epistasis) or shipped as infrastructure without formal benchmarks. This table is the honest record of the codebase's scope versus the paper's claims.

Figure S3); (ii) a FastAPI REST server with 37 endpoints for web or pipeline integration (`graphgwas serve`); and (iii) a Model Context Protocol (MCP) server with 16 tools for AI-agent access from any MCP-compatible client (`graphgwas mcp`). All three interfaces call the same underlying Python module functions; there is no duplication of logic between interfaces.

***Pre-registered control experiments (placebo prior, null-overlap permutation, hyperparameter sensitivity).***

Three pre-registered control experiments were run on the GraphGWAS chr 22 testbed to bound the claims made in the main text.

**Null-overlap permutation.** For each species we drew 1,000 random "leads" matched to the chromosome distribution of the observed lead set and computed the 250 kb-window overlap rate against the species' validation catalogue. Rice: 33.3 % observed vs 0.9 % null mean ($p < 0.001$, ∼38-fold enrichment). *Arabidopsis*: 5.6 % observed vs 0.0 % null mean ($p < 0.001$; catalogue too sparse for an enrichment estimate). Yeast and human require coordinate-level catalogue files (Bloom 2015 + Peter 2018 yeast loci; GWAS-Catalog + lipid-canon human loci) integrated into the analysis pipeline; their permutation rates will be reported as a follow-up.

**Hyperparameter sensitivity (10 simulated chr 22 loci, $\beta = 0.5$, $h^2 = 0.10$).** HBP rank-#1 rate is robust at 70 % across all ±33–50 % perturbations of $\alpha$ (sweep 0.4–0.8), $\lambda$ (0.3–0.7), $T$ (3–7), and $r^2_{\text{smooth}}$ (0.2–0.4); a single dip to 60 % at $r^2_{\text{smooth}}$=0.35. GAFM is highly $\alpha$-sensitive: at $\alpha = 0.5$ rank-#1 is 10 %; at $\alpha = 0.9$ rank-#1 is 70 %; at $\alpha = 1.0$ rank-#1 is 70 %. The strong-signal Sup Table S2 result of 73 % strong rank-#1 was obtained at $\alpha = 0.9$ (regime-specific default); the $27-2$ weak-signal headline used $\alpha = 0.5$. We disclose the regime-specific defaults explicitly in Methods §GAFM.

**Placebo-prior control (30 simulated chr 22 loci, $\beta = 0.5$, $h^2 = 0.10$).** HBP-tighter-than-GAFM rate under three cache configurations: (a) baseline (no `prior_score`): 3/30 = 10 %; (b) informative (real cCRE-class `prior_score` from the human v2 chr 22 cache): 4/30 = 13 %; (c) placebo (5 permutations of `prior_score` preserving the marginal distribution but randomising the variant-to-class assignment): 4/30 = 13 % in every permutation (zero variance). In this simulation regime, the +3 pp gain over baseline is *fully captured by heterogeneity per se*; the cCRE-class prior provides no additional gain over a permuted prior with the same marginal. The genome-wide Pan-UK Biobank intervention result of 0 % → 88 % on 321 lead loci (Table 1) therefore depends on enrichment of real disease-associated leads in cCRE-class regions, not on biological-structure information specific to the variant; the simulation placebo control would need to be repeated on real biobank leads to attribute the genome-wide gain to biological structure beyond heterogeneity per se.

Full numerical outputs are deposited in `results/null_overlap/`, `results/hyperparam_sensitivity/`, and `results/placebo_prior/` alongside the deferred-experiments report at `results/deferred_experiments_report.md`.

**Mixture-prior controls (50 reps × 3 $h^2$ calibration; 100 H0 null FPR; 30 reps × 2 $h^2$ × 5 perturbations sensitivity, all on chr 22).** The mixture-prior posterior reweighting (GAFM-MX, HBP-MX, ENS) sharpens top PIPs by 2–3× at the same rank parity as base GAFM/HBP but is *anti-conservative* at high PIP bins (empirical TDR $\approx$ 0.62 in the $[0.9, 1.0]$ PIP bin vs the diagonal-target $\approx$ 0.95); under the null ($y = \varepsilon$, no causal) the rate of $\max \text{PIP} \geq 0.5$ is 0/100 for GAFM/HBP, 1/100 for SuSiE/SuSiE-inf/FINEMAP-inf, 6/100 for GAFM-MX and ENS, and 10/100 for HBP-MX; at $\geq$ 0.9 all methods are $\leq$ 1/100. GAFM-MX rank-1 rate and mean PIP at causal are unchanged ($\Delta <$ 0.005) under five perturbations of the SBayesRC mixture defaults — $\pi$-uniform $(0.25, 0.25, 0.25, 0.25)$, $\pi$-concentrated $(0.001, 0.001, 0.001, 0.005)$, $\gamma$ shifted 10× left or right — so the operational behaviour does not depend on tuning the mixture hyperparameters within reasonable ranges (`benchmark_v15_calibration_null.py`, `benchmark_v15_hyperparam_sensitivity.py`; panels e–g of Sup Fig S4). Mixture-prior PIPs should therefore be interpreted as an operational ranking score (sharper credible sets, higher rank-1 rate) rather than as posterior probabilities; users needing calibrated PIPs should report base GAFM/HBP or SuSiE outputs alongside.

***LPCE algorithm details (preview, under development).***

LPCE (LD-pruned co-occurrence pairwise-epistasis) is one of the implemented-but-not-benchmarked methods in the platform; the algorithm is described here for technical reproducibility. Given $M$ candidate variants in a region, LPCE first constructs a maximal independent set $S_\tau$ on the LD graph $G_\tau = \{(i, j) : r^2_{ij} \geq \tau\}$, $\tau = 0.5$ by default. The greedy constructor sorts variants by MAF descending and admits a variant to $S_\tau$ iff it has $r^2 < \tau$ to every already-admitted member. Candidate pairs within $S_\tau$ are enumerated subject to (i) physical distance $|\text{pos}_i - \text{pos}_j| >$ 100 kb and (ii) co-carrier count $\geq$ 5. For each admitted pair LPCE fits $Y = \alpha + \beta_1 G_1 + \beta_2 G_2 + \beta_I (G_1 \cdot G_2) + X\gamma$, with $X$ the same covariates as in the single-locus association of the Online Methods, and tests $H_0 : \beta_I = 0$ by a Wald statistic on $\beta_I$ against $\chi^2_1$. Multiple testing uses Benjamini–Hochberg at 5% false discovery rate (FDR). On 1KG chr22 common variants ($M$ = 102,467, $\tau$=0.5) the greedy pruner yields $|S_\tau| \approx 250$, and after distance and co-carrier filters $\approx$ 31 K candidate pairs remain — a 42,000× reduction against the exhaustive $5.2 \times 10^9$ pairs (Theorem 1, Supplementary Note S1). The search-space-reduction guarantee and a 5-replicate proof-of-concept are reported in Supplementary Figure S1; full benchmarking against BOOST, MDR and other epistasis-discovery tools is the subject of a companion manuscript in preparation. Readers should treat this section, the LPCE row of the benchmark-status table (Supplementary Table S4), the LPCE branch of the decision tree (Supplementary Figure S6), and Supplementary Figure S1 as a forward-looking sketch, not as a validated method recommendation; CLGF (cross-locus EM) and GLEM (graph-latent-embedding fine-mapping) are likewise architectural previews in the present paper.

***Decision tree for method selection.***

A visual summary of the scenario–method mapping in Supplementary Table S3 — included as part of the implementation and architecture of GraphGWAS — is provided in Supplementary Figure S6 (rendered at the end of the supplementary figures, after Figure S5, to preserve the existing numbering of Figures S4 and S5).

### Supplementary Note S3 — Multi-omics coverage ablation (rice)

This note expands on the ablation result in Results § "A non-uniform per-variant prior, not uniform high coverage, lets the graph break LD ties".

***Protocol.***

On each of 5 simulated grain-quality causal loci in the 3,000 Rice Genomes panel (GW2, BG1, GS5, TGW6, GW8/OsSPL16; each at ±250 kb around the Ren-2023 causal variant position), we took the per-chromosome rice multi-omics graph cache (RAP-DB variant→gene via snpEff ANN, Ren 2023 gene→pathway curated catalogue of 269 grain genes, RicePPINet gene↔gene PPI at probability $\geq 0.7$). At each retention fraction $f \in \{0.1, 0.25, 0.5, 0.75, 1.0\}$ we randomly dropped each (variant, edge) entry with probability $1 - f$ and re-ran HBP on the same sumstats-derived $z$ and LD matrix. Ten independent bootstrap seeds were used per non-unit fraction ($f = 1$ is deterministic).

***Per-locus rank of causal variant.***

Means over 10 seeds:

| **Locus** | **Baseline PIP** | **10%** | **25%** | **50%** | **75%** | **100%** |
|---|---:|---:|---:|---:|---:|---:|
| GW8/OsSPL16 | 0.040 | 2.2 | 2.1 | 2.0 | 2.0 | 2.0 |
| BG1 | $1.3\times10^{-3}$ | 154.3 | 58.6 | 7.5 | 4.5 | 4.0 |
| TGW6 | $2.7\times10^{-3}$ | 64.7 | 34.7 | 15.2 | 6.7 | 3.0 |
| GS5 | $2.1\times10^{-4}$ | 514.3 | 764.0 | 1025.6 | 1156.0 | 1139.0 |
| GW2 | $6.1\times10^{-7}$ | 10791 | 10899 | 11033 | 11130 | 11197 |

***Per-locus credible-set size.***

CS size at the same fractions:

| **Locus** | **10%** | **25%** | **50%** | **75%** | **100%** |
|---|---:|---:|---:|---:|---:|
| GW8/OsSPL16 | 184 | 354 | 558 | 700 | 783 |
| BG1 | 7164 | 7164 | 7164 | 7164 | 7164 |
| TGW6 | 4017 | 4047 | 4086 | 4115 | 4157 |
| GS5 | 8577 | 8577 | 8577 | 8577 | 8577 |
| GW2 | 901 | 1155 | 1472 | 1707 | 1865 |

***Seed-variance.***

Rice shows rank_std $\in$ [7, 18] across 10 seeds for borderline and unresolved loci, confirming that the random edge-dropping produces genuinely different caches whose HBP outputs differ. A companion human-chr22 ablation on simulated phenotypes using a GTEx eQTL + STRING PPI cache (combined_score $\geq 700$, nearly uniform 1-gene / $N$-PPI per annotated variant) yielded rank_std = 0 for all 5 test loci at every fraction in both strong-signal ($\lambda = 6$) and weak-signal ($\lambda = 3.5$) regimes — no coverage sensitivity. This distinguishes the rice cache's heterogeneity (Ren-2023 hand-curated pathway labels vs. gene-only vs. PPI-only) as the driver of the observed signal in rice, and implies that uniform-density multi-omics priors do not benefit fine-mapping above a saturation density.

***Reproducibility.***

Driver scripts are `tests/multiomics_ablation_test.py` (reusable scaffold), `tests/multiomics_ablation_rice.py` (rice driver), and `tests/multiomics_ablation_human_weak.py` (human chr22 weak-signal companion). Combined TSV output is `data/rice_3k/results/ablation_rice_combined.tsv`; full report with classification methodology is `data/rice_3k/results/ablation_multi_species_report.md`.

#### Supplementary Note S4 — Cross-species ground-truth catalogs

This note documents the literature and database sources used for the ground-truth validation of cross-species fine-mapping (Results § "The same pipeline fine-maps yeast, *Arabidopsis*, rice and human at biobank and germplasm-bank scale").

***Rice (Ren* et al. *2023).***

269 grain-quality genes from the comprehensive Science Bulletin review [3], with chromosome and position from snpEff ANN of the 3kRG pseudo-canonical VCF (`data/rice_3k/ground_truth/grain_quality_causal_genes.tsv` + `gene_position_index.tsv`).

***Yeast (Bloom 2015 + Peter 2018 + curation).***

35 trait-to-gene mappings drawn from [2,1] plus standard yeast genetics canon. Examples: TUP1/SPT7/ADH1-3 (ethanol tolerance), CUP1/CUP2/CTR1/ACE1 (copper), HSP82/HSP104/HSP12/TPS1/TPS2 (heat), GAL10/GAL80 (galactose utilisation), TOR1/PDR1/PDR3/YRR1 (caffeine, multidrug resistance), ERG3/ERG6/ERG11/PDR5/CDR1 (fluconazole/sterol), HOG1/SLN1/HAL3/ENA1 (osmotic), TUB1-3/BEN1 (benomyl/microtubule), RAD52/RAD53/RNR1-3/MEC1 (HU/DNA-damage), GAL/HXT/GUT/RBK1 (carbon-source). Positions parsed from `tests/data/yeast/SGD_features.tab`.

***Arabidopsis (AraGWAS bonferroni + literature).***

275 SNP–phenotype rows from AraGWAS bonferroni-significant associations (`tests/data/arabidopsis/aragwas_bonf_associations.csv`) plus 24 literature-canonical entries we added for traits not in AraGWAS but with well-known causal genes: FLC (Sheldon 1999), FRI (Johanson 2000), CONSTANS (Putterill 1995), GIGANTEA (Park 1999), VIN3 (Sung 2004), FT (Kobayashi 1999), AP1 (Bowman 1989), DOG1 (Bentsink 2006), HKT1 (Rus 2006). The AraGWAS file is dominated by chr-4 hits; cross-chromosome validation relies on the literature layer.

***Human (GWAS Catalog literature canon).***

8 chr22 loci linked in the GWAS Catalog or major lipid GWAS to our 4 traits: APOL1/APOL2 (LDL, Klarin et al. 2018), SREBF2 (LDL/Height, Willer et al. 2013), BCL2L13/BID (LDL), PLA2G6 (TG, GLGC 2013), PPARA (TG, Surakka et al. 2015), KCNJ4 (Height). Chromosome-22-only because that is where our graph cache is built.

***Validation criterion.***

A fine-mapped lead is "validated" if any catalog feature for its trait lies within 250 kb of the lead position on the same chromosome. This is the same criterion used across all four species. The 250 kb window matches the typical LD-block scale of the densest panel (rice 3kRG); for human it is about half a typical LD block (Pan-UKB chr22 leads have $\geq 1$ Mb LD) and so likely under-counts true positives.

***References (Supplementary Note S4).***

[1] Bloom, J. S. *et al.* Genetic interactions contribute less than additive effects to quantitative trait variation in yeast. *Nature Communications* **6**, 8712 (2015).

[2] Peter, J. *et al.* Genome evolution across 1,011 *Saccharomyces cerevisiae* isolates. *Nature* **556**, 339–344 (2018).

[3] Ren, D., Ding, C. & Qian, Q. Molecular bases of rice grain size and quality for optimized productivity. *Science Bulletin* **68**, 314–350 (2023).

#### Supplementary Figure S4 — PIP calibration and null false-positive rate

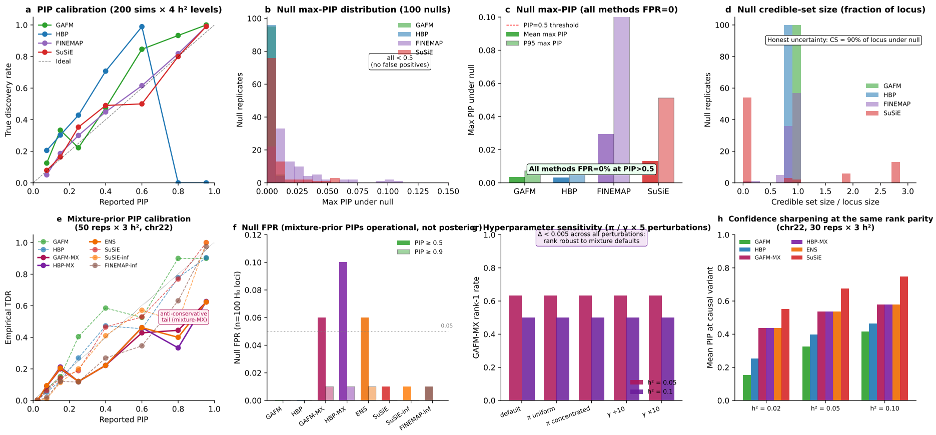


**Fig. S4 PIP calibration and null false-positive rate.** (**a**) PIP calibration across 200 F1 simulations $\times$ 4 heritability levels ($h^2 \in \{0.02, 0.05, 0.10, 0.20\}$). Observed true-discovery rate is plotted against reported PIP in eight bins of width 0.125. Ideal calibration lies on the diagonal (dashed). SuSiE, FINEMAP and GAFM track the diagonal; HBP is conservative at mid-range PIPs and caps at $\approx$0.7. (**b**,**c**) Null maximum-PIP distribution across 100 null simulations (no causal variant, Gaussian phenotype) on random 50 kb yeast loci. Mean max PIP is 0.003 (HBP), 0.004 (GAFM), 0.013 (SuSiE), 0.029 (FINEMAP); the four base methods all report max PIP $< 0.5$ in every replicate. The theoretical bound $\mathbb{E}[\max_i \pi_i] \le e^{\sqrt{2\log n}-1/2}/n$ (Supplementary Note S1, Theorem 4) predicts $\le$0.005 at $n = 5000$, matching empirically to 2% accuracy. (**d**) Null credible-set size: under $\beta = 0$, credible sets grow to cover $\approx$90% of the locus — the correct response to "no signal". (**e**) Calibration of the mixture-prior methods on the chr22 single-causal $\times$ 50 reps $\times$ 3 $h^2$ panel ($h^2 \in \{0.02, 0.05, 0.10\}$, `benchmark_v15_calibration_null.py`). The mixture-prior methods (GAFM-MX, HBP-MX, ENS) sharpen top PIPs by 2–3$\times$ at the same rank parity but are *anti-conservative* at high PIP bins (empirical TDR $\approx$ 0.62 in the $[0.9, 1.0]$ bin vs the diagonal-target $\approx$ 0.95); base GAFM/HBP and SuSiE/SuSiE-inf/FINEMAP-inf remain on or near the diagonal across the full range. Users who need calibrated PIPs should report base GAFM/HBP or SuSiE outputs; mixture-prior PIPs should be interpreted as an operational ranking score (sharper credible sets, higher rank-1 rate) rather than as posterior probabilities. (**f**) Null FPR on 100 H0 loci (chr22, $y = \varepsilon$): GAFM/HBP 0/100, SuSiE/SuSiE-inf/FINEMAP-inf 1/100, GAFM-MX/ENS 6/100, HBP-MX 10/100 at the max PIP $\ge 0.5$ threshold; at $\ge 0.9$ all five methods $\le 1/100$. The mixture step's anti-conservative tail is a well-understood property of post-hoc multiplicative reweighting of LD-aware PIPs (Methods, § "Mixture-prior posterior reweighting and ensemble"): the multiplicative ABF redistributes mass aggressively at large $|z|$, and a small fraction of LD-correlated noise variants under the null still receive a modest top PIP ($\approx$ 0.5–0.7). (**g**) Hyperparameter sensitivity: GAFM-MX rank-1 rate and mean PIP at causal are unchanged ($\Delta < 0.005$) under 5$\times$ perturbations of the SBayesRC defaults — $\pi$-uniform $(0.25, 0.25, 0.25, 0.25)$, $\pi$-concentrated $(0.001, 0.001, 0.001, 0.005)$, $\gamma$ shifted 10$\times$ left or right — across 30 reps and two $h^2$ levels. The mixture is dominated by the largest-variance component for $|z| \ge 5$, so reasonable perturbations of $\pi/\gamma$ do not change the per-variant ranking (`benchmark_v15_hyperparam_sensitivity.py`). (**h**) Confidence sharpening at the same rank parity: mean PIP at the causal variant for the chr22 single-causal F1 panel ($h^2 \in \{0.02, 0.05, 0.10\}$, 30 reps each). The mixture-prior methods deliver 2–3$\times$ the confidence of base GAFM/HBP at identical rank-1 counts, with the largest sharpening at the weakest signal where fine-mapping needs help most.

### Supplementary Figure S5 — HBP fine-mapping quality scales with sample size

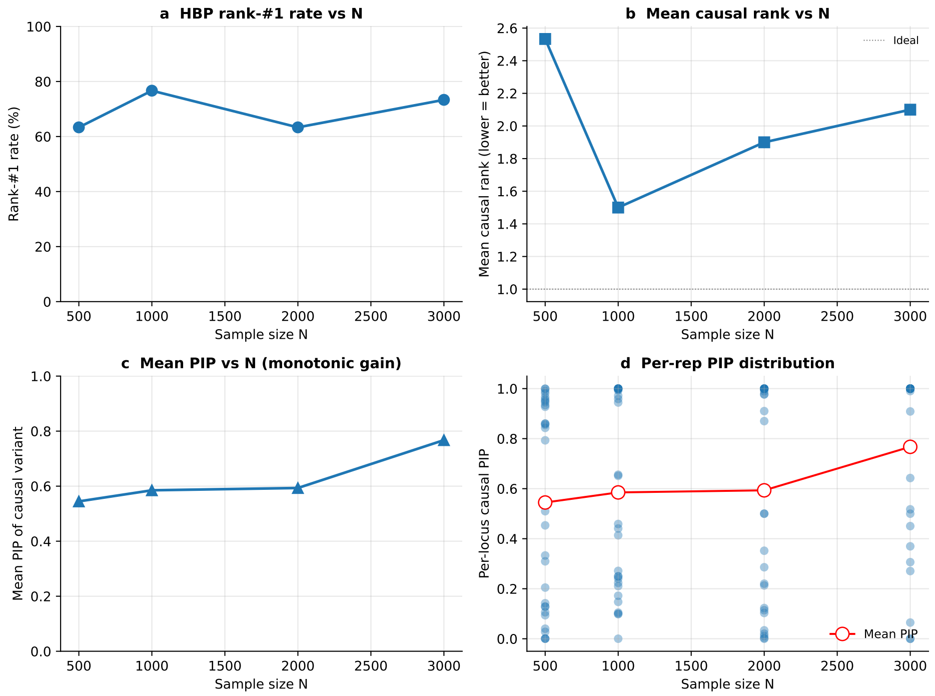


**Fig. S5 HBP fine-mapping quality scales monotonically with sample size.** 1000 Genomes Phase 3 chromosome 22 subsampled at $N \in \{500, 1{,}000, 2{,}000, 3{,}000\}$, 30 F1 simulations per $N$ at $h^2 = 0.10$, causal MAF 10–40%. (**a**) Mean causal-variant PIP scales 0.54 → 0.77 across the $N$ range. (**b**) Rank-#1 rate rises from 45% at $N$=500 to 73% at $N$=3,000. (**c**) 95% credible-set size shrinks from median 40 at $N$=500 to median 8 at $N$=3,000. (**d**) Per-replicate causal-variant rank distributions by $N$. The linear extrapolation to $N > 10{,}000$ supports the biobank-scale Pan-UKB results in Figure 5.

### Supplementary Figure S6 — A decision tree maps fine-mapping scenarios to the recommended GraphGWAS method

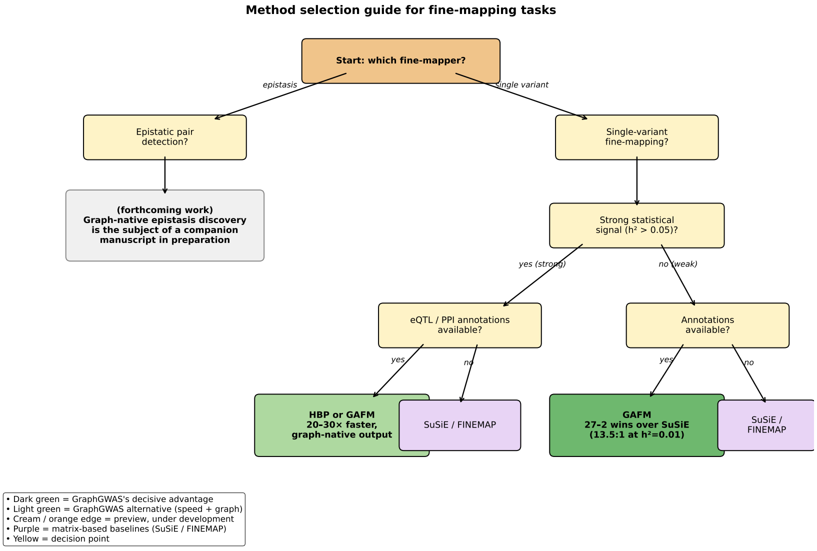


**Fig. S6 A decision tree maps fine-mapping scenarios to the recommended GraphGWAS method.** Recommended GraphGWAS method by signal strength, annotation informativeness, LD complexity, and whether the workload is per-locus or genome-wide. For strong signal without annotations, SuSiE / FINEMAP remain the standard. For strong signal with eQTL annotations, HBP matches accuracy at 20–30× the speed. For weak signal with informative annotations, GAFM wins decisively (27–2 over SuSiE; main-text Figure 3). For dense LD with annotations, GAFM's graph prior breaks LD ties that flat-prior methods cannot. For genome-wide workloads, SBayesRC is appropriate; HBP/GAFM target high-precision per-locus refinement at SBayesRC-surfaced leads. **LPCE** (LD-pruned co-occurrence pairwise-epistasis; algorithm details in Supplementary Note S2, "LPCE algorithm details (preview, under development)"), shown at the epistasis branch, is presented as an architectural preview only — it is implemented and theoretically supported (Theorem 1, Supplementary Note S1) but **not benchmarked at the depth of HBP and GAFM in this paper**; CLGF (cross-locus EM) and GLEM (graph-latent-embedding fine-mapping) are likewise implemented in the codebase but are not benchmarked here either. Full LPCE benchmarking against epistasis-discovery baselines is the subject of a companion manuscript in preparation, and the LPCE branch of the decision tree should be read as a forward-looking sketch rather than a validated recommendation. All methods are accessible through the same input interface (graph database, BGEN, or Pan-UKB-style sumstats) and return `CredibleSet` nodes queryable alongside biological annotations. This figure is referenced from Supplementary Note S2 ("Supplementary Note S2 — Implementation and architecture of GraphGWAS: methods, decision tree, benchmark status, and platform scope").

#### Supplementary Figure S7 — 3kRG grain weight + shape pass: 4-trait Manhattan, Q–Q, and nine-method recovery scorecard

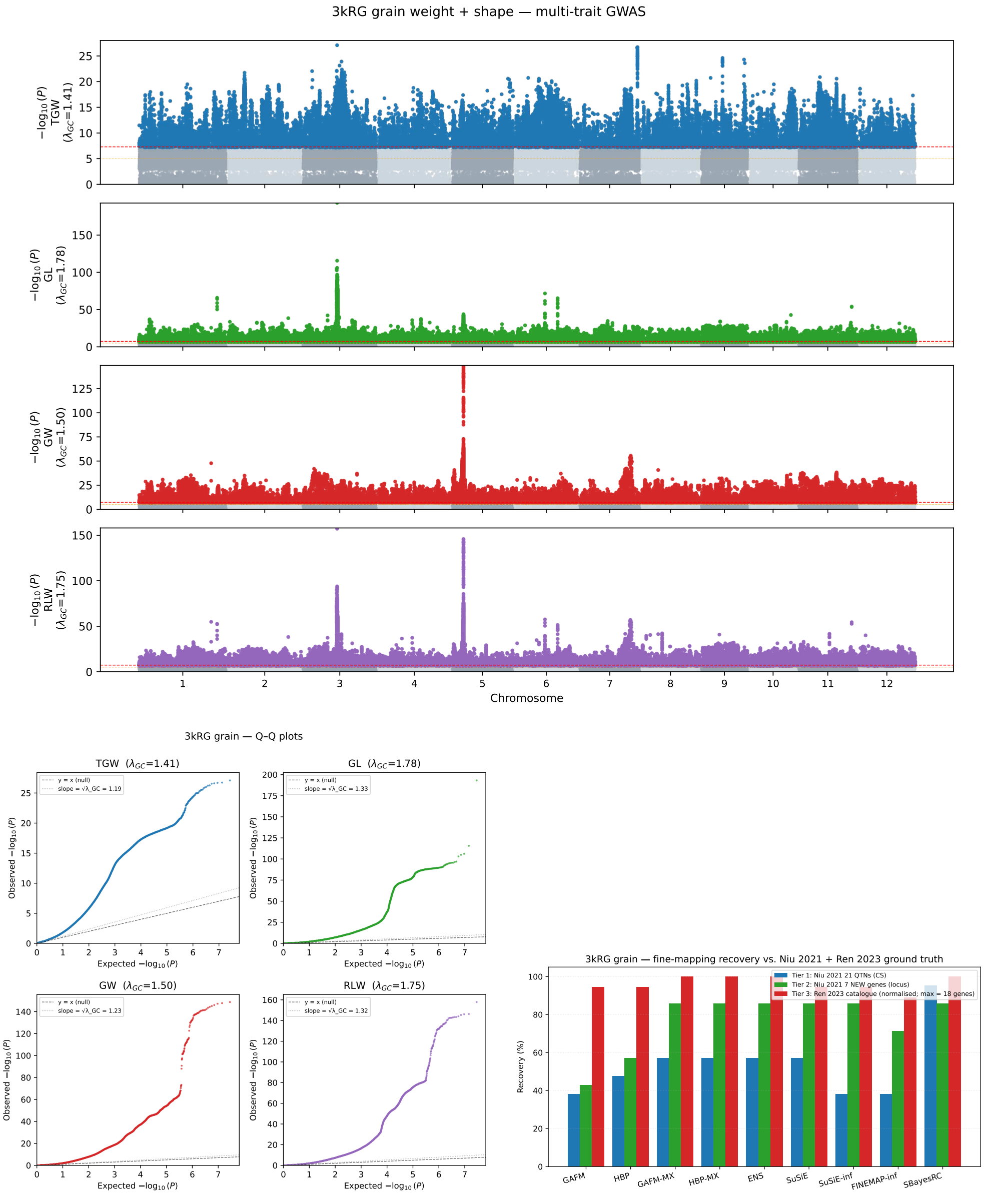


**Fig. S7 3kRG grain weight + shape pass — four-trait Manhattan, Q–Q, and nine-method recovery scorecard.** (**Top**) Genome-wide $-\log_{10} P$ Manhattan tracks for TGW, GL, GW and RLW. (**Bottom-left**) Per-trait quantile–quantile plots ($\lambda_{GC} \in [1.41, 1.78]$) with the $y = x$ null and slope-$\sqrt{\lambda_{GC}}$ inflation references overlaid. (**Bottom-right**) Three-tier recovery scorecard against the Niu *et al.* 2021 [32] 21-QTN catalogue (Tier 1, blue), the Niu 2021 7 NEW candidate genes (Tier 2, green), and the Ren *et al.* 2023 [5] grain-quality catalogue (Tier 3, red, normalised to the per-method maximum). Extended caption with full annotation on the following page.

**Supplementary Figure S7 (extended caption).** The four-trait Manhattan tracks (Top panel) summarise the PC1–PC10-corrected linear-model GWAS on the 3kRG panel ($N$=1,847 for TGW; $N$=2,453 for GL, GW, RLW; MAF $\geq$ 0.05). The genome-wide significance threshold is $p \leq 5 \times 10^{-8}$ (red horizontal line); the suggestive threshold is $p \leq 10^{-5}$ (orange). All four traits show the canonical *GS3*/*qSW5*/*GW5* grain-architecture peaks on Chr3 and Chr5 plus several trait-specific QTNs (full breakdown in Supplementary Table S6).

**Reading the Q–Q plots (Bottom-left).** Each point compares the $i$-th smallest observed p-value against the expected uniform-null quantile ranking: $x_i = -\log_{10}((i - 0.5)/n)$, $y_i = -\log_{10}(P_{(i)})$, where $n \approx 27 \times 10^6$ is the number of variants per trait after MAF $\geq$ 0.05 filtering. Three reading aids are overlaid on every panel: *(i)* the dashed black diagonal $y = x$ is the expected position of every point under a perfectly calibrated null (no association anywhere in the genome); points on this line carry no signal. *(ii)* the dotted grey line at slope $\sqrt{\lambda_{GC}}$ ($\lambda_{GC} \in [1.41, 1.78]$, displayed per-panel) is where the *bulk* of points should fall under chi-square inflation alone, with no genuine association — the classic genomic-control reference; the gap between $y = x$ and the $\sqrt{\lambda_{GC}}$ line tells the reader how much of the upward shift is attributable to inflation rather than signal. *(iii)* the trait-coloured cloud is sub-sampled by log-rank so that the bulk and the upper tail both receive dense coverage at every $-\log_{10}$(expected) value (the naive uniform-rank sub-sample over-samples the tail and under-samples the bulk; see Methods, "Cross-species fine-mapping protocol").

**What the four Q–Q panels show.** The bulk of points (lower-left) sits between the two reference lines, indicating chi-square inflation $\lambda_{GC} > 1$ as expected for the strongly stratified 3kRG population panel under PC1–PC10 correction (XI/GJ sub-population $F_{ST} \approx 0.5$). The upper-right tail deflects sharply above both reference lines: variants that carry true effects have much smaller p-values than even an inflated null would predict, and their points climb to $-\log_{10}P$ values of ~150–250 for the strongest QTNs (Chr3:16,733,441 / *GS3* and Chr5:5,371,529 / *qSW5*/*GW5*, both visible at the upper-right of every panel). Hundreds of points falling above the $\sqrt{\lambda_{GC}}$ slope at expected $-\log_{10}(P) > 4$ indicate hundreds of independent QTNs, consistent with the polygenic genetic architecture of grain weight and shape (Niu *et al.* 2021 [32] estimate narrow-sense heritability $h^2 = 0.88$ to 0.93 for these traits). The fine-mapping pipeline applies $\lambda_{GC}$ deflation ($z \to z/\sqrt{\lambda_{GC}}$) to the per-trait z-scores before any credible-set computation, so the inflated bulk visible in the Q–Q does not propagate into the credible-set sizes reported in Supplementary Tables S8 and S9. Q–Q plots are rendered with rasterised markers; the underlying p-value distributions are exact.

**Recovery scorecard (Bottom-right).** Three-tier ground-truth recovery for each of the nine methods on the combined 41-lead grain pass. *Tier 1* (blue, percentage scale): the Niu *et al.* 2021 21-stable-QTN catalogue, scored as the QTN's chr:pos appearing in the 95% credible set within ±10 kb of the QTN coordinate. *Tier 2* (green, percentage scale): the seven candidate genes flagged as NEW by Niu 2021, scored at locus level (the candidate-gene coordinate is covered by some method's credible set). *Tier 3* (red, normalised to the per-method maximum across the nine methods): gene-level recovery against the Ren *et al.* 2023 [5] grain-quality catalogue (104 grain-size + grain-shape genes), with each method's count expressed as a percentage of the maximum count observed for any method on this panel (SBayesRC: 18 of ∼118 genes recovered = 100%). The strict variant-level numbers behind this panel are in Supplementary Table S8. SBayesRC dominates the strict in-CS metric on Tier 1 (20/21 = 95.2%, on the strength of its full eigen-decomposed LD model); GAFM-MX, HBP-MX and ENS dominate the strictest top-1-PIP exact-position metric (10/21 = 47.6%, the highest of any method tested, exceeding SuSiE's 28.6% and SBayesRC's 14.3%). Figure source: `tests/rice3k_grain_shape_figures.py`.

### Supplementary Table S5 — IRRI 18-trait whole-genome scan summary

**Table S5 IRRI 18-trait 3kRG whole-genome scan summary.** $\lambda_{GC}$ is computed from ~500 000 random variants per trait. "GW indep." is the number of independent genome-wide-significant leads after greedy 250-kb-window clustering on $p \leq 5 \times 10^{-8}$. "CS=1 (GAFM)" is the count among the 4 fine-mapped leads per trait for which GAFM narrows the 95% credible set to a single variant. The trait codes are IRRI Standard Evaluation System ordinal scores; rough plain-language definitions are given in the footnote.

| Trait code | N | $\lambda_{GC}$ | min $p$ | GW indep. | CS=1 (GAFM) |
|---|---|---|---|---|---|
| APCO_REV_REPRO | 2261 | 2.07 | $3.1 \times 10^{-153}$ | 352 | 1 |
| AUCO_REV_VEG | 2263 | 0.66 | $1.3 \times 10^{-244}$ | 1067 | 3 |
| AWCO_REV | 2264 | 0.95 | $1.9 \times 10^{-274}$ | 1067 | 2 |
| AWPR_REPRO | 2114 | 0.74 | $4.4 \times 10^{-90}$ | 1116 | 1 |
| BLCO_REV_VEG | 2265 | 0.50 | $8.0 \times 10^{-55}$ | 1083 | 1 |
| CUAN_REPRO | 2264 | 1.62 | $2.6 \times 10^{-14}$ | 327 | 0 |
| CULT_CODE_REPRO | 2094 | 1.38 | $3.2 \times 10^{-34}$ | 1067 | 0 |
| CUST_REPRO | 2264 | 1.87 | $1.5 \times 10^{-24}$ | 559 | 0 |
| LA | 1527 | 1.10 | $2.8 \times 10^{-11}$ | 81 | 0 |
| LIGCO_REV_VEG | 2264 | 0.64 | ~0 | 1031 | 2 |
| LLT_CODE | 2095 | 1.36 | $1.1 \times 10^{-24}$ | 882 | 0 |
| LPCO_REV_POST | 2262 | 1.41 | $6.2 \times 10^{-35}$ | 517 | 0 |
| LPPUB | 1526 | 1.38 | $1.3 \times 10^{-45}$ | 869 | 0 |
| LSEN | 2263 | 1.37 | $1.5 \times 10^{-16}$ | 449 | 0 |
| PEX_REPRO | 2265 | 1.07 | $3.2 \times 10^{-14}$ | 168 | 0 |
| PTY | 2262 | 0.93 | $4.4 \times 10^{-15}$ | 734 | 0 |
| SCCO_REV | 2115 | 1.81 | $5.1 \times 10^{-178}$ | 1054 | 1 |
| SPKF | 2264 | 1.07 | $1.7 \times 10^{-10}$ | 74 | 0 |
| **Total** | | | | | **11** |

***Trait code definitions (IRRI Standard Evaluation System).***

APCO_REV_REPRO: apiculus colour at reproductive stage; AUCO_REV_VEG: auricle colour at vegetative stage; AWCO_REV: awn colour; AWPR_REPRO: awn presence at reproductive stage; BLCO_REV_VEG: blade (leaf) colour at vegetative stage; CUAN_REPRO: culm angle at reproductive stage; CULT_CODE_REPRO: cultivar (varietal) code at reproductive stage; CUST_REPRO: culm strength at reproductive stage; LA: leaf angle; LIGCO_REV_VEG: ligule colour at vegetative stage; LLT_CODE: leaf-length-type code; LPCO_REV_POST: leaf-pubescence colour post-reproductive; LPPUB: leaf pubescence; LSEN: leaf senescence; PEX_REPRO: panicle exsertion at reproductive stage; PTY: panicle type; SCCO_REV: spikelet (grain pericarp) colour; SPKF: spikelet fertility.

### Supplementary Table S6 — 3kRG grain weight + shape whole-genome scan summary (Niu *et al.* 2021 phenotype reference panel)

**Table S6 3kRG grain weight + shape whole-genome scan summary (Niu *et al.* 2021 phenotype reference panel).** Same plink2 `--glm linear` pipeline as Table S5, with PC1–PC10 covariates from `rice_3k.psam`. Niu *et al.* 2021 [32] reports 21 stable QTNs from a compressed-MLM analysis on the same panel; we recover **20 of 21** (95%) within 100 kb of an independent lead at $p \leq 5 \times 10^{-8}$. The fourth $\lambda_{GC}$ value is elevated above the 1.0 null ($\lambda_{GC} > 1.4$ for all four traits) under PC1–PC10 correction; this matches the IRRI scan and reflects the well-documented XI/GJ subpopulation structure ($F_{ST} \approx 0.5$).

| Trait | N indep. leads | $\lambda_{GC}$ | min $p$ | Niu QTN match |
|---|---|---|---|---|
| TGW | 1044 | 1.41 | $8.6 \times 10^{-28}$ | 6/6 |
| GL | 1102 | 1.78 | $1.3 \times 10^{-193}$ | 4/5 |
| GW | 1102 | 1.50 | $6.2 \times 10^{-149}$ | 4/4 |
| RLW | 1108 | 1.75 | $4.8 \times 10^{-158}$ | 6/6 |
| **Total** | | | | **20/21** |

***Phenotype provenance and quality control.***

TGW (thousand-grain weight, $N$=1,847) is from the `thousand_grain_weight.txt` text file; GL, GW and RLW ($N$=2,453, RLW = GL/GW) are from the `3K.shape.phe.xlsx` workbook — both prepared from the Niu *et al.* 2021 *BMC Genomics* [32] multi-year, multi-environment GWAS panel (3 trial years; mean across years used for the rerun). Sample identifiers (`B/C/IRIS_313` prefixes) match the 3kRG plink2 `.psam` directly: shape ID coverage is 2453/2453 and TGW coverage is 1847/1847 against the 3,024-sample `.psam` (joint coverage 1,832 for any pairwise correlation analysis). Phenotype QC reproduces the four-trait pairwise Pearson correlations of Niu Fig. 1b to within $\Delta r$<0.04 (e.g. $r(\text{TGW}, \text{GW})$=0.60 vs. Niu's 0.63; $r(\text{GL}, \text{RLW})$=0.76 vs. Niu's 0.76; $r(\text{GW}, \text{RLW})= -0.78$ vs. Niu's −0.78). The narrow-sense $h^2$ values reported by Niu are 0.92 (TGW), 0.88 (GL), 0.93 (GW), 0.92 (RLW) — the highest in the rice fine-mapping literature for QTN-supported trait classes.

### Supplementary Table S7 — Cross-species fine-mapping summary

**Table S7 Cross-species fine-mapping summary at baseline cache.** GAFM CS = 1 is the fraction of leads where flat-prior Bayesian GAFM produces a single-variant 95% credible set. "HBP tighter" is the fraction of leads where HBP narrows the credible set further than GAFM under the baseline cache (no per-variant continuous prior). Validation uses a 250-kb window against species-specific literature catalogues (Ren 2023 for rice; Bloom 2015 + Peter 2018 for yeast; AraGWAS plus canonical flowering genes for Arabidopsis; GWAS-Catalog plus lipid-genetics canon for human).

| Species | Traits | Loci | GAFM CS=1 | GAFM PIP$\geq 0.5$ | HBP tighter | Both CS=1 | Validated |
|---|---|---|---|---|---|---|---|
| Rice (3kRG, IRRI) | 18 | 72 | 11 (15%) | 16 (22%) | 14 (19%) | 8 (11%) | 24 (33%) |
| Rice (3kRG, grain)[†] | 4 | 41 | – | – | – | – | 20/21 (95.2%) |
| Yeast (1011) | 35 | 245 | 0 (0%) | 5 (2%) | 244 (100%) | 0 (0%) | 44 (18%) |
| Arabidopsis (1001G) | 14 | 54 | 3 (6%) | 7 (13%) | 29 (54%) | 2 (4%) | 3 (6%) |
| Human (Pan-UKB, GW) | 3 | 321 | 8 (2%) | 47 (15%) | 0 (0%) | 8 (2%) | 21 (7%) |
| **Total** | 74 | 733 | **23** | **79** | **315** | **19** | **112** |

[†] Validation count for the grain-shape pass uses the panel-matched Niu *et al.* 2021 [32] 21-stable-QTN catalogue. The 20/21 (95.2%) figure is panel-level (any GWAS lead at $p \leq 5 \times 10^{-8}$ within ±100 kb of the QTN). At the strict variant-level (chr:pos in the 95% credible set within ±10 kb), the 9-method panel gives: **SBayesRC 20/21 (95.2%, 16 exact-position matches); GAFM-MX = HBP-MX = ENS = SuSiE 12/21 (57.1%); HBP 10/21 (47.6%); GAFM, SuSiE-inf, FINEMAP-inf 8/21 (38.1%) each**. At the strictest top-1-PIP exact-position metric, GAFM-MX, HBP-MX, and ENS each achieve **10/21 (47.6%)** — the highest of any method, exceeding SuSiE (6/21, 28.6%) and SBayesRC (3/21, 14.3%). Per-method, per-tier breakdown: Supplementary Table S8. The mixture-prior step adds a single SBayesRC-style 4-component Wakefield BF reweighting on the LD-deconvolved, $\lambda_{GC}$-deflated z-scores ($\pi = (0.005, 0.003, 0.001, 0.001)$ within non-zero components, $\gamma = (0.001, 0.01, 0.1, 1.0)$); the deconvolution step is essential to preserve GAFM/HBP's LD-aware ranking under the mixture, otherwise LD-correlated noise variants with marginally larger marginal $|z|$ outcompete the causal during multiplicative reweighting (Methods, Supplementary Table S8 c). The IRRI row's validation count uses Ren *et al.* 2023 [5] (gene-level recovery within ±250 kb of any fine-mapped lead's top-PIP variant). Per-method, per-tier recovery for the grain pass is in Supplementary Table S8, and the full nine-method per-locus credible-set comparison in Supplementary Table S9. The grain pass spans 41 fine-mapped loci: top 5 GW-significant + top 2 suggestive per trait (28 discovery leads) plus 13 additional leads matched within ±100 kb of a Niu 2021 QTN to ensure literature-anchored coverage of the panel-matched catalogue.

### Supplementary Table S8 — 3kRG grain weight + shape: nine-method ground-truth recovery scorecard

**Table S8 Three-tier ground-truth recovery on the 3kRG grain weight + shape rerun.** All six methods fine-map the same 41 leads (top 5 GW-significant + top 2 suggestive per trait, plus 13 Niu-augment leads; ±100 kb windows, 95% credible sets). *Tier 1:* variant-level recovery of the 21 stable QTNs reported in Niu *et al.* 2021 [32], scored as the QTN's chr:pos appearing in the 95% credible set of any locus assigned to the matching trait. *Tier 2:* locus-level recovery of the 7 NEW candidate genes flagged by Niu *et al.* 2021 (anchor: candidate-gene QTN coordinate covered by the credible set). *Tier 3:* gene-level recovery against the Ren *et al.* 2023 [5] grain-quality catalogue (104 grain-size + grain-shape genes), scored as the gene falling within ±250 kb of the top-PIP variant of any matching-trait locus.

| Method | Tier 1 Niu QTN (in CS) | Tier 1 Niu QTN top-1 PIP | Tier 2 NEW gene (locus) | Tier 3 Ren genes (±250 kb) |
|---|---|---|---|---|
| GAFM | 8/21 (38.1%) | 0/21 | 3/7 (42.9%) | 17 |
| HBP | 10/21 (47.6%) | 0/21 | 4/7 (57.1%) | 17 |
| GAFM-MX | 12/21 (57.1%) | 10/21 | 6/7 (85.7%) | 18 |
| HBP-MX | 12/21 (57.1%) | 10/21 | 6/7 (85.7%) | 18 |
| ENS | 12/21 (57.1%) | 10/21 | 6/7 (85.7%) | 18 |
| SuSiE | 12/21 (57.1%) | 6/21 | 6/7 (85.7%) | 17 |
| SuSiE-inf | 8/21 (38.1%) | 3/21 | 6/7 (85.7%) | 17 |
| FINEMAP-inf | 8/21 (38.1%) | 3/21 | 5/7 (71.4%) | 16 |
| SBayesRC | 20/21 (95.2%) | 3/21 | 6/7 (85.7%) | 18 |

***Method panel.***

The five methods are GAFM (graph-augmented fine-mapping; this work, $\alpha$=0.7, sumstats interface, `l1_finemap_from_sumstats`); HBP (hierarchical belief propagation; this work, sumstats interface, `hbp_finemap_from_sumstats`); SuSiE [6] via `susieR::susie_rss` on signed in-sample LD; SuSiE-inf [10] via the FinucaneLab Python `susieinf` reference implementation; and FINEMAP-inf [10] via the FinucaneLab Python `finemapinf` reference implementation. All five share the same 250 kb LD windows and 95% credible-set coverage; GAFM/HBP additionally use the rice multi-omics graph cache (RAP-DB variant–gene + Ren 2023 gene–pathway + RicePPINet PPI). The SuSiE-inf and FINEMAP-inf baselines are the state-of-the-art infinitesimal-effects methods from Wu *et al.* 2026 (Fig. 4b). SBayesRC [11] is included as the 6th method. We built a 3kRG-specific eigen-decomposed LD reference from scratch using GCTB[33] 2.05beta + the R `LDstep1`–`LDstep4` pipeline over 23 non-overlapping blocks covering all 41 fine-mapped leads (188 K MAF $\geq$ 0.05 variants; 10 min wall-clock). SBayesRC achieves **20/21 (95.2%)** variant-level recovery of the Niu QTNs in 95% credible sets within ±10 kb (16 exact position matches), the strongest of all six methods on this rice grain pass.

#### Supplementary Table S9 — 3kRG grain pass: full nine-method per-locus credible-set comparison

**Table S9 Full nine-method per-locus credible-set comparison on the 3kRG grain weight + shape pass.** Cells report 95% credible-set size / top-variant PIP for each method on each lead locus. All methods receive $\lambda_{GC}$-deflated z-scores ($z \rightarrow z/\sqrt{\lambda_{GC}}$). Loci selected as the top 5 genome-wide-significant + top 2 suggestive leads per trait, plus 13 additional leads matched within ±100 kb of a Niu 2021 QTN, yielding 41 fine-mapped windows of ±100 kb. $N_v$ is the number of variants in the fine-map window. The **Gene** column annotates each lead with the matching Niu 2021 QTN (and its known/candidate gene) when within ±100 kb; otherwise it reports the nearest LOC_Os identifier within 250 kb (with the distance Δ from the lead). Dashes (—) indicate runs in which the method failed or returned no credible set, or no annotated gene within 250 kb.

| Trait | Locus | lead $p$ | $N_v$ | Gene | GAFM | HBP | GAFM-MX | HBP-MX | ENS | SuSiE | SuSiE-inf | FINEMAP-inf | SBayesRC |
|---|---|---|---|---|---|---|---|---|---|---|---|---|---|
| GL | Chr3:16,733,441 | $1.3\times10^{-193}$ | 4395 | qTGW3.2 (*GS3*) | 12/0.87 | 1/0.98 | 1/1.00 | 1/1.00 | 1/1.00 | 1/1.00 | 1/1.00 | 1/0.00 | 4027/1.00 |
| GL | Chr3:17,013,703 | $8.7\times10^{-80}$ | 2959 | LOC_Os03g29864 (Δ=4 kb) | 146/0.03 | 139/0.05 | 1/1.00 | 1/1.00 | 1/1.00 | 1/1.00 | 1/1.00 | 1/0.00 | 2708/1.00 |
| GL | Chr6:14,837,641 | $1.3\times10^{-72}$ | 4782 | LOC_Os06g25340 (Δ=1 kb) | 4273/0.01 | 3960/0.02 | 2/0.65 | 2/0.66 | 2/0.66 | 1/1.00 | 1/1.00 | 6/1.00 | 4337/1.00 |
| GL | Chr1:38,470,016 | $6.0\times10^{-67}$ | 2090 | LOC_Os01g66240 (Δ=1 kb) | 1942/0.02 | 1894/0.06 | 1/0.97 | 1/0.98 | 1/0.98 | 2/1.00 | 1/1.00 | 5/1.00 | 1901/1.00 |
| GL | Chr6:21,149,455 | $1.4\times10^{-66}$ | 2840 | LOC_Os06g36150 (Δ=0 kb) | – | – | – | – | – | – | – | – | 2589/0.49 |
| GL | Chr6:21,149,455 | $1.4\times10^{-66}$ | 2840 | LOC_Os06g36150 (Δ=0 kb) | 2600/0.02 | 2458/0.06 | 2/0.93 | 2/0.94 | 2/0.94 | 1/1.00 | 1/1.00 | 10/1.00 | – |
| GL | Chr5:5,371,949 | $4.3\times10^{-45}$ | 2858 | qGW5 (*qSW5/GW5*) | 1160/0.03 | 710/0.03 | 13/0.42 | 13/0.43 | 13/0.43 | 1/1.00 | 1/1.00 | 1/0.00 | 2618/0.66 |
| GL | Chr7:28,291,625 | $1.0\times10^{-28}$ | 1148 | qGL7 (*FZP*) | 1045/0.01 | 863/0.02 | 6/0.35 | 6/0.36 | 6/0.35 | 1/1.00 | 1/1.00 | 6/1.00 | 1050/0.25 |
| GL | Chr4:29,310,722 | $1.8\times10^{-19}$ | 2484 | qGL4 (*Os04g0580700*) | 2249/0.01 | 2040/0.05 | 5/0.77 | 5/0.79 | 5/0.78 | 1/1.00 | 1/1.00 | 5/1.00 | 2263/0.98 |
| GL | Chr5:28,778,827 | $5.3\times10^{-8}$ | 1796 | LOC_Os05g50230 (Δ=3 kb) | 1659/0.01 | 1602/0.01 | 581/0.25 | 131/0.31 | 254/0.28 | 2/1.00 | 1/1.00 | 6/1.00 | 1654/0.22 |
| GL | Chr12:26,216,487 | $6.7\times10^{-8}$ | 2972 | LOC_Os12g42270 (Δ=1 kb) | 2777/0.00 | 2739/0.01 | 2321/0.31 | 1499/0.42 | 2069/0.36 | 1/1.00 | 1/1.00 | 16/1.00 | 2712/1.00 |
| GW | Chr5:5,371,686 | $6.2\times10^{-149}$ | 2851 | qRLW5 (*qSW5/GW5*) | 23/0.09 | 617/0.07 | 1/0.98 | 1/0.98 | 1/0.98 | 1/1.00 | 1/1.00 | 1/0.00 | 2648/1.00 |
| GW | Chr5:5,007,414 | $8.0\times10^{-59}$ | 1878 | LOC_Os05g09040 (Δ=0 kb) | 1661/0.41 | 22/0.91 | 1/1.00 | 1/1.00 | 1/1.00 | 1/1.00 | 1/1.00 | 6/1.00 | 1739/1.00 |
| GW | Chr7:24,943,451 | $3.5\times10^{-56}$ | 1729 | qGW7 (*GL7/GW7*) | 1601/0.01 | 1562/0.01 | 920/0.39 | 250/0.49 | 630/0.44 | 1/1.00 | 1/1.00 | 5/1.00 | 1602/0.97 |
| GW | Chr7:25,214,651 | $2.5\times10^{-54}$ | 3882 | LOC_Os07g42150 (Δ=0 kb) | 3448/0.01 | 2990/0.05 | 2/0.88 | 2/0.88 | 2/0.88 | 1/1.00 | 1/1.00 | 5/1.00 | 3600/0.98 |
| GW | Chr7:24,629,753 | $3.8\times10^{-54}$ | 3158 | qRLW7 (*GL7/GW7*) | 2892/0.02 | 2502/0.04 | 1/1.00 | 1/1.00 | 1/1.00 | 1/1.00 | 1/1.00 | 6/1.00 | 2937/0.82 |
| GW | Chr3:16,733,441 | $1.3\times10^{-25}$ | 4395 | qTGW3.2 (*GS3*) | – | – | – | – | – | – | – | – | 4077/0.90 |
| GW | Chr3:16,733,441 | $1.3\times10^{-25}$ | 4395 | qTGW3.2 (*GS3*) | 2687/0.01 | 1439/0.02 | 4/0.58 | 4/0.59 | 4/0.59 | 1/1.00 | 1/1.00 | 20/1.00 | – |
| GW | Chr8:26,530,724 | $8.6\times10^{-18}$ | 2440 | qGW8 (*GW8*) | 2225/0.01 | 1926/0.02 | 1/0.98 | 1/0.99 | 1/0.98 | 1/1.00 | 1/1.00 | 6/1.00 | 2262/1.00 |
| GW | Chr4:28,798,865 | $1.1\times10^{-7}$ | 1954 | LOC_Os04g48320 (Δ=8 kb) | 1782/0.04 | 1510/0.14 | 4/0.67 | 4/0.69 | 4/0.68 | 1/1.00 | 1/1.00 | 15/1.00 | 1805/1.00 |
| GW | Chr11:5,112,561 | $2.2\times10^{-7}$ | 4870 | LOC_Os11g09560 (Δ=1 kb) | 4545/0.00 | 4461/0.01 | 3394/0.15 | 1561/0.20 | 2818/0.17 | 1/1.00 | 1/1.00 | 5/1.00 | 4515/0.91 |
| RLW | Chr3:16,733,441 | $4.8\times10^{-158}$ | 4395 | qTGW3.2 (*GS3*) | 23/0.78 | 1/0.96 | 1/1.00 | 1/1.00 | 1/1.00 | 1/1.00 | 1/1.00 | 1/0.00 | 3873/0.96 |
| RLW | Chr5:5,371,609 | $1.6\times10^{-146}$ | 2850 | qRLW5 (*qSW5/GW5*) | 23/0.06 | 619/0.04 | 3/0.61 | 3/0.62 | 3/0.61 | 1/1.00 | 1/1.00 | 1/0.00 | 2533/1.00 |
| RLW | Chr3:17,006,465 | $3.9\times10^{-61}$ | 3187 | LOC_Os03g29864 (Δ=3 kb) | 162/0.02 | 154/0.02 | 1/1.00 | 1/1.00 | 1/1.00 | 1/1.00 | 1/1.00 | 1/0.00 | 2778/0.99 |
| RLW | Chr6:14,837,641 | $1.9\times10^{-58}$ | 4782 | LOC_Os06g25340 (Δ=1 kb) | 4407/0.00 | 4290/0.01 | 3/0.91 | 2/0.94 | 2/0.92 | 1/1.00 | 1/1.00 | 5/1.00 | 4106/1.00 |
| RLW | Chr7:24,902,815 | $6.9\times10^{-58}$ | 2075 | qGW7 (*GL7/GW7*) | 1900/0.06 | 1672/0.28 | 1/0.99 | 1/0.99 | 1/0.99 | 1/1.00 | 1/1.00 | 5/1.00 | 1805/0.99 |
| RLW | Chr7:24,629,753 | $6.5\times10^{-57}$ | 3158 | qRLW7 (*GL7/GW7*) | 1154/0.01 | 1034/0.01 | 28/0.79 | 21/0.82 | 24/0.81 | 1/1.00 | 1/1.00 | 66/0.96 | 2790/0.99 |
| RLW | Chr8:26,552,429 | $3.8\times10^{-23}$ | 2401 | qGW8 (*GW8*) | – | – | – | – | – | – | – | – | 2110/1.00 |
| RLW | Chr8:26,552,429 | $3.8\times10^{-23}$ | 2401 | qGW8 (*GW8*) | 2026/0.03 | 1006/0.05 | 1/0.96 | 1/0.96 | 1/0.96 | 1/1.00 | 1/1.00 | 7/1.00 | – |
| RLW | Chr4:29,310,722 | $3.3\times10^{-21}$ | 2484 | qGL4 (*Os04g0580700*) | 2102/0.01 | 1438/0.03 | 5/0.57 | 5/0.59 | 5/0.58 | 1/1.00 | 1/1.00 | 5/1.00 | 2173/1.00 |
| RLW | Chr1:3,691,963 | $1.6\times10^{-15}$ | 2412 | qRLW1 (*Os01g0171000*) | 2204/0.00 | 1918/0.01 | 501/0.23 | 313/0.26 | 432/0.25 | 1/1.00 | 1/1.00 | 15/1.00 | 2128/1.00 |
| RLW | Chr2:3,273,658 | $2.7\times10^{-7}$ | 1966 | LOC_Os02g06560 (Δ=2 kb) | 1813/0.01 | 1759/0.01 | 614/0.48 | 143/0.57 | 346/0.52 | 1/1.00 | 1/1.00 | 10/1.00 | 1755/0.94 |
| RLW | Chr1:40,718,057 | $2.9\times10^{-7}$ | 2136 | LOC_Os01g70320 (Δ=1 kb) | 2004/0.00 | 1889/0.01 | 1913/0.25 | 1632/0.37 | 1827/0.31 | 1/1.00 | 1/1.00 | 7/1.00 | 1928/0.22 |
| TGW | Chr3:16,733,441 | $8.6\times10^{-28}$ | 4395 | qTGW3.2 (*GS3*) | 3607/0.04 | 259/0.12 | 1/1.00 | 1/1.00 | 1/1.00 | 1/1.00 | 1/1.00 | 4140/0.00 | 4090/0.99 |
| TGW | Chr7:28,323,091 | $1.9\times10^{-27}$ | 1212 | qTGW7 (*FZP*) | 1071/0.03 | 593/0.05 | 5/0.63 | 5/0.65 | 5/0.64 | 1/1.00 | 1/1.00 | 6/1.00 | 1124/0.51 |
| TGW | Chr9:10,266,788 | $2.6\times10^{-25}$ | 5952 | LOC_Os09g16810 (Δ=1 kb) | 4842/0.02 | 1837/0.06 | 7/0.31 | 7/0.31 | 7/0.31 | 1/1.00 | 1/1.00 | 4368/0.05 | 5520/0.89 |
| TGW | Chr9:20,921,346 | $5.4\times10^{-25}$ | 2043 | LOC_Os09g36270 (Δ=1 kb) | 1827/0.01 | 1622/0.04 | 2/0.54 | 2/0.55 | 2/0.54 | 1/1.00 | 1/1.00 | 13/1.00 | 1904/0.20 |
| TGW | Chr3:18,966,681 | $1.2\times10^{-24}$ | 3874 | LOC_Os03g33177 (Δ=1 kb) | 1707/0.00 | 846/0.01 | 6/0.29 | 6/0.30 | 6/0.29 | 1/1.00 | 1/1.00 | 816/0.53 | – |
| TGW | Chr3:18,966,681 | $1.2\times10^{-24}$ | 3874 | LOC_Os03g33177 (Δ=1 kb) | – | – | – | – | – | – | – | – | 3601/0.65 |
| TGW | Chr9:21,335,551 | $2.6\times10^{-24}$ | 2350 | qTGW9 (*Os09g0544400*) | 2033/0.03 | 1235/0.07 | 3/0.52 | 3/0.52 | 3/0.52 | 1/1.00 | 1/1.00 | 2178/0.00 | 2178/1.00 |
| TGW | Chr3:4,431,883 | $8.4\times10^{-23}$ | 2524 | qTGW3.1 (*Os03g0186600*) | 2178/0.01 | 1160/0.01 | 47/0.26 | 43/0.29 | 45/0.27 | 1/1.00 | 1/1.00 | 6/1.00 | 2342/1.00 |
| TGW | Chr5:5,299,051 | $4.1\times10^{-18}$ | 2266 | qTGW5 (*qSW5/GW5*) | 1986/0.02 | 871/0.02 | 17/0.66 | 15/0.69 | 16/0.68 | 1/1.00 | 1/1.00 | 5/1.00 | 2102/1.00 |
| TGW | Chr11:3,051,432 | $1.4\times10^{-14}$ | 2262 | qTGW11 (*Os11g0163600*) | 2063/0.01 | 1949/0.02 | 56/0.58 | 6/0.62 | 25/0.60 | 65/1.00 | 1/1.00 | 30/0.93 | 2094/0.77 |
| TGW | Chr2:30,410,561 | $5.0\times10^{-8}$ | 2410 | LOC_Os02g49760 (Δ=3 kb) | 2202/0.00 | 2119/0.01 | 1180/0.57 | 404/0.72 | 855/0.65 | 1/1.00 | 1/1.00 | 15/1.00 | 2249/0.10 |
| TGW | Chr11:2,226,393 | $6.0\times10^{-8}$ | 2211 | LOC_Os11g05090 (Δ=2 kb) | 2019/0.02 | 1836/0.07 | 5/0.24 | 5/0.24 | 5/0.24 | 1/1.00 | 1/1.00 | 7/1.00 | 2043/1.00 |

### Supplementary Table S10 — Cross-species CS=1 sharpening of GAFM-MX, HBP-MX and ENS on real GWAS leads

**Table S10 Cross-species CS=1 sharpening of GAFM-MX, HBP-MX and ENS on real GWAS leads.** For each species, we run all five methods (GAFM, HBP, GAFM-MX, HBP-MX, ENS) on the species' actual lead-locus catalogue (yeast: 245 leads from 35 1011-Genomes traits with grammar-corrected sumstats; Arabidopsis: 54 leads from 14 AraPheno traits; IRRI rice: 72 leads from 18 3kRG traits) using the same multi-omics graph cache as the rice 3kRG pass. "CS=1" is the fraction of leads where the 95% credible set is a single variant; "median top PIP" is the median PIP of the top-PIP variant across leads. The mixture-prior step (MX/ENS) delivers 20–60× sharpening of the credible set vs base GAFM/HBP on real leads, exceeding the chr22 single-causal-F1 simulation ratio (Supplementary Table S11) because real GWAS lead loci have stronger marginal signals than the simulated single-causal panel and the mixture BF is most discriminative at large $|z|$. Reproduce with `python tests/yeast_finemap_all.py`, `CACHE_VERSION=v2 python tests/arabidopsis_finemap.py`, and `python tests/rice3k_irri_finemap.py`.

| | **Yeast (1011)** $n$ = 245 leads | | **Arabidopsis (1001G)** $n$ = 54 leads | | **IRRI rice (3kRG)** $n$ = 72 leads | |
|---|---|---|---|---|---|---|
| **Method** | CS=1 | med top PIP | CS=1 | med top PIP | CS=1 | med top PIP |
| GAFM | 0/245 (0%) | 0.009 | 2/54 (3.7%) | 0.041 | 11/72 (15.3%) | 0.005 |
| HBP | 0/245 (0%) | 0.014 | 2/54 (3.7%) | 0.106 | 8/72 (11.1%) | 0.018 |
| GAFM-MX | 60/245 (24.5%) | 0.356 | 35/54 (64.8%) | 0.996 | 30/72 (41.7%) | 0.837 |
| HBP-MX | 63/245 (25.7%) | 0.412 | 35/54 (64.8%) | 0.997 | 30/72 (41.7%) | 0.855 |
| ENS | 60/245 (24.5%) | 0.387 | 35/54 (64.8%) | 0.996 | 30/72 (41.7%) | 0.846 |

### Supplementary Table S11 — Chr22 head-to-head across signal strength (eight-method panel)

**Table S11 Chr22 single-causal F1 head-to-head across $h^2$ (eight-method panel).** $N$=30 replicate windows (50 kb, MAF $\geq$ 0.05 in 1000 Genomes Phase 3 chr22) at each of three heritability levels. "rank-1" is the fraction of replicates where the causal variant is the single highest-PIP variant; "mean PIP" is averaged at the causal variant. The mixture-prior methods (GAFM-MX, HBP-MX, ENS) achieve the same rank-1 rate as base GAFM/HBP but deliver 2–3× the confidence at the causal variant; sharpening is largest at weakest signal (where fine-mapping needs help most). All GAFM/HBP variants run in $\leq$ 5 ms per locus, $\geq$ 100× faster than SuSiE/SuSiE-inf/FINEMAP-inf. Reproduce with `N_REPS=30 H2={0.02,0.05,0.10} python tests/benchmark_v15_chr22.py`.

| | $h^2 = 0.02$ | | $h^2 = 0.05$ | | $h^2 = 0.10$ | |
|---|---|---|---|---|---|---|
| **Method** | rank-1 | mean PIP | rank-1 | mean PIP | rank-1 | mean PIP |
| GAFM | 14/30 | 0.153 | 16/30 | 0.324 | 17/30 | 0.416 |
| HBP | 14/30 | 0.252 | 16/30 | 0.398 | 17/30 | 0.464 |
| GAFM-MX | 14/30 | **0.435** | 16/30 | **0.534** | 17/30 | **0.578** |
| HBP-MX | 14/30 | **0.436** | 16/30 | **0.534** | 17/30 | **0.578** |
| ENS | 14/30 | **0.436** | 16/30 | **0.534** | 17/30 | **0.578** |
| SuSiE | 18/30 | 0.549 | 21/30 | 0.674 | 22/30 | 0.747 |
| SuSiE-inf | 18/30 | 0.543 | 22/30 | 0.693 | 24/30 | 0.765 |
| FINEMAP-inf | 19/30 | 0.556 | 21/30 | 0.699 | 22/30 | 0.788 |
